# Biohybrid Robots with Embedded Conductive Fibers for Actuation, Sensing, and Closed-loop Control

**DOI:** 10.64898/2026.04.01.715915

**Authors:** Xinran Xie, Yuhui Zhao, Ruiheng Wu, Wenjia Xu, Michael J. Bennington, Rachel Daso, Junyi Liu, Abhijith Surendran, Josiah Hester, Victoria A. Webster-Wood, Tingyu Cheng, Jonathan Rivnay

## Abstract

Living organisms achieve adaptive actuation through the seamless integration of neural motor control circuitry and proprioceptive feedback. While biohybrid robotics aims to replicate these capabilities by merging engineered muscle with synthetic scaffolds, the field remains limited by interfaces that lack the efficiency and closed-loop regulation of natural neuromuscular systems. Here, we introduce a biohybrid muscle actuator system featuring a bioelectronic interface based on soft poly(3,4-ethylenedioxythiophene) (PEDOT) fibers for stimulation and sensing. These fibers conformally couple to muscle tissues, eliciting robust contractions at voltages as low as 1 V—requiring ultra-low power (0.376 ± 0.034 mW) and preserving long-term tissue viability. By leveraging the independent addressability of these fibers, we demonstrate selective actuation of individual muscle units to achieve precise spatiotemporal control of a two-muscle-powered walking biohybrid robot, reaching a locomotion speed of 5.43 ± 0.79 mm/min. When configured as strain sensors, the fibers exhibit a high gauge factor of 155.45 ± 6.59 and resolve contractile displacements within tens of micrometers. We demonstrate that this sensing modality can be integrated into a closed-loop controller to autonomously modulate stimulation based on real-time feedback, significantly mitigating muscle fatigue (p = 0.038) during continuous operation. This work establishes a versatile platform for efficient actuation and intrinsic feedback sensing, providing a blueprint for efficient, autonomous, and adaptive biohybrid machines.

**Summary:** Soft conductive fibers enable a bioelectronic interface for low-power actuation and closed-loop control in biohybrid robots.

## Introduction

Living organisms achieve efficient, adaptive actuation through the tight integration of motor and sensory pathways, enabling robust performance in dynamic environments. Inspired by this biological paradigm, biohybrid robotics couples living muscle tissues with synthetic scaffolds to achieve life-like behaviors, including swimming (*1–5*), crawling or walking (*6–12*), pumping (*13*), and object manipulation (*14–17*). Muscle actuators offer several advantages, including adaptability, high energy conversion efficiency (*18, 19*), and intrinsic self-repair (*20*), that make them well suited for biomedical devices, targeted drug delivery, and autonomous soft robotic platforms. However, the field remains constrained by the lack of stimulation and sensing interfaces that approach the efficiency, selectivity, and closed-loop control of natural neuromuscular systems.

In animals, peripheral motor neurons deliver finely tuned electrical signals that drive muscle contraction through a dedicated neural architecture. Engineered muscle tissues, by contrast, typically rely on external electrical pulses requiring high voltages or invasive contact for effective coupling (*21–23*). Rigid inorganic electrodes interface poorly with soft tissues, and medium-based field stimulation lacks the spatial precision required for complex movements (*10, 24*). Although optogenetics offers high selectivity, it necessitates complex genetic modification and relies on relatively power-intensive light sources (*4, 8, 9, 12, 25, 26*). To address these limitations, a tissue-conformal, direct-contact interface mimicking the localized delivery of biological nerves would enable power-efficient actuation, surpassing the efficiency of current optogenetic methods while allowing high-performance recruitment of individual muscle units.

Yet actuation alone is insufficient for robust motor control. Biological systems rely on continuous sensory feedback to regulate contraction dynamics, maintain stability, and prevent fatigue (*27, 28*). Translating this principle to biohybrid robotics requires integrated sensing modalities capable of reporting muscle deformation in real time (*29*). While prior efforts have used resistive strain sensors (*30–34*) or organic transistor-based devices (*35*), few have achieved closed-loop regulation (*36*), and none have established a consistent correlation between real-time muscle contraction amplitude and sensory output. An interface that realizes efficient stimulation and high-reliability proprioceptive sensing would therefore represent a significant step toward autonomous, adaptive biohybrid robots.

Here, we report a biohybrid robotic system with a bioelectronic interface for low-power, selective actuation and real-time sensory feedback for closed-loop control. Soft PEDOT fibers provide conformal, contact-based stimulation of engineered muscle tissues (Fig. 1Ai–1Aii), eliciting robust contractions at voltages as low as 1 V, and achieving ultra-low-power consumption (0.376 ± 0.034 mW) while preserving tissue viability. Notably, this organic interface elicited greater contractile displacements with reduced power requirements compared to conventional inorganic electrodes or optogenetic systems, highlighting its superior energy efficiency. Directly embedding the bioelectronic interface within the muscle also supported highly specific actuator addressability with minimal crosstalk when multiple actuators were in close proximity. Leveraging the independent addressability of these fibers, we constructed a walking biohybrid robot with two distinct muscle bundles (Fig. 1Aiii), where alternating stimulation enabled precise gait modulation and directional turning, achieving a locomotion speed of 5.43 ± 0.79 mm/min.

**Fig. 1.**
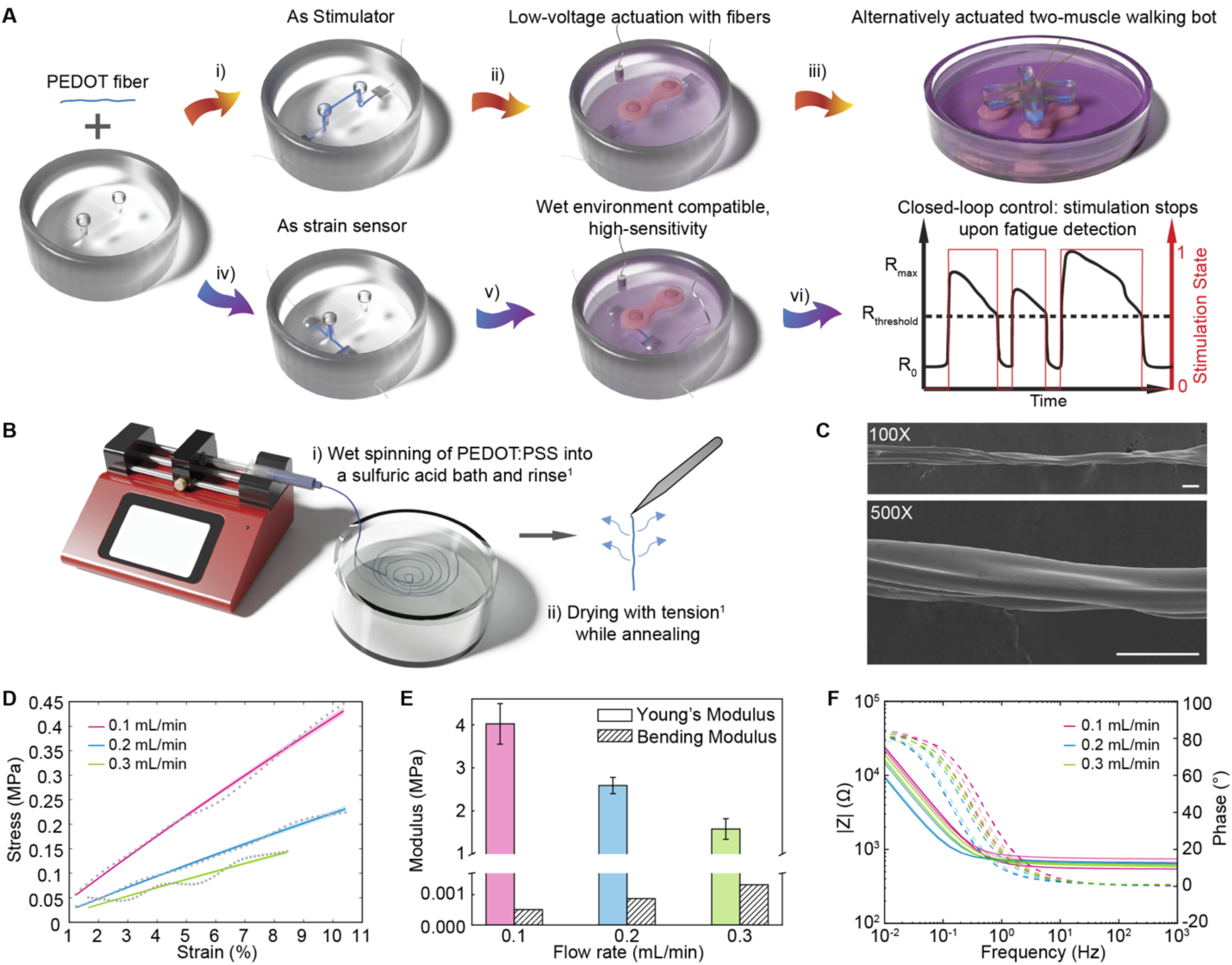
Fabrication and characterization of PEDOT fibers. (**A**) Schematic of the biohybrid robotic system integrating an organic bioelectronic interface for actuation and strain sensing, with fiber placed strategically for different functions. (**B**) Schematic illustration of the wet-spinning fabrication process and subsequent post-processing stages. (**C**) Scanning electron microscopy (SEM) micrographs of post-processed PEDOT fibers. Scale bar, 100 μm. (**D**) Representative tensile stress–strain curves for fibers fabricated at flow rates of 0.1, 0.2, and 0.3 mL/min. (**E**) Mechanical properties (Young’s modulus and bending modulus) as a function of spinning speed. Data are presented as means ± std (N = 3 for 0.1 and 0.3 mL/min; N = 4 for 0.2 mL/mL). (**F**) Bode plots of PEDOT fibers across different spinning speeds (N = 3).

Beyond actuation, the PEDOT fibers demonstrated exceptional sensitivity to physiologically relevant strains. With a gauge factor of 155.45 ± 6.59, two orders of magnitude higher than previously reported muscle strain sensors (*30–34*). The fiber-enabled sensing platform resolved contractile displacements within tens of micrometers with high linearity (R^2^ = 0.891; Fig. 1A, iv and v). We demonstrate that this sensing modality can be integrated into a closed-loop controller to autonomously modulate stimulation based on real-time feedback (Fig. 1Avi), significantly mitigating muscle fatigue compared to a continuous, open-loop stimulation system (p = 0.038).

Together, this work establishes a multimodal (both stimulation and proprioception) interface for integrated biohybrid actuators and multi-actuator biohybrid robots, thereby advancing the development of autonomous, adaptive, and energy-efficient biohybrid machines.

## Results

### PEDOT fiber fabrication and characterization

PEDOT fibers were fabricated by wet spinning PEDOT:PSS solution into a sulfuric acid coagulation bath at flow rates of 0.1, 0.2, or 0.3 mL/min (Fig. 1B). This process promotes polymer crystallization and removes excess PSS, thereby enhancing electrical conductivity (*37, 38*). Subsequent tension-drying and thermal annealing further increased crystallinity, domain alignment, and mechanical robustness (*39*). Scanning electron microscopy (SEM) confirmed that post-treated fibers exhibited smoother surfaces and improved straightness, facilitating reliable integration into the robotic platform (Fig. 1C).

Because these fibers directly contact engineered muscle tissues, mechanical compliance is critical to avoid restricting contractile motion. We therefore evaluated fiber mechanics under physiological conditions in PBS at 37°C (Fig. 1D). Using a Neo-Hookean model, we determined the Young’s modulus for each fiber group and calculated the bending modulus based on fiber geometry (Fig. 1E, fig. S1). Across all spinning conditions, the fiber bending modulus remained within the range of native skeletal muscle (*40*), ensuring mechanical compatibility with dynamic muscle deformation.

Electrical properties were assessed using electrochemical impedance spectroscopy (EIS) (Fig. 1F). We utilized the Randles circuit (fig. S2) to model the electrode-electrolyte interface and determine the double-layer capacitance, which is a key indicator of the fibers’ capacity for safe and effective charge injection. Among the tested flow rates, fibers produced at 0.2 mL/min exhibited the highest capacitance (1414.93 ± 331.77 µF) (fig. S2, table S1), suggesting superior performance for tissue stimulation. Consequently, the 0.2 mL/min fibers were selected for all subsequent device integration.

### Interface design and tissue engineering

To establish a stable bioelectronic interface, the wet-spun fibers were embedded into a polydimethylsiloxane (PDMS) device featuring two pillars designed for muscle growth. The fiber geometry was specifically tailored to its functional role: longitudinal alignment across both pillars for electrical stimulation, or a U-shaped loop within a single pillar for strain sensing (Fig. 1A, i and iv, fig. S3). For the stimulator, the two-pillar structure maintained the fiber under mild tension, with an exposed central segment to ensure direct coupling with the tissue. For the sensor, the fiber was fully encapsulated within the pillar to translate pillar deflection into electrical signals.

C2C12 myoblasts were encapsulated in a Matrigel–fibrin hydrogel and cast in a figure-eight–shaped well (Fig. 2Ai). The two pillars from the devices were then inserted into the cell/hydrogel mixture (Fig. 2Aii, fig. S4). During early proliferation, the confined geometry of the figure-eight well promoted tissue compaction around the pillars (Fig. 2Aiii); subsequent removal of the well allowed for nutrient exchange, muscle maturation, and imaging with an inverted microscope (Fig. 2Aiv, fig. S5). The casting method enables simultaneous muscle maturation and establishment of a robust, conformal fiber–tissue interface.

**Fig. 2.**
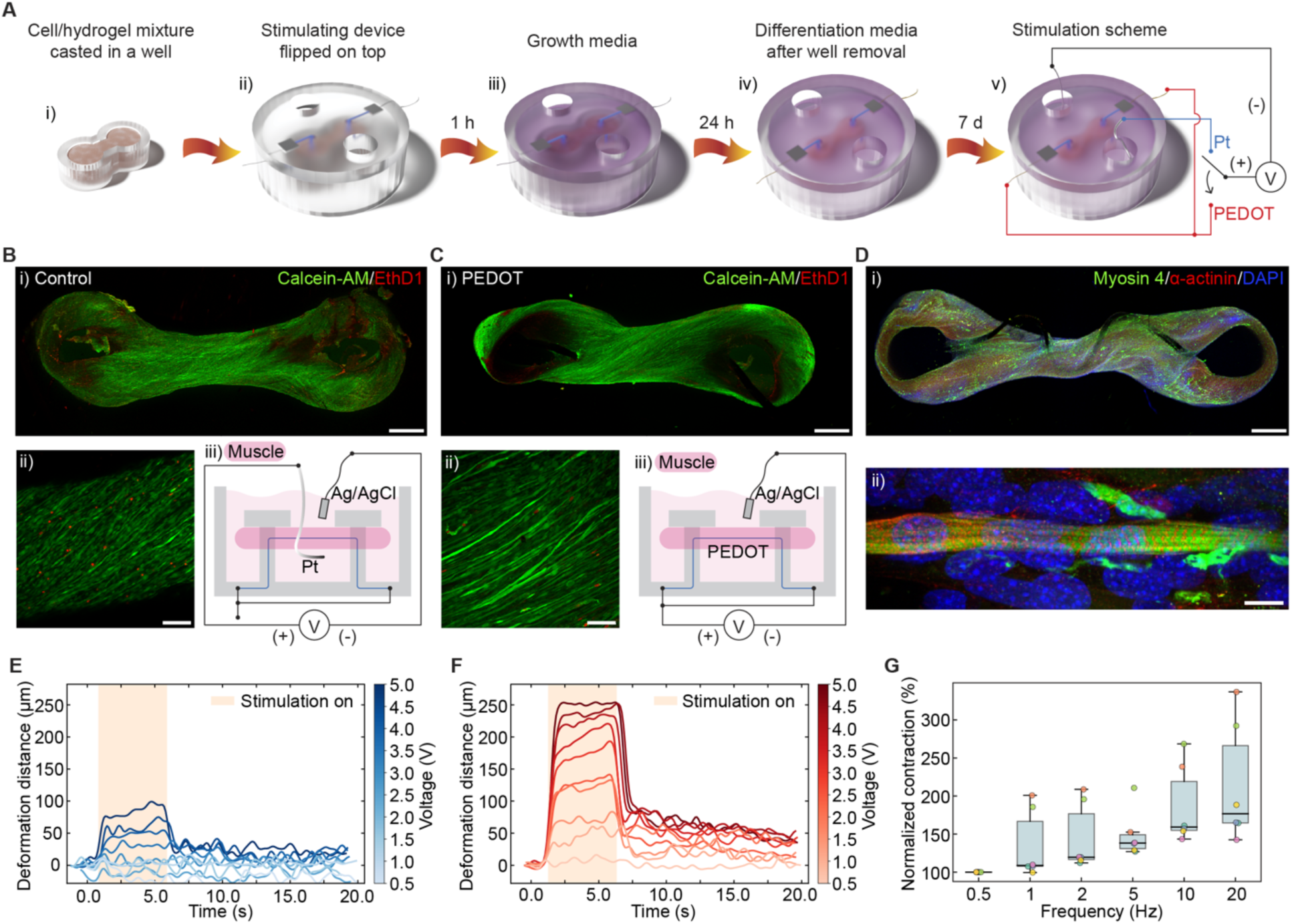
Skeletal muscle differentiation and electro-mechanical coupling via embedded PEDOT fibers. (**A**) Schematic depicting the cell–hydrogel casting process and subsequent skeletal muscle differentiation around a PEDOT fiber–based stimulation device. (**B**, **C**) Live/dead viability assays for (**B**) control muscle tissues and (**C**) PEDOT fiber-embedded muscle tissues. Panels (i) and (ii) show whole-tissue (scale bar, 1 mm) and high-magnification views (scale bar, 100 μm), respectively. Panel (iii) provides a schematic comparison of the external vs. embedded stimulation configurations. (**D**) Immunofluorescence characterization of PEDOT-integrated muscle tissues: (i) whole-tissue view (scale bar, 1 mm); (ii) high-magnification view of sarcomeric organization showing myosin heavy chain (green), sarcomeric-α-actinin (red), and DAPI-stained nuclei (blue). Scale bar, 10 μm. (**E**, **F**) Comparative contraction amplitude as a function of applied stimulation voltage for (**E**) external Pt wires versus (**F**) embedded PEDOT fibers within the same muscle sample. All stimulations were conducted at 20 Hz with a 5 ms pulse width for a 5 s duration. (**G**) Frequency-dependent normalized contraction response using embedded PEDOT fibers. Distinct colors represent individual muscle replicates, with data presented as medians and interquartile ranges (IQR) (N = 6). All stimulations were conducted at 5 V with a 5 ms pulse width.

Live/dead assays confirmed that muscle growth around the fibers did not induce cytotoxicity (Fig. 2, Bi, Bii, Ci, and Cii, fig. S6). After two weeks of differentiation, immunofluorescence staining for myosin heavy chain (Myosin 4) and sarcomeric α-actinin revealed sarcomeric structures, confirming the development of mature myofibers (Fig. 2D, fig. S7). Notably, early-phase myotubes aligned along the fiber axis, indicating stable mechanical coupling and tissue–fiber interface (fig. S8).

### Low-voltage muscle actuation via embedded fiber interfaces

To evaluate stimulation performance, we compared muscle activation outcomes using either the tissue-embedded fibers or conventional platinum (Pt) wire electrodes submerged in the culture medium, using a shared Ag/AgCl counter-reference. Cross-sectional schematics (Fig. 2, Biii and Ciii) illustrate the distinct current pathways: the fiber interface enables direct charge injection into the tissue, whereas the Pt electrode relies on field stimulation through the resistive culture medium. To eliminate inter-sample variability, comparisons were performed on the same muscle unit by switching between the two electrode types (Fig. 2Av).

Relative to external Pt wire stimulation, the embedded fibers substantially reduced the activation threshold and elicited significantly stronger contractions across the same voltage range (Fig. 2, E and F, fig. S9). The embedded interface reliably triggered robust contractions at voltages as low as 1 V (Fig. 2F). To decouple the material advantages of PEDOT from the effects of electrode placement, we developed a four-post validation platform where one fiber was embedded within the muscle while an identical second fiber remained in the surrounding medium (fig. S10). Activation studies on the same muscle confirmed that, at identical voltages, the embedded fiber consistently achieved larger contractile displacements (fig. S11). This finding confirms that the enhanced actuation is primarily due to the localized, conformal electrode placement.

We further quantified stimulation efficiency by calculating the power consumption of both configurations (fig. S12, table S2). An average power of only 0.376 ± 0.034 mW via the fiber interface was sufficient to elicit tetanic muscle contraction (at 1 V). This power requirement is three orders of magnitude lower than a typical commercial stimulator (∼1.3 W) designed for muscle activation (*41*). Across all tested voltages, embedded stimulation consumed equal or less power while triggering larger contractile output compared to the Pt electrode. Notably, even at the highest tested voltage (5 V), the power consumption remained below 10 mW, which is lower than the power requirement for optogenetic actuation (∼2.0 mW/mm^2^ or more) (*12, 25, 26*). These results underscore the system’s ultra-low power consumption and superior energy efficiency.

Leveraging this high efficiency, we characterized how stimulation frequency modulates contractile output. With the pulse width fixed at 5 ms and voltage at 5 V, increasing the frequency yielded a progressive increase in contraction amplitude up to 20 Hz, which produced stable tetanic contractions (Fig. 2G, fig. S13). Consequently, 20 Hz was selected as the standard frequency for sustained, low-voltage muscle activation in subsequent muscle actuator trials.

### Selective muscle activation and robotic locomotion

Building on the enhanced efficiency of embedded stimulation, we investigated whether these fibers could selectively activate individual muscles within a multi-muscle system. We designed a four-post scaffold architecture integrated with two parallel fiber interfaces (Fig. 3Ai, fig. S14). Using a modified casting protocol (Fig. 3Aii, fig. S15), we generated a biohybrid walking robot (biobot) consisting of two parallel muscle bundles, each embedded with a single, independently addressable stimulation fiber (Fig. 3Aiii).

**Fig. 3.**
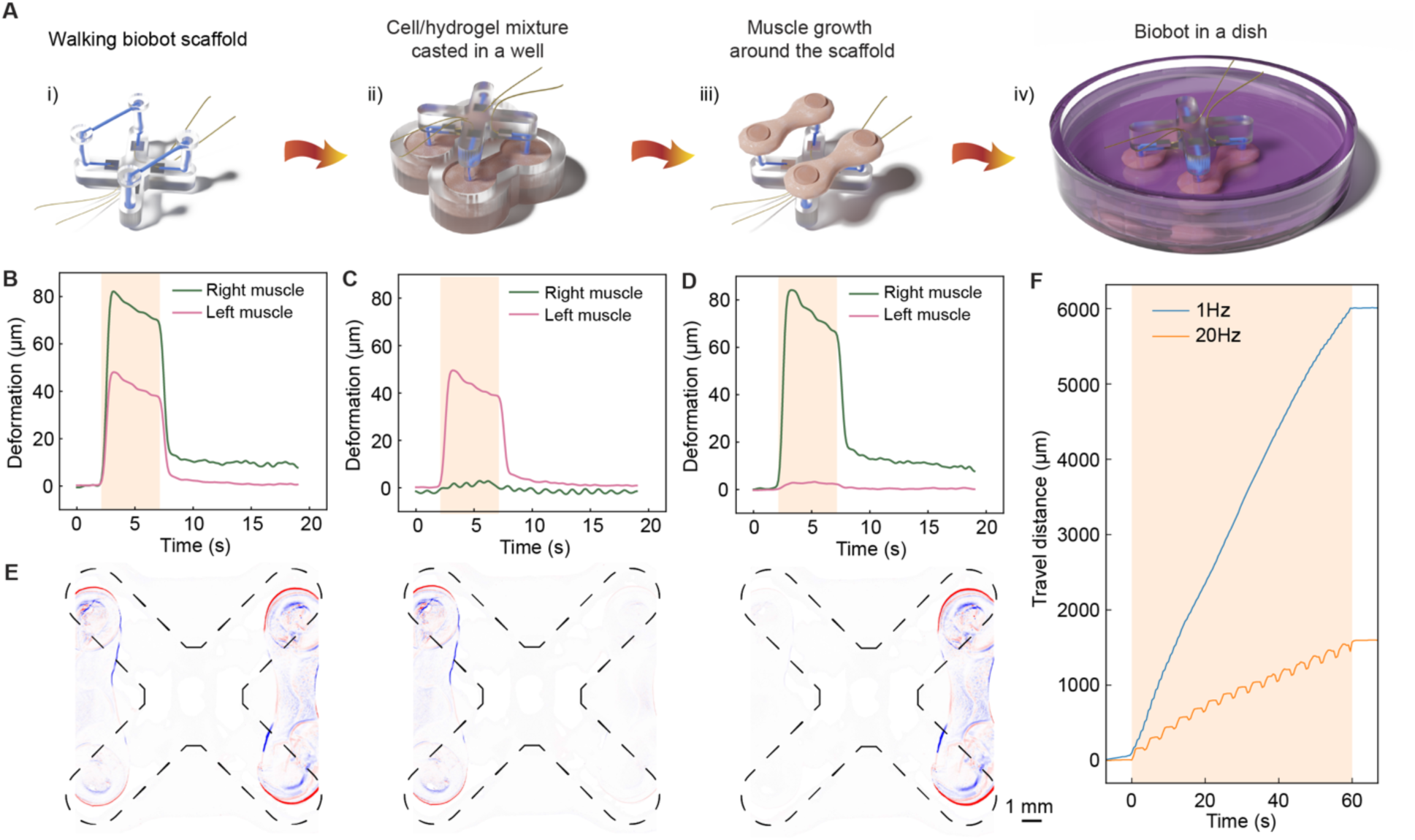
Spatiotemporal control and selective activation of biohybrid robotic systems. (**A**) Schematic of the biohybrid robot fabrication process, featuring independent PEDOT fiber integration into dual-muscle units. (**B**-**D**) Selective contraction profiles of left and right muscles under different stimulation regimes: (**B**) simultaneous activation via both fibers, (**C**) isolated stimulation via the left fiber, and (**D**) isolated stimulation via the right fiber. All stimulations were conducted at 5 V, 20 Hz, with a 5 ms pulse width for a 5 s duration. (**E**) Representative pixel intensity difference maps showing displacement at peak contraction relative to the resting state. Color-coded heatmaps indicate localized tissue movement (red: increased intensity; blue: decreased intensity). Scale bar, 1 mm. (**F**) Representative travel-distance-over-time of the biobot undergoing alternating locomotory gates at stimulation frequencies of 1 Hz and 20 Hz for 60 s.

Selective actuation was achieved by delivering electrical pulses through only one embedded fiber at a time. To quantify the no-load contractile displacement of individual muscles, we utilized a custom handle to suspend the biobot, preventing its contact with the plate bottom (fig. S16). We measured deformation under three distinct conditions: simultaneous bilateral stimulation, left-muscle-only, and right-muscle-only activation (Fig. 3B–D, fig. S17).

To spatially resolve the localized contraction, we generated pixel intensity difference maps relative to the initial frame. Targeted contractile motion was confirmed by localized increases in pixel intensity on the outer edges of the muscle bundles (red) and decreases on the inner edges (blue) (Fig. 3E). While these results confirmed targeted activation, partial co-activation of the non-stimulated muscle was observed in some constructs (fig. S18), likely reflecting variability in muscle maturity or across-tissue excitation thresholds.

By alternating stimulation between the two muscle bundles at a waveform frequency of 1 Hz, the biobot achieved steady directional locomotion with a walking speed of 5.43 ± 0.79 mm/min, maintaining consistent performance over a 60-second stimulation period (Fig. 3F, fig. S19). Analysis of different stimulation protocols (fig. S20) showed that the 1 Hz waveform produced a higher absolute walking speed than the 20 Hz waveform (1.37 ± 0.20 mm/min) (fig. S21). This discrepancy is attributed to hardware-constrained differences in the programmed pacing frequency (the rate of alternation between muscles), which was 2 Hz for the 1 Hz waveform and 0.5 Hz for the 20 Hz waveform. When normalized for pacing frequency, the 20 Hz waveform resulted in a comparable net walking speed to the 1 Hz waveform.

The platform also demonstrated the ability to execute turning maneuvers through unilateral stimulation (fig. S22). By actuating only the left or right muscle, the biobot’s travel trajectory exhibits a 90° directional shift. In the representative biobot shown, the right muscle bundle dominated the locomotion direction. Collectively, these findings demonstrate that embedded fibers provide the selective, independently addressable control required for complex gait modulation and steering in multi-muscle biohybrid robots.

### Fabrication and performance of fiber-based strain sensors

Post-processed PEDOT fibers were first evaluated as standalone strain sensors by applying cyclic tensile strains (0.1% to 0.5%) while monitoring real-time resistance changes. The relative resistance change (ΔR/R) scaled proportionally with strain amplitude and exhibited stability over repeated cycles, demonstrating mechanical durability under physiologically relevant strains (Fig. 4A, fig. S23). From the peak ΔR/R values, we calculated a gauge factor of 155.45 ± 6.59 (R^²^ = 0.9206), indicating high sensitivity and linear response (Fig. 4B). These results motivated the integration of the fibers into a single-post configuration for intrinsic muscle strain sensing (Fig. 1Aiv, fig. S3 and S24).

**Fig. 4.**
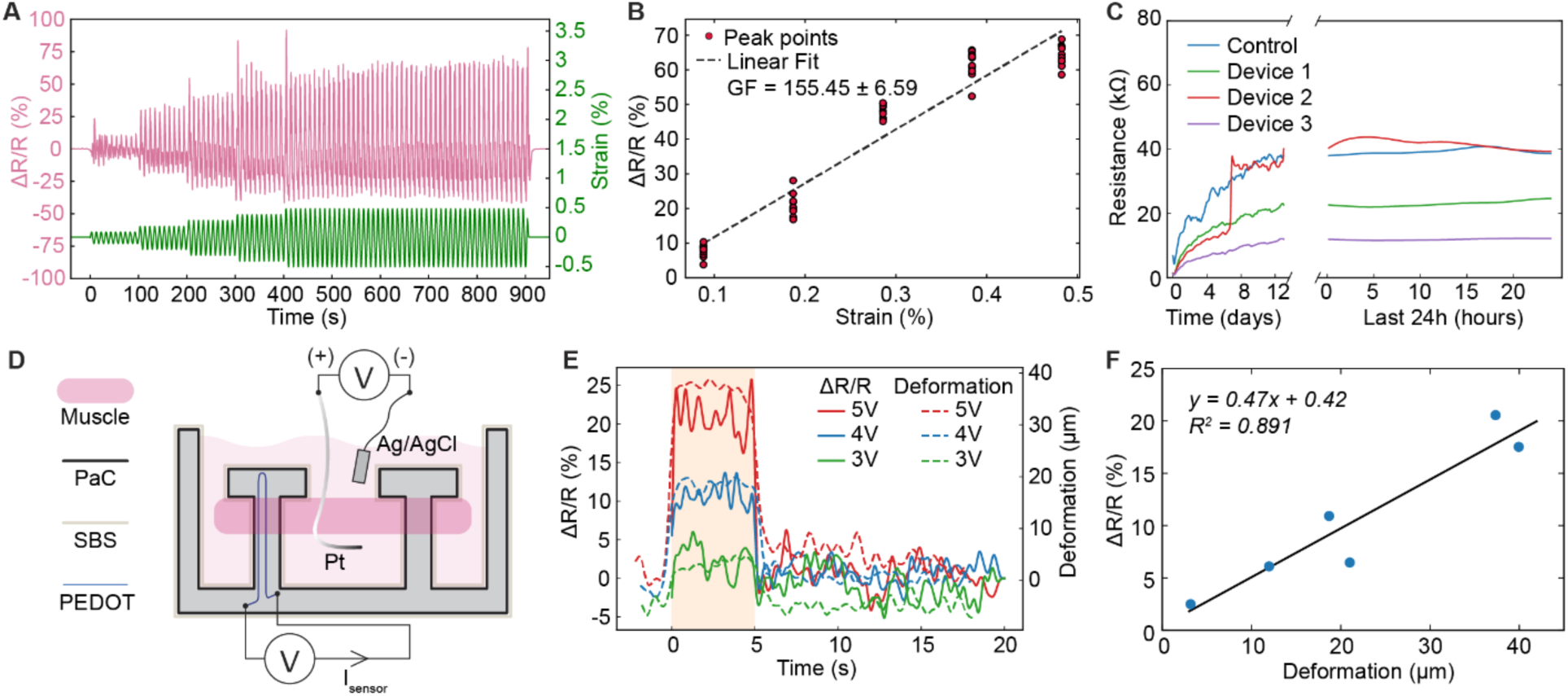
Strain sensing and real-time monitoring of muscle actuation. (**A**) Relative resistance change (ΔR/R, pink) of PEDOT fibers subjected to cyclic sinusoidal loading at strains ranging from 0.1% to 0.5% (green). (**B**) Sensitivity analysis showing the gauge factor (GF) derived from the linear regression of ΔR/R versus applied strain. (**C**) Long-term electrochemical stability comparison between uncoated (control) and encapsulated (coated, N = 3) sensing devices during continuous immersion in PBS. The abrupt increase in resistance for Device 2 around Day 7 implies the onset of encapsulation failure. (**D**) Schematic illustration of the experimental configuration for in situ strain sensing, featuring a PEDOT fiber integrated within a flexible PDMS pillar to monitor muscle-generated deformations. (**E**) Synchronized time-resolved profiles showing the fiber resistance change (ΔR/R, solid lines) and the corresponding muscle contraction amplitude (dashed lines) quantified via optical image analysis. All stimulations were performed at 20 Hz with a 5 ms pulse width for 5 s. (**F**) Linear correlation between the fiber’s electrical response (ΔR/R) and the optically measured muscle contraction amplitude, demonstrating the fiber’s utility as an internal displacement sensor.

To ensure the long-term reliability of these sensors in aqueous culture environments, we developed a multilayer encapsulation strategy (fig. S25). Devices were first coated with parylene-C to minimize water permeation and prevent electrical cross-talk with the stimulation signals conducted through the electrolyte. A secondary, spray-deposited styrene–butadiene–styrene (SBS) elastomer coating was applied to enhance hydrophobicity and promote cell adhesion. Comparative analysis of various elastomer formulations identified SBS to be the most effective in protecting sensor signals from stimulation artifacts (fig. S26).

Encapsulated devices exhibited long-term stability after two weeks of continuous immersion in PBS. Real-time resistance monitoring revealed that sensor resistances converged to steady baselines following the initial 10-day immersion period, a timeline that aligns with muscle maturation and defines the onset of experimental testing (Fig. 4C, fig. S27). To further validate the efficacy of the encapsulation, we monitored sensor signals during electrical stimulation in the absence of muscle tissue after long-term immersion in physiological conditions. While uncoated devices exhibited significant signal artifacts due to ionic interference, the encapsulated sensors remained stable (fig. S28), confirming that the protective coatings successfully isolated the sensing modality from the stimulation field.

Despite the challenges of maintaining sensor performance during multi-week tissue maturation, our integrated platform successfully resolved active contractile dynamics in real time. A cross-sectional schematic (Fig. 4D) illustrates the device with the fiber residing within a single PDMS post to translate mechanical pillar deflection from muscle into electrical resistance changes. The sensor reliably captured the dynamics of tetanic contractions across varying amplitudes (Fig. 4E), as evidenced by the high consistency between the sensor signal and the contractile displacements monitored via optical microscopy. Furthermore, a strong linear correlation between the relative change in resistance (ΔR/R) and contractile displacement (R^2^ = 0.891) was established, achieving a detection resolution on the order of tens of micrometers (Fig. 4F). Although encapsulation reduced the absolute deformation transferred to the fiber, finite element simulations suggested that muscle force output remained comparable between coated and uncoated conditions (fig. S29).

### Closed-loop feedback control via fatigue mitigation

Biohybrid robots are inherently limited by short operational lifespans and performance decay caused by continuous actuation and aggregated muscular fatigue (*42, 43*). To overcome these challenges, we developed a closed-loop control approach designed to enhance functional longevity through adaptive stimulation. We integrated the fiber–based strain sensor with an Arduino microcontroller capable of real-time signal processing and autonomous decision-making. The controller continuously monitored resistance changes corresponding to muscle contraction and dynamically modulated stimulation parameters to mitigate fatigue.

A dual-channel circuit architecture enabled concurrent electrical stimulation and strain sensing (Fig. 5A). To ensure high signal fidelity and eliminate electrochemical artifacts, we implemented a temporal gating strategy: sensor output is sampled exclusively during the quiescent intervals between active stimulation pulses.

**Fig. 5.**
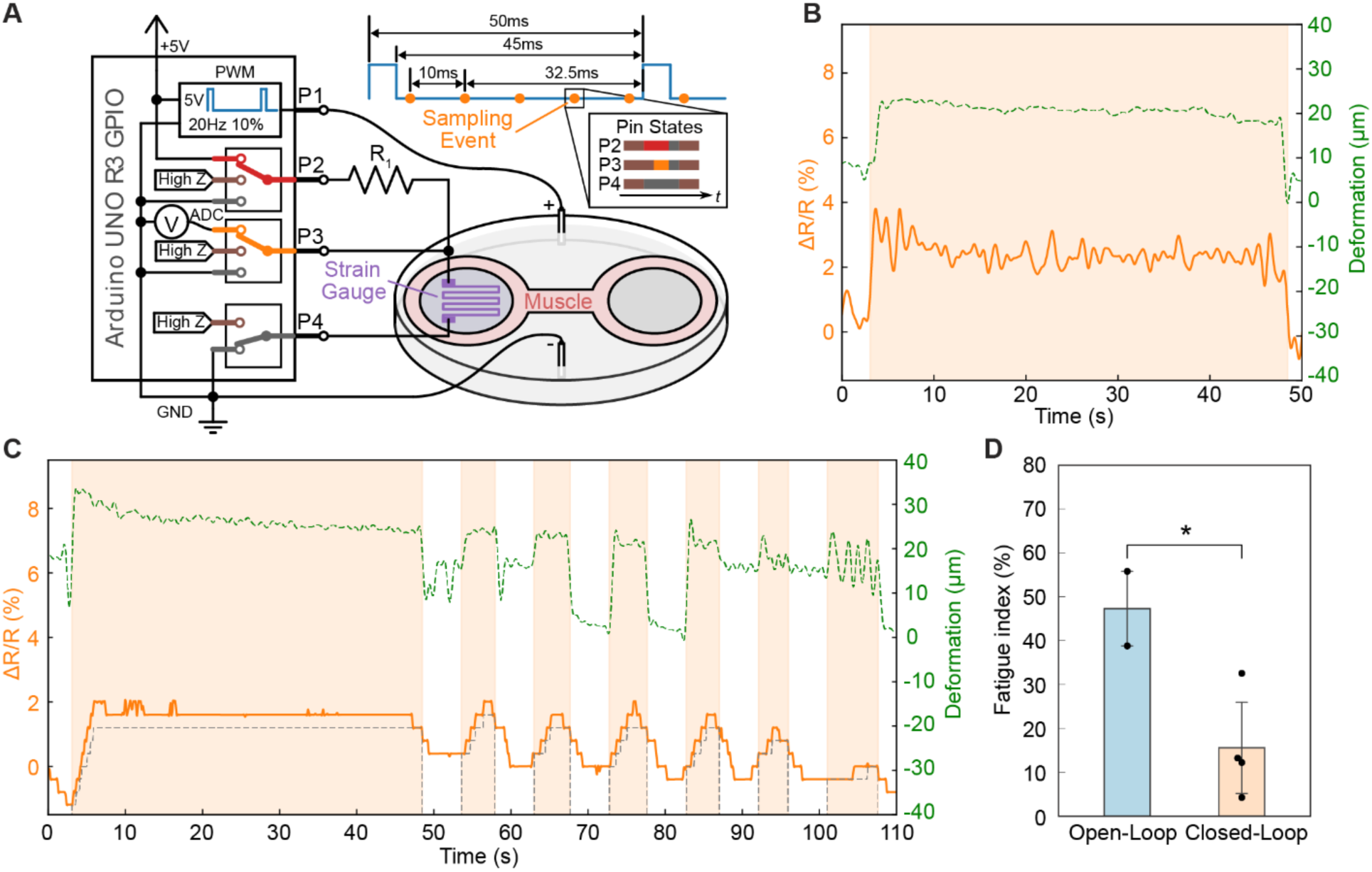
Proprioceptive feedback and autonomous closed-loop control of muscle actuation. (**A**) Schematic of the experimental architecture for Arduino-based closed-loop control, integrating real-time PEDOT fiber strain sensing with adaptive muscle stimulation. The sensor signal is sampled during the inter-pulse intervals of the active stimulation to minimize electrical interference. (**B**) In situ detection of muscle fatigue, characterized by a progressive attenuation in the relative resistance change (ΔR/R, orange solid line) and contractile displacement (green dashed line) during sustained repetitive stimulation. (**C**) Representative performance of the closed-loop control system driven by sensor feedback. Light-orange shaded regions denote active stimulation periods, while the gray dashed line represents the dynamic fatigue threshold (set at R_25%_). (**D**) Statistical comparison of the fatigue index between open-loop (N = 2) and closed-loop (N = 4) control regimes, showing a significant improvement in performance stability under feedback (p = 0.038, two-tailed Student’s t-test). All stimulations were conducted at 5 V, 20 Hz, with a 5 ms pulse width.

Control parameters were established by characterizing baseline resistance during resting states (𝑅_!_) and initial, non-fatigued contractions (𝑅_max_). Upon tetanic stimulation, the resistance increased sharply, reflecting maximum contractile displacement (Fig. 1Avi, Fig. 5B). During sustained tetanus, the resistance signal exhibited a characteristic decay from 𝑅_max_ as fatigue developed, correlating with the progressive decrease in contractile displacement observed via optical microscope (Fig. 5B).

To operationalize this physiological response, we implemented a dynamic fatigue threshold (R_x%_), defined as a percentage drop (x%) relative to the peak contractile range:

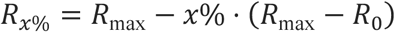

The algorithm evaluates the running average of the resistance (𝑅*_avr_*) to 𝑅_x%_ to determine the optimal termination point for stimulation (when 𝑅*_avr_* < 𝑅_x%_). This smoothing technique prevents stochastic noise or outliers from triggering premature shutoff. In a representative demonstration (Fig. 5C), stimulation was automatically terminated once the 𝑅*_avr_* fell below a 25% fatigue threshold (𝑅_25%_). Raw resistance and corresponding deformation data are provided for reference (fig. S30). Following a predefined recovery period, stimulation resumed, with 𝑅_!_ and 𝑅_max_ recalibrated at each contraction cycle to accommodate the evolving physiological state of the bio-actuator.

To evaluate the efficacy of this strategy, we compared the fatigue index of muscles under closed-loop control against those subjected to a continuous, open-loop stimulation of equivalent duration. The closed-loop system demonstrated superior fatigue mitigation, preventing overstimulation and significantly attenuating performance degradation (Fig. 5D, fig. S31). Beyond immediate fatigue management, this sensor-driven architecture provides a scalable foundation for adaptive training protocols and the maintenance of homeostasis in long-term biohybrid robot operations.

## Discussion

In this work, we report an integrated biohybrid actuator system with bioelectronic interfaces capable of both eliciting and detecting muscle contractions with high fidelity. Furthermore, we deploy these individually addressable biohybrid actuators on a multi-actuator biohybrid robot platform. By demonstrating a proof-of-concept closed-loop control strategy that autonomously mitigates muscle fatigue, we establish a pathway to improve the energy efficiency, robustness, autonomy, and operational lifespan of biological actuators.

The soft, conformal nature of the embedded organic fibers creates a seamless interface that enables ultra-low-power stimulation (0.376 ± 0.034 mW), outperforming even state-of-the-art optogenetic methods in energy efficiency (*12, 25, 26*). This high-efficiency coupling allows for the development of multi-muscle systems where units can be paired with dedicated stimulation fibers for individualized, independent control. Our walking biohybrid robot validates this architecture: the selective, alternating recruitment of left and right muscle bundles produced a walking speed of 5.43 ± 0.79 mm/min and enabled directional steering. These results suggest that scaling this approach to more complex, multi-muscle architectures could achieve the rich, task-specific behaviors required for advanced soft robotic applications.

The integration of PEDOT fibers as intrinsic strain sensors provided high-resolution proprioceptive feedback, resolving contractile displacements down to ∼10 µm. While our current closed-loop setup utilizes a voltage-divider circuit requiring manual resistor selection, future iterations could replace this with a current-mode digital-to-analog converter (DAC). By delivering a fixed probe current and measuring the resulting voltage (𝑅 = 𝑉/𝐼), the system could implement an automated calibration process. Such an advancement would protect the fiber interface from overcurrent while significantly reducing the manual tuning required for autonomous operation.

Furthermore, while our study established the feasibility of closed-loop bioelectronic control, the optimization of control parameters, such as the fatigue threshold (x%), recovery intervals, and sampling windows, remains a fertile ground for future research. A systematic investigation into these variables will be essential for developing sophisticated muscle training protocols and advanced control laws that can adapt to the non-linear dynamics of biological tissues.

The current platform demonstrates the effectiveness of the bioelectronic interface in both stimulation and sensing as separate modalities. A primary objective for future development is the integration of these dual functions into a unified two-fiber-based interface. This will require a refined encapsulation strategy to ensure electrical isolation between stimulation and sensing signals without compromising actuation efficiency. Coupled with miniaturized, onboard electronics for signal processing, such a system would represent a significant step toward wireless, intelligent, and self-adaptive biohybrid robots.

In conclusion, our integrated platform demonstrates that a high-performance bioelectronic interface can bridge the gap between synthetic control and biological actuation. By enabling the selective recruitment and real-time monitoring of biohybrid muscle actuators, this work provides a generalizable toolbox for the next generation of autonomous biohybrid systems, promising enhanced performance and longevity for biomedical and soft robotic applications.

## Materials and Methods

### Fabrication and post-processing of PEDOT fibers

Aqueous PEDOT:PSS solution (Clevios™ PH 1000, Heraeus Epurio) was processed into fibers via wet-spinning into a 75% sulfuric acid (SA) coagulation bath (500 mL). The solution was delivered at a constant flow rate of 0.1, 0.2, and 0.3 mL/min using a syringe pump (Model No. 300, New Era Pump Systems, Inc.) and a 3 mL syringe coupled to a PTFE delivery tube (1/16” O.D., Kimble). To ensure axial uniformity and prevent morphological defects, a custom Teflon holder secured the delivery tube at the base of the SA bath, enabling vertical injection. This configuration leveraged the density gradient between the aqueous dope and the acid bath, allowing the fibers to rise buoyantly and float naturally at the surface (fig. S32). Following coagulation, the PEDOT fibers were harvested and rinsed with deionized (DI) water until the effluent reached a neutral pH. To maintain structural alignment and prevent coiling during dehydration, the rinsed fibers were secured at both ends for handling and suspended vertically with one end fixed. The fibers were then dried and annealed at 140 °C for 15 min to enhance crystallinity and mechanical stability. Post-processed fibers were mounted on cleanroom wipes for subsequent device integration. For extended storage, fibers were kept in DI water and re-immersed in the 75% SA bath for 2 hours prior to further processing.

### Gauge factor analysis for PEDOT fibers

The gauge factor was calculated from the electrical resistance (𝑅) and strain (𝜀) difference during the increasing strain segments. The current (𝐼) was measured via chronoamperometry at a constant applied bias of 5 mV using a PalmSens 4 potentiostat. The electrical resistance of the PEDOT fiber was calculated from the measured current using Ohm’s Law. The resistance was then filtered using a Savitzky-Golay filter to reduce high-frequency noise while preserving the physiological relevance of the signal. A dynamic baseline (𝑅_!_) was established using a low-pass filter to account for baseline drift due to the viscoelastic nature of conjugated polymers. The sensitivity was then expressed as the relative change in resistance (Δ𝑅/𝑅_0_):

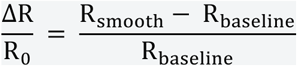

The mechanical strain (𝜀) profiles were measured from a dynamic mechanical analyzer (TA Instruments RSA G2). Because the electrical and mechanical measurements were recorded on independent hardware clocks, the resistance and strain data were temporally aligned using the Pearson correlation coefficient to synchronize the two signals. The gauge factor was then calculated from the slope of the Δ𝑅/𝑅_0_versus strain (𝜀) curve during these increasing strain segments, defined as:

Final values were averaged across multiple loading cycles to characterize the sensor’s linearity, sensitivity, and repeatability.

### Assembly of stimulation and sensing devices, and biohybrid walking robot skeletons

Device geometries, including two-post architectures for stimulation/sensing and four-post configurations for biohybrid walking skeletons, were designed in AutoCAD (dimensions in fig. S33-35). Master patterns were fabricated via stereolithography (SLA) using a Formlabs 3+ printer with Black Resin V5 (Formlabs). These patterns served as templates for constructing soft negative molds using Ecoflex 00-30 (fig. S36). After curing, the resin masters were manually delaminated, and the mold apertures were trimmed to ensure uniformity. To facilitate the subsequent release of polydimethylsiloxane (PDMS, SYLGARD™ 184) components, the Ecoflex molds were coated with 2 g of Parylene C using an SCS Labcoter 2 (Specialty Coating Systems™), which served as a critical anti-adhesion layer.

Conductive fiber integration was tailored to specific device functions. For stimulation devices, a lateral incision was made between the two post channels using a 1-cm-wide precision blade (Jetmore®). A 3.5-cm PEDOT fiber was embedded such that the fiber body was positioned within the incision, with turning points located at the channel necks and terminal ends extending through the post cavities (fig. S37). The fiber ends were secured to PDMS pads using surgical sutures (Walter Products™) to maintain optimal tension during subsequent processing. Electrical contact was established by interfacing lead wires with the fiber ends, reinforced with conductive silver epoxy (Circuitworks), and cured at 90 °C for 30 min (fig. S38). For sensing devices, a 2.5-cm fiber was placed within a single channel in a U-shaped configuration oriented orthogonally to the plane of the pillars and offset from the device midline (fig. S28). This placement concentrates strain on the fiber during muscle contraction, thus enhancing sensing resolution. The electrical interfacing followed the same silver epoxy protocol used for the stimulating units. For walking biobot skeletons, the integration procedure was duplicated across both sets of posts (fig. S39). To ensure that the electrical connections did not impede the mechanical gait of the biobot, thin, flexible Ball Grid Array (BGA) soldering conductor wires were used in place of standard metal lead wires. These softer, lightweight interconnects permitted unrestricted movement of the biobot skeleton during actuation. In all cases, circuit connectivity was verified via a multimeter (Fluke) to ensure stable electrical paths prior to the final encapsulation step.

PDMS was prepared at base-to-curing agent ratios of 10:1 (stimulation/biobot skeleton) or 20:1 (sensing) to modulate the mechanical modulus. The uncured PDMS was poured into the negative Ecoflex molds. To preserve the integrity of the delicate electrical connections and prevent fiber displacement, vacuum degassing was avoided. Instead, the assembly was left at room temperature for 1 h to allow for natural buoyancy-driven bubble release, supplemented by manual evacuation of trapped air at the distal ends of the post channels using a 200 µL pipette. Following a 2-h thermal cure at 90 °C, the devices were demolded by micro-dissection of the Ecoflex (fig. S40). To prevent air entrapment during subsequent upside-down muscle tissue engineering, 1-cm apertures were created using a circular punch. Sensing devices received an additional 2-g Parylene C coating followed by 30 layers of spray-coated styrene-butadiene-styrene (SBS) (10 mg/mL in toluene) using a 0.3-mm airbrush nozzle (25 psi, 15 cm standoff distance). During initial optimization of the elastomer barrier, styrene-ethylene-butylene-styrene (SEBS) and styrene-isobutylene-styrene (SIBS) were also evaluated using identical deposition parameters. This composite coating was applied to provide robust environmental and electrical insulation for the internal components while simultaneously promoting cell adhesion for the integrated muscle actuators.

### Muscle tissue engineering and biohybrid assembly

C2C12 myoblasts (ATCC) were maintained in complete medium [DMEM (Corning) supplemented with 5% fetal bovine serum (FBS, Gibco) and 1% Pen/Strep (Gibco)] at 37 °C and 5% CO2. Prior to tissue fabrication, myoblasts at < 70% confluency were harvested via trypsinization (0.05% trypsin, 10 min), neutralized with complete medium, and centrifuged at 1250 rpm for 5 min. The resulting cell pellet was resuspended in specialized C2C12 growth medium (GM, table S3). Hydrogel precursors—including Matrigel (Corning) thawed at 4 °C and fibrinogen (Sigma-Aldrich) dissolved in 0.9% NaCl and sterile-filtered (0.22 µm)—were prepared on ice to prevent premature gelation. Thrombin (Sigma-Aldrich) was dissolved in sterile 0.1% (w/v) BSA solution and kept on ice prior to use to maintain enzymatic stability and ensure consistent gelation kinetics during casting.

Tissue casting was performed using figure-eight–shaped PDMS wells (dimensions in fig. S41) fabricated via Ecoflex molding. These wells were secured to the centers of 6-well plate cavities using medical-grade adhesive (cured at 60 °C for 1 h). The entire assembly, along with the fabricated stimulation and sensing devices, underwent a dual-sterilization protocol: submersion in 75% ethanol for 30 min followed by UV irradiation. Post-sterilization, the wells were treated with a 1% (w/v) Pluronic F-127 solution for 1 h to create a non-adhesive surface, ensuring the engineered tissue would eventually anchor to the device posts rather than the casting well. The plates were then pre-cooled to -20 °C to further decelerate the thrombin-induced gelation during the casting process.

The cell/hydrogel mixture (detailed in table S4) (*6*) was formulated on ice by combining C2C12 cells, Matrigel, and fibrinogen. Upon the addition of thrombin, 225 µL of the mixture was rapidly dispensed into the chilled figure-eight wells. Immediately following injection, the stimulation or sensing devices were inverted and submerged into the wells, ensuring the post structures were centered within the hydrogel volume. For biohybrid walking skeletons, a specialized PDMS block with dual figure-eight wells (dimensions in fig. S42) was utilized to cast two independent muscle actuators simultaneously (fig. S15).

Constructs were incubated for 1 h to ensure complete gelation before the addition of 5 mL of GM (Day 0). On Day 1, the medium was transitioned to a differentiation medium (DM, table S5) supplemented with L-glutamine (Gibco), insulin-like growth factor-I (IGF-I, Sigma-Aldrich), and 6-aminocaproic acid (ACA, Sigma-Aldrich). This specific formulation was chosen to promote robust myogenesis while inhibiting fibrinolysis of the hydrogel scaffold. The biohybrid devices were transferred to new 6-well plates—omitting the figure-eight casting wells—on Day 3 or 5, contingent upon the observed degree of tissue compaction. This transfer allowed the matured muscle to interface directly with the device posts under tension, preparing the constructs for functional characterization starting on Day 10. For stimulation devices, the conductive fiber swells upon wetting; in some instances, this expansion leaves the fiber partially embedded in the surrounding tissue.

### Muscle stimulation and strain sensing

A precision source/measure unit (B2912B, Keysight) was employed for synchronized actuation and real-time strain sensing. The unit was controlled via PathWave BenchVue IV Curve Measurement Software to enable programmable, multi-channel protocols. For simultaneous sensing and actuation, the Keysight SMU was configured into two independent channels: Channel 1 delivered a constant bias voltage of 0.01 V to the sensing PEDOT fiber to monitor resistance changes (ΔR/R0), while Channel 2 generated the requisite stimulation waveforms to elicit muscle contraction.

Electrical connections for sensing (Channel 1) were established by interfacing alligator clips from the SMU to the integrated metal lead wires of the sensing device. To facilitate sterile, multi-well stimulation, the cap of a standard 6-well plate was modified with integrated platinum (Pt) and Ag/AgCl electrodes extending into each well (fig. S43). Prior to use, the modified cap was sterilized via 75% ethanol treatment and UV irradiation. The working electrode (either the Pt wire or the lead wires for stimulation PEDOT fiber) was connected to the positive terminal, while the circuit was completed via a ground-connected Ag/AgCl reference electrode submerged in the tissue culture medium. Specific stimulation parameters, including frequency and pulse width, are detailed in the respective figure legends. For selective actuation studies, the biobot was suspended using a custom-designed handle (dimensions in fig. S44), 3D-printed with a Formlabs 3+ printer using BioMed Clear Resin (Formlabs) and secured to the 6-well plate cap with medical-grade adhesive (fig. S16).

To correlate electrical signals with mechanical output, muscle contractions were captured using a widefield microscope (Nikon Ti2 Widefield). To maintain physiological viability during extended testing, all measurements were conducted within a dedicated microscope incubation chamber, regulated at 37 °C and 5% CO2.

### Power consumption analysis

To evaluate the electrical efficiency of the stimulation interface, the power consumption of PEDOT fiber electrodes was characterized and compared against standard platinum (Pt) wires. Power (𝑃) was calculated using the measured instantaneous current (𝐼) and the applied voltage (𝑉) according to the relationship:

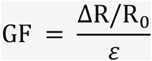

Measurements were performed using the Keysight B2912B SMU, recording the current response to the specific stimulation waveforms (e.g., 20 Hz square pulses) used during muscle stimulation. The average power consumption per contraction cycle was determined by integrating the instantaneous power over the pulse duration (𝜏) and dividing by the total cycle period (𝑇):

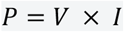

For both electrode types, power analysis was conducted in the same conductive tissue culture medium to account for environmental impedance. The reduction in power consumption for PEDOT fibers was attributed to their high effective surface area and lower interfacial impedance compared to Pt, facilitating efficient charge injection into the biological tissue at lower overpotentials.

### Alternative stimulation code for biohybrid robots

Alternating muscle stimulation was applied via an ADALM1000 SMU (Analog Devices) and was programmed using the pysmu library. Code is available at https://github.com/dasorachel/Muscle_Stimulation.git.

### Closed-loop control architecture

To quantify the resistance changes in the PEDOT fiber strain gauge, a voltage divider was constructed by placing the fiber in series with a reference resistor (𝑅_1_). The central node voltage (*Vout*) was monitored via the 10-bit analog-to-digital converter (ADC) of an Arduino UNO R3. To prevent irreversible electrochemical damage to the fiber, the potential across the strain gauge (𝑉*_fiber_*) was restricted to < 2 V. Given a 5 V supply from the microcontroller’s General Purpose Input/Output (GPIO) pins, *R1* was selected based on the baseline resistance of each specific fiber sample (𝑅*_fiber_*), such that:

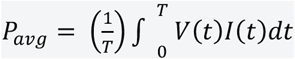

Empirical testing across samples (1 kΩ to 100 kΩ) indicated that selecting 𝑅_1_ ≈ 2 × 𝑅*_fiber_* ensured safety margins were maintained even during maximum muscle contraction (∼ 25% resistance change).

Due to the non-ideal insulation of the sensing fibers, a continuous measurement bias would introduce unwanted DC offsets and alter the electric field distribution during stimulation. We implemented an active-sampling protocol that leverages the high-impedance state of GPIO pins to “isolate” the sensing circuit when not in use.

As illustrated in Figure 5A, the sensing pins (Pins 2–4) remain in a high-impedance state by default. During a sampling event, the circuit is momentarily activated:

1. Activation: Pin 2 is driven HIGH (5 V) and Pin 4 is driven LOW (GND).
2. Stabilization: A brief delay is introduced to allow for the charging of stray capacitance and the stabilization of the divider node.
3. Acquisition: Pin 3 triggers the ADC conversion. To avoid signal contamination from the pulse-width modulation (PWM) stimulation, the ADC event is synchronized to trigger only during the “low” state of the PWM cycle.
4. Discharge and Reset: Pins 2–4 are briefly grounded to fully discharge the circuit before returning to a high-impedance state.

### Muscle fatigue characterization

The fatigue index was determined by comparing the mechanical output at the end of a stimulation trial to the initial peak performance. The index (𝐹𝐼) is defined as:

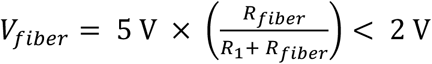

Where: 𝐹_+326245_: The average peak force measured during the first stable contraction cycles; 𝐹_*2345_: The peak force measured during the final cycle of the stimulation period. A fatigue index of 0% represents no loss in performance, while higher percentages indicate greater degrees of tissue exhaustion. Fatigue indices were compared across different control architectures, specifically evaluating the performance of open-loop stimulation versus closed-loop adaptive control to determine the impact of sensing-informed feedback on muscle performance. All statistical comparisons were performed using Student’s t-test with a significance threshold of p < 0.05.

### Data synchronization and processing for closed-loop control

The optical imaging and electrical sensing data were synchronized using a fast normalized cross-correlation method (*44*) to match the commanded stimulation triggers with the onset of mechanical movement.

### Muscle deformation characterizations

Muscle contraction-induced deformation was quantified using a custom image processing pipeline implemented in Python. High-speed TIFF stacks were processed to track the displacement of the device posts. For each frame, a region of interest (ROI) was defined around the post edge and averaged along the vertical axis to generate a one-dimensional intensity profile. To achieve subpixel resolution, the intensity profiles were smoothed using a Savitzky-Golay filter. The post edge was identified by locating the zero-crossing of the second derivative, specifically selecting the crossing point closest to the maximum absolute slope of the first derivative. Linear interpolation was applied at the zero-crossing to determine the edge position with subpixel accuracy. The pixel-wise displacement was converted to physical distance (µm) using a calibration factor of 4.74 µm/pixel. This factor was computationally determined by the Nikon imaging software (NIS-Elements) based on the specific optical parameters and metadata of the objective (CFI Plan Achro 1X, Nikon) used during acquisition.

### Biobot tracking

Video sequences of the walking biobots were processed using the OpenCV library. In the first frame, a Region of Interest (ROI) was manually defined to isolate a specific feature on the biobot. This ROI was then extracted from all sequential frames to generate a chronological image stack, preserving the spatial context of the biobot’s gait. Locomotion was quantified using a fast normalized cross-correlation template matching algorithm (*44*) from the skimage library. A sub-region from the initial ROI served as the tracking template. For each subsequent frame, the algorithm performed a sliding-window search to identify the pixel coordinates (x, y) that maximized the correlation coefficient between the template and the current frame. The trajectory was plotted from the pixel coordinates of each frame and converted to physical distance (µm) using a calibration factor of 4.74 µm/pixel, as described in the **Muscle deformation characterization** section. To visualize and verify the tracking accuracy, the calculated centroid of the matched region was marked on each frame, and a bounding rectangle was overlaid to confirm stable alignment. The resulting coordinate data provided a frame-by-frame motion path, which was then used to calculate the following kinematic parameters: 1. Total Displacement: The cumulative Euclidean distance traveled from the starting coordinates (*x0, y0*). 2. Instantaneous Velocity: Derived from the change in position between frames relative to the calibrated frame rate (FPS). 3. Gait Periodicity: Extracted by analyzing the fluctuations in the *x* and *y* position vectors over time, correlated to the 1 Hz stimulation frequency. The final trajectories and correlation metrics were exported as standardized CSV files for further statistical analysis and comparison across different stimulation voltages and pulse widths.

### Electrochemical impedance spectroscopy (EIS)

The electrochemical properties of the PEDOT fibers were characterized via EIS using a PalmSens4 potentiostat. To isolate the fiber-electrolyte interface, a custom test cell was constructed by sandwiching the fibers between two PDMS blocks featuring center apertures. Electrical contact was established using conductive silver epoxy and metal lead wires external to the electrolyte-wetted region. The EIS measurements were conducted in a two-electrode configuration within PBS: the PEDOT fiber served as the working electrode, while a submerged Ag/AgCl electrode acted as the combined counter and reference electrode (fig. S45).

Impedance spectra were acquired over a frequency range of 0.01 Hz to 10 kHz with a sinusoidal AC amplitude of 10 mV. To quantify the interfacial dynamics, the resulting data were fit to an equivalent electrical circuit (Randles circuit, fig. S2) model using the impedance.py Python package.

### Histological and viability analysis

To evaluate myogenic maturation and sarcomere organization, muscle tissues were harvested by sectioning the device posts. The constructs were fixed and permeabilized in 4% paraformaldehyde (PFA) supplemented with 0.5% Triton X-100 at 4 °C for 24 h. Following three 30-min rinses in PBS, non-specific binding was blocked using 5% goat serum (Gibco) with 0.5% Triton X-100 at 4 °C for 24 h. Tissues were then incubated with primary antibodies—Myosin 4 Monoclonal Antibody (1:400, eBioscience™) and Anti-Sarcomeric Alpha Actinin (1:200, Abcam)—at 4 °C for 72 h. After triple-washing in PBS, secondary antibodies—Alexa Fluor™ 488 (Goat anti-Mouse, 1:250) and Alexa Fluor™ 568 (Goat anti-Rabbit, 1:200)—were applied at 4 °C for an additional 72 h. The extended incubation periods were utilized to ensure thorough antibody penetration throughout the dense 3D muscle constructs. Finally, tissues were mounted in ProLong™ Gold Antifade Mountant with DAPI (Invitrogen™) on glass-bottom petri dishes and secured with coverslips for imaging.

The metabolic health of the integrated biological actuators was verified using a LIVE/DEAD™ Viability/Cytotoxicity Kit (Invitrogen™). Harvested tissues were incubated in a solution of EthD-1 (4 µM) and Calcein AM (2 µM) at 37 °C for 2 h. Following three PBS washes, the constructs were mounted in ProLong™ Gold Antifade Mountant (Invitrogen™) on glass-bottom petri dishes and secured with coverslips for imaging.

High-resolution fluorescence and viability imaging were performed using a laser scanning confocal microscope (Nikon A1R). Z-stacks were acquired to visualize the internal cellular distribution and protein expression within the 3D biohybrid structures.

### Mechanical testing and data analysis of PEDOT fibers

To determine the mechanical properties of the PEDOT fibers, tensile tests were performed on the fibers using an MTS Criterion Model 42 (C42) universal test stand with an MTS Bionix EnviroBath installed. All samples were tested in 1x PBS at 37°C. Samples were loaded into specialized hooks that minimized stress concentrations in the test region due to clamping (fig. S46). Once installed, the machine was positioned to remove any slack that was visible in the fiber, and the gauge length was measured. Fibers were then cycled five times at a strain rate of 10%/min for a max strain of either 10% (for Group 1) or 7.5% (for Groups 2 and 3). This difference in max strain was based on preliminary tests where Group 2 and 3 fibers would fail if cyclically strained to 10%. After the cycles, the fiber was relaxed and left to dwell for approximately 20 seconds before being strained again at 10%/min until failure. Force and displacement data from these tests were sampled at 10 Hz.

To determine an approximate cross-sectional geometry of the fibers, samples that had been mechanically tested were laid out in Petri dishes filled with 1x PBS and taped at the ends so that they remained approximately straight during imaging. Then, a series of images were taken along the length of each fiber using an AmScope MU023 digital CMOS camera connected to an AmScope trinocular microscope. All images were taken so that the length of the fiber was approximately horizontal in the image. The diameter of the fiber along its length was then estimated using a custom image analysis script in Matlab (MathWorks). Briefly, images were binarized to isolate the dark fibers from the white background. Then, the top and bottom edges of the fiber were identified using a two-pixel kernel (K=[-1,1]^T^), and the midline of the fiber was estimated as the average of these two edges. Linear functions were fitted to the midline data in overlapping windows of 11 pixels to find the local direction of the fiber midline. A perpendicular line was then constructed at the middle of the window, and the intersections of this perpendicular line with the top and bottom lines were found. The distance between these intersection points was then calculated as the local diameter of the fiber. From this analysis, a distribution of diameter values for each fiber was constructed.

Using the force-displacement data from the mechanical tests and the cross-sectional data from the microscopy study, the material properties of the fibers were determined by fitting the data to an incompressible Neo-Hookean material model. This model was selected because it captured both the cyclic and pull-to-failure data well with minimal parameters. To account for the uncertainty introduced by the variability in fiber diameter, a bootstrapping approach was used. For each replicate, the diameter data were resampled, the cross-sectional area was calculated assuming a circular cross-section, and the average area was used to calculate the engineering stress (force / average rest cross-sectional area). To account for the fiber not being at its rest length (either due to being stretched or slightly slack) at the start of the test, an offset in the rest length was also fitted to the data. This fitting helps to ensure that stress-strain data passes through zero stress for zero strain. The offset parameter and the stiffness parameter of the Neo-Hookean model (C1) were fitted simultaneously by minimizing the total squared error between the model data and experimental data from the cyclic strain tests. The Young’s modulus was then calculated as 6C1, and the bending modulus was calculated as Young’s modulus times the average second moment of area (also estimated from the resampled diameter data assuming a circular cross-section). This procedure was repeated 10,000 times for each fiber to estimate the uncertainty in the material parameters.

### Finite element analysis

The mechanical response of the device pillars under lateral contractile forces was simulated using COMSOL Multiphysics (Version 6.3). The PDMS pillar geometry was modeled as a composite structure consisting of two coaxial cylindrical segments: a primary lower cylinder (height H = 5 mm, diameter D = 1.5 mm) and a structural upper cylinder (H = 1 mm, D = 3 mm). To accurately replicate the experimental conditions, a 2-μm-thick outer shell was incorporated into the model to represent the Parylene C protective coating. Simulations were performed using the Solid Mechanics module under a stationary study. The base of the lower cylinder was defined with a fixed constraint, while the lateral contractile force was applied as a boundary load along the circular edge at the junction of the two cylinders. This loading configuration replicates the attachment point of the muscle and the resultant force vector during stimulated contraction. PDMS was modeled at mixing ratios of 10:1 and 20:1 (base to curing agent) for pure-PDMS and Parylene-coated pillars, respectively, to match experimental fabrication. Specific values for Young’s modulus, Poisson’s ratio, and density for each material layer are detailed in Supplementary table S6. This finite element model was used to validate the analytical force-displacement relationships derived from Euler-Bernoulli beam theory and to assess the influence of the Parylene C coating on overall pillar stiffness.

### Statistical analysis

Individual data points were analyzed using Python (SciPy and Pandas libraries) and are presented as means ± std. Student’s 𝑡-test was applied to compare the means of two groups using either paired or two-sample functions. P values were used to indicate statistical significance (*P < 0.05, **P < 0.01, and ***P < 0.001).

## Acknowledgments

Imaging work was performed at the Northwestern University Center for Advanced Microscopy (RRID: SCR_020996), generously supported by NCI CCSG P30 CA060553 awarded to the Robert H Lurie Comprehensive Cancer Center. This research used the Micro/Nano Fabrication Facility at Northwestern University.

## Author contributions

Conceptualization and design: X.X., Y.Z., A.S., V.A.W., and J.R. Data acquisition, analysis, or interpretation: X.X., R.W., W.X., M.J.B., R.D. Computational modeling: J.L. Writing: X.X., Y.Z., R.W., M.J.B., R.D., J.L., J.H., V.A.W., T.C., and J.R.

## Data and materials availability

All data are available in the main text or the Supplementary Materials. Code and scripts are available upon request.

## Supplementary Materials

This PDF file includes:

Figs. S1 to S46

Tables S1 to S6

## Other Supplementary Material for this manuscript includes the following

Movies S1 to S7

