## Supplementary Materials for "Biohybrid Robots with Embedded Conductive Fibers for Actuation, Sensing, and Closed-loop Control"

Xinran Xie *et al.*

**This PDF file includes:**

Figs. S1 to S46

Tables S1 to S6

**Other Supplementary Material for this manuscript includes the following:**

Movies S1 to S7

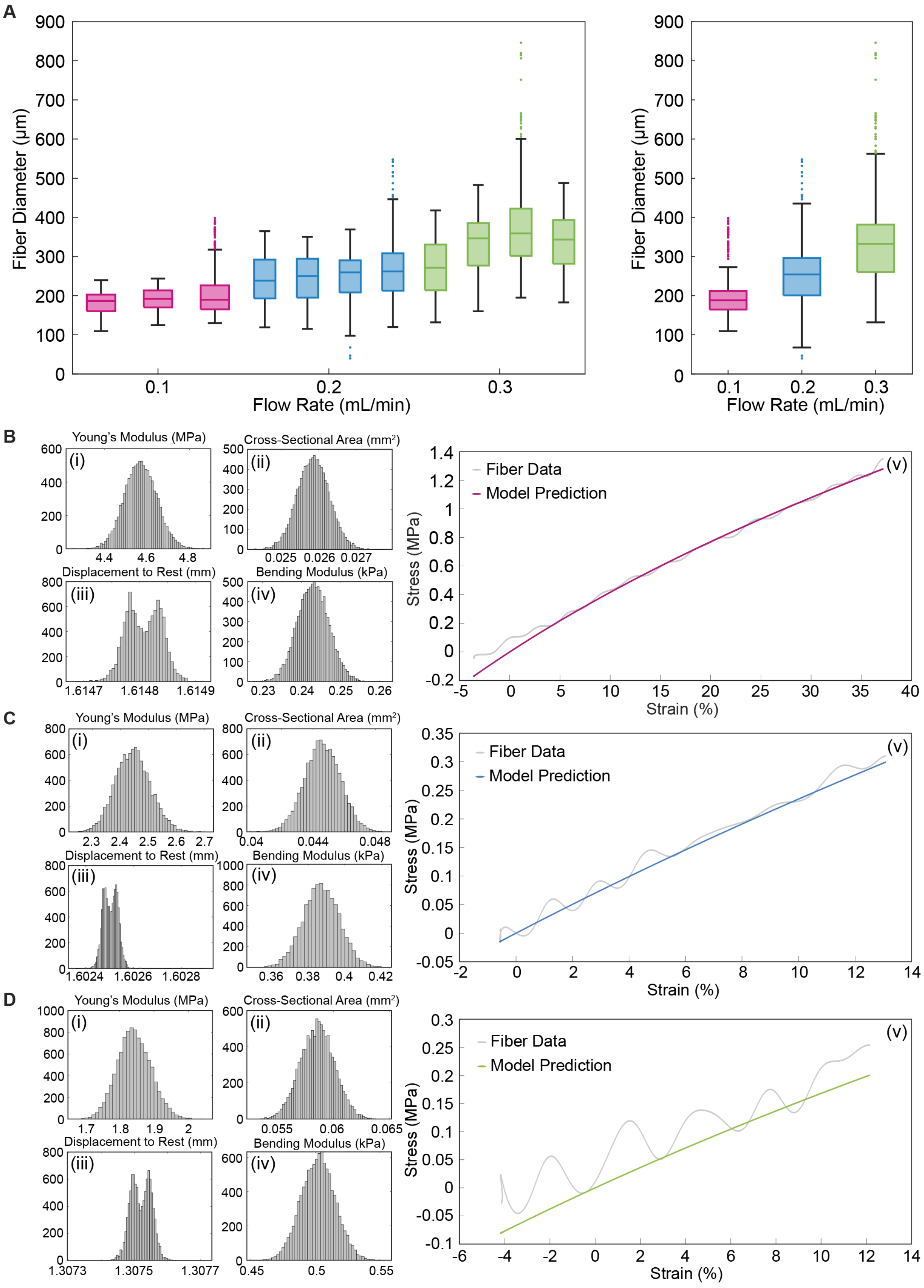
A
Fiber Diameter (μm)
Flow Rate (mL/min)
B
Young's Modulus (MPa)
Cross-Sectional Area (mm²)
Displacement to Rest (mm)
Bending Modulus (kPa)
(i)
(ii)
(iii)
(iv)
(v)
Fiber Data
Model Prediction
Stress (MPa)
Strain (%)
C
Young's Modulus (MPa)
Cross-Sectional Area (mm²)
Displacement to Rest (mm)
Bending Modulus (kPa)
(i)
(ii)
(iii)
(iv)
(v)
Fiber Data
Model Prediction
Stress (MPa)
Strain (%)
D
Young's Modulus (MPa)
Cross-Sectional Area (mm²)
Displacement to Rest (mm)
Bending Modulus (kPa)
(i)
(ii)
(iii)
(iv)
(v)
Fiber Data
Model Prediction
Stress (MPa)
Strain (%)

**Fig. S1.**

**Morphological and mechanical characterization of wet-spun PEDOT fibers across varying flow rates.** (**A**) Geometric analysis of PEDOT fibers, including individual fiber diameters and group-wise statistical distributions (N = 3 for 0.1 mL/min; N = 4 for 0.2 and 0.3 mL/min). (**B**–**D**) Comprehensive mechanical profiling for fibers fabricated at spinning speeds of 0.1, 0.2, and 0.3 mL/min: (i) Young's modulus; (ii) corresponding cross-sectional area measurements; (iii) displacement-to-rest characterization; (iv) bending modulus; (v) representative pull-to-failure stress–strain curves. Experimental data in (v) are fitted with the Neo-Hookean hyperelastic model to characterize the non-linear mechanical response.

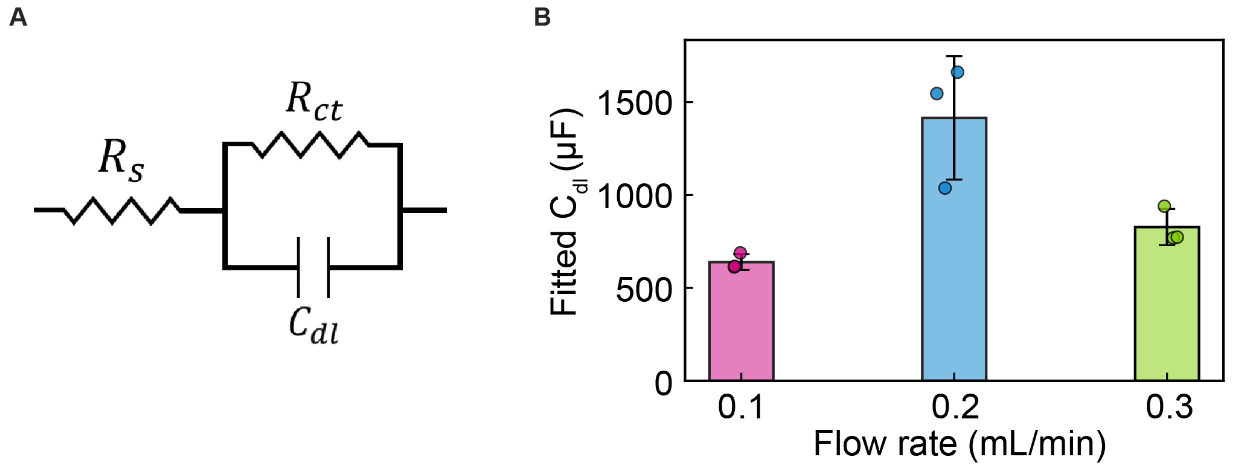


**Fig. S2.**
**Equivalent circuit modeling and capacitive characterization of PEDOT fibers.** (**A**) Schematic of the Randles equivalent circuit utilized for fitting the electrochemical impedance spectroscopy (EIS) data. The circuit includes elements representing solution resistance ($R_s$), charge-transfer resistance ($R_{ct}$), and the double-layer capacitance ($C_{dl}$). (**B**) Calculated double-layer capacitance values derived from the circuit fitting for PEDOT fibers fabricated at flow rates of 0.1, 0.2, and 0.3 mL/min. Data are presented as means ± std (N = 3 per group).

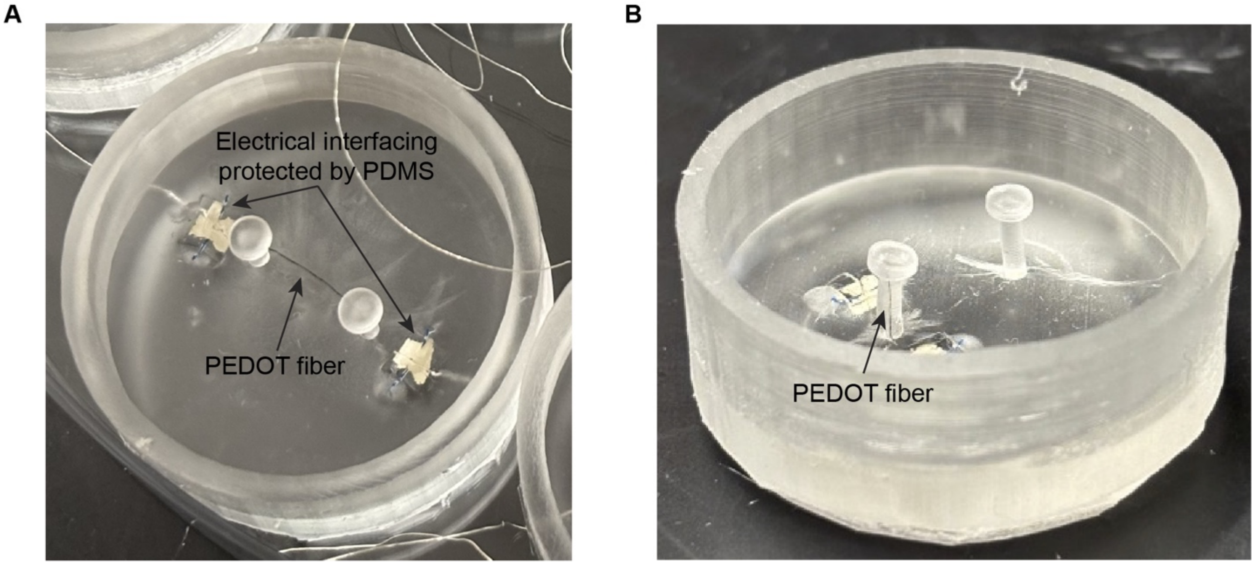


**Fig. S3.**
**Optical photographs of the PEDOT fiber-based stimulation and sensing devices.** (**A**) Photograph of the assembled stimulation device, illustrating the mechanical and electrical connection interface for the PEDOT fiber electrodes. (**B**) Photograph of the sensing device configuration, highlighting the precise integration of the PEDOT fiber within a single flexible PDMS pillar for localized strain monitoring.

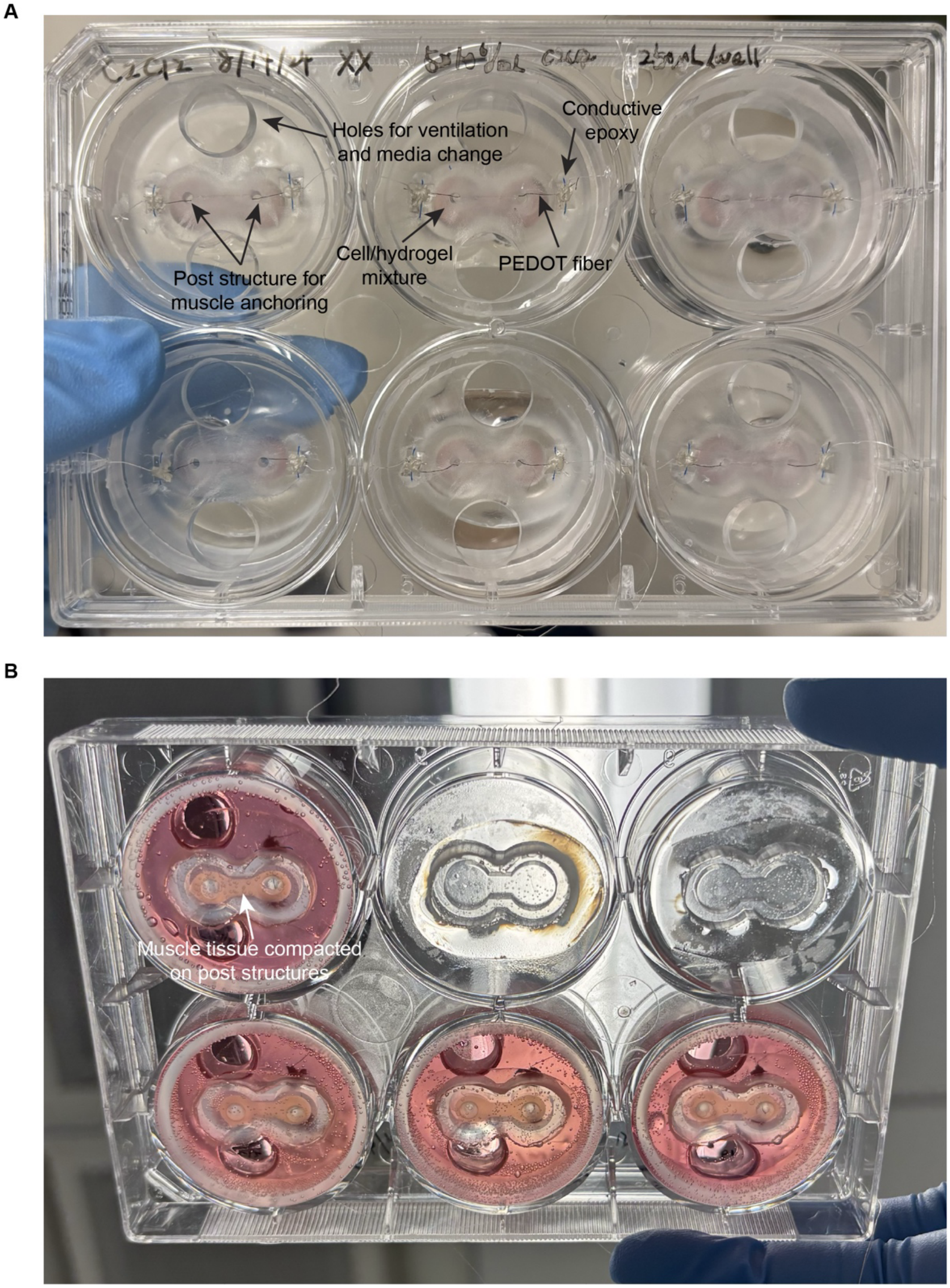


**Fig. S4.**
**Fabrication and maturation of engineered skeletal muscle tissues.** (**A**) Optical photograph of the muscle casting procedure, showing the positioning of the dual-post structures within the cell-laden hydrogel mixture. (**B**) Optical photograph characterizing tissue maturation and compaction. Complete delamination from the figure-eight casting well boundary indicates successful tissue remodeling, at which point the biohybrid devices are prepared for transfer to a standard culture plate for subsequent experimentation.

A

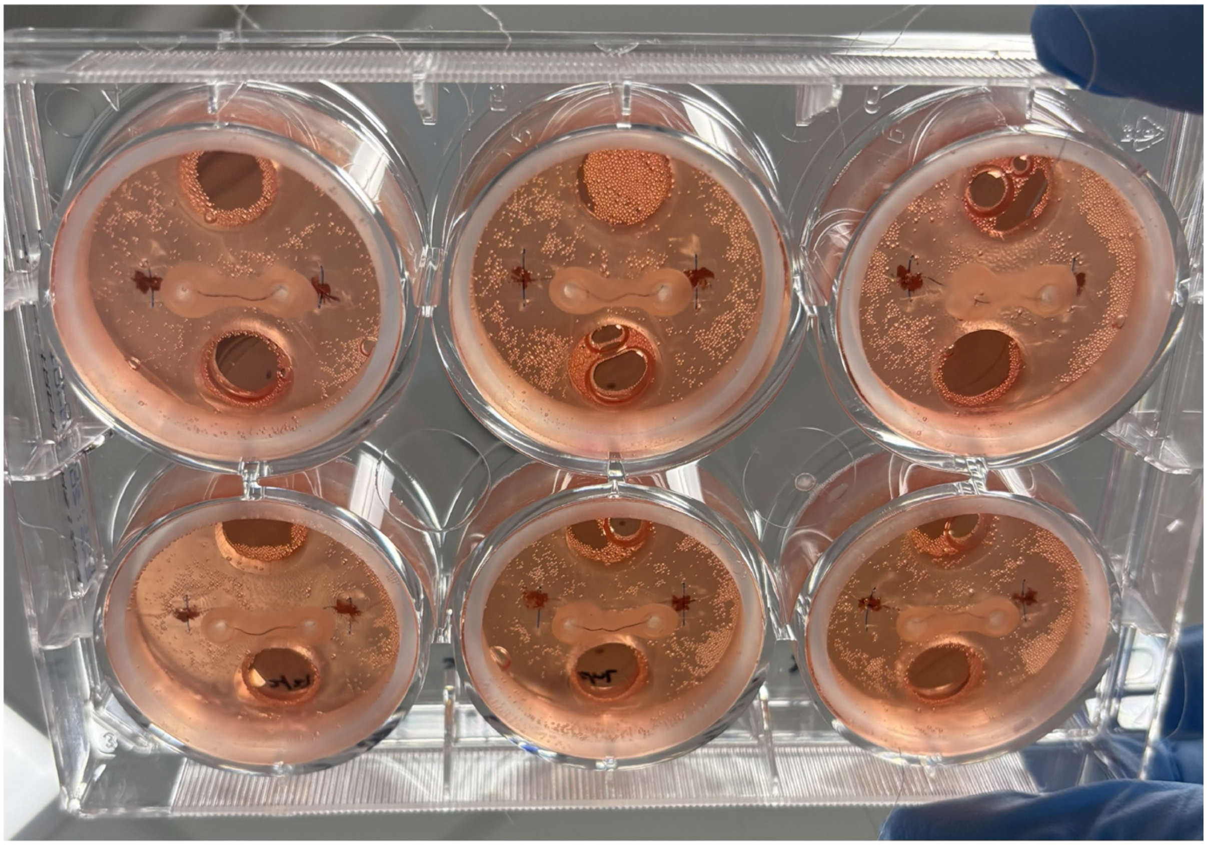

**Fig. S5.**

**Integration and anchoring of skeletal muscle tissues onto PEDOT fiber interfaces.** (**A**) Optical photograph of the biohybrid assembly following the removal of the casting wells. The engineered muscle tissue is shown anchored to the dual-post structures and circumferentially wrapping the PEDOT fiber, ensuring a robust mechanical and electrical interface for subsequent stimulation.

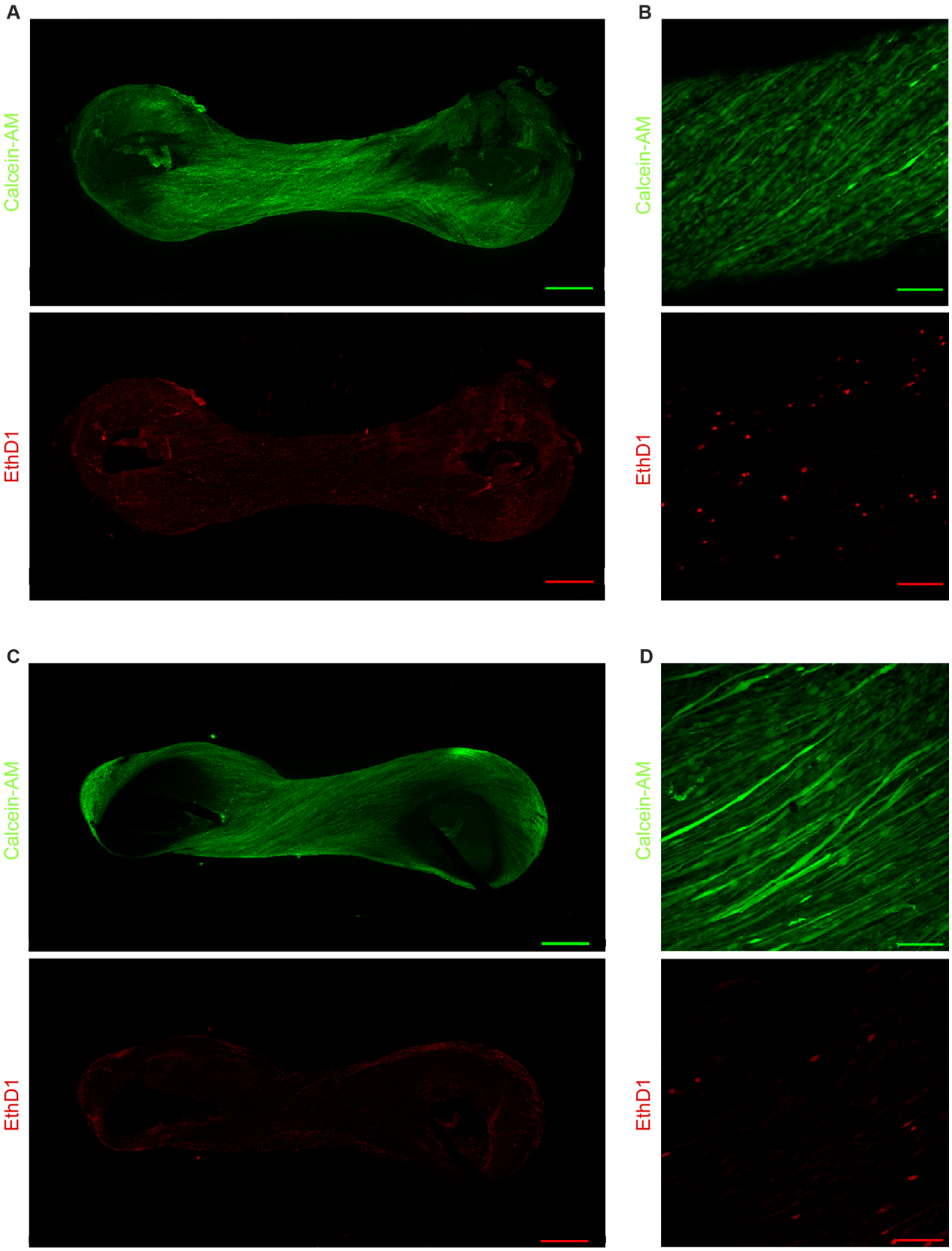


**Fig. S6.**

**Single-channel fluorescence micrographs of live/dead viability assays.** (**A**, **B**) Representative fluorescence images of the control muscle group: (**A**) whole-tissue overview (scale bar, 1 mm) and (**B**) high-magnification view (scale bar, 100 µm). (**C**, **D**) Representative fluorescence images of the PEDOT fiber-embedded muscle group: (**C**) whole-tissue overview (scale bar, 1 mm) and (**D**) high-magnification view (scale bar, 100 µm). Single-channel images are provided to demonstrate the distribution of live (green) and dead (red) cells.

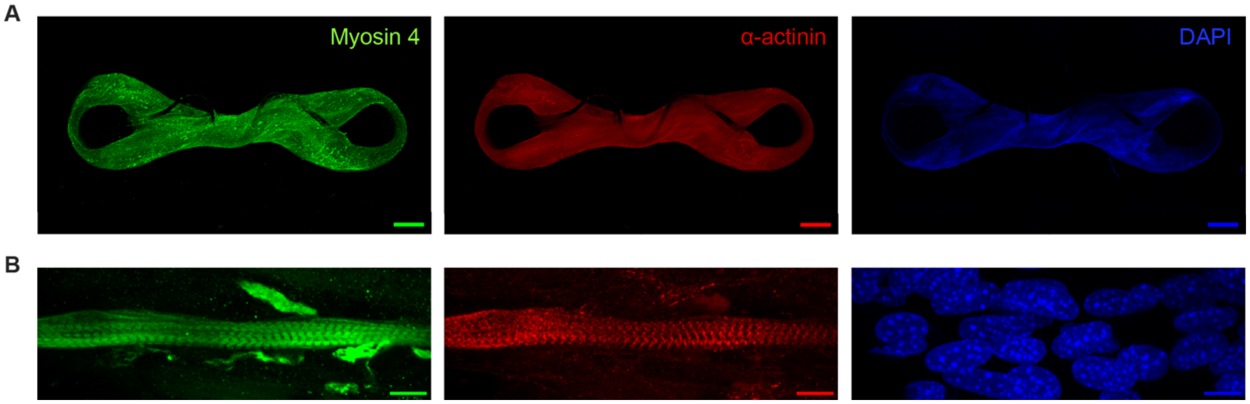


**Fig. S7.**

**Single-channel immunofluorescence characterization of muscle maturity.** (**A**) Whole-tissue overview of a fully differentiated, fiber-embedded muscle, showing the global distribution of contractile proteins. Scale bar, 1 mm. (**B**) High-magnification micrographs highlighting the sarcomeric organization. Micrographs display individual channels for myosin heavy chain (MHC) (Myosin 4, green), sarcomeric-α-actinin (red), and DAPI-stained nuclei (blue). The clear striation patterns in both MHC and α-actinin channels confirm the structural maturation of the biohybrid actuator. Scale bar, 10 μm.

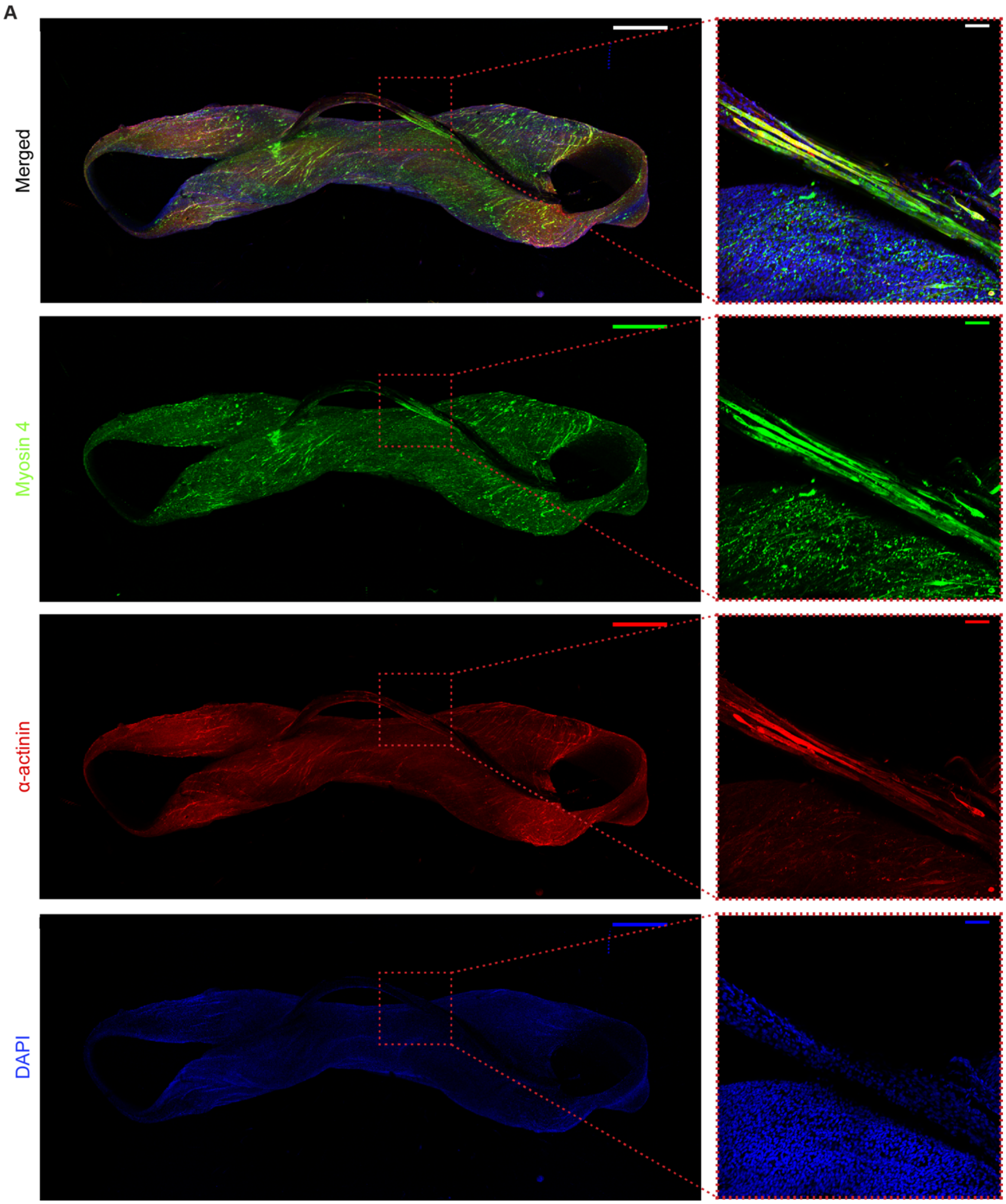


**Fig. S8.**
**Directional alignment of myofibers along the PEDOT fiber interface.** (**A**) Merged and single-channel immunofluorescence micrographs illustrating the longitudinal alignment of differentiated myofibers along the axis of the PEDOT fiber. Micrographs show myosin heavy chain (MHC) (Myosin 4, green), sarcomeric-α-actinin (red), and DAPI-stained nuclei (blue). The whole-tissue view (scale bar, 1 mm) demonstrates global tissue orientation, while the high-magnification inset (scale bar, 100 μm) highlights the high degree of cellular anisotropy and uniaxial organization achieved at the fiber interface.

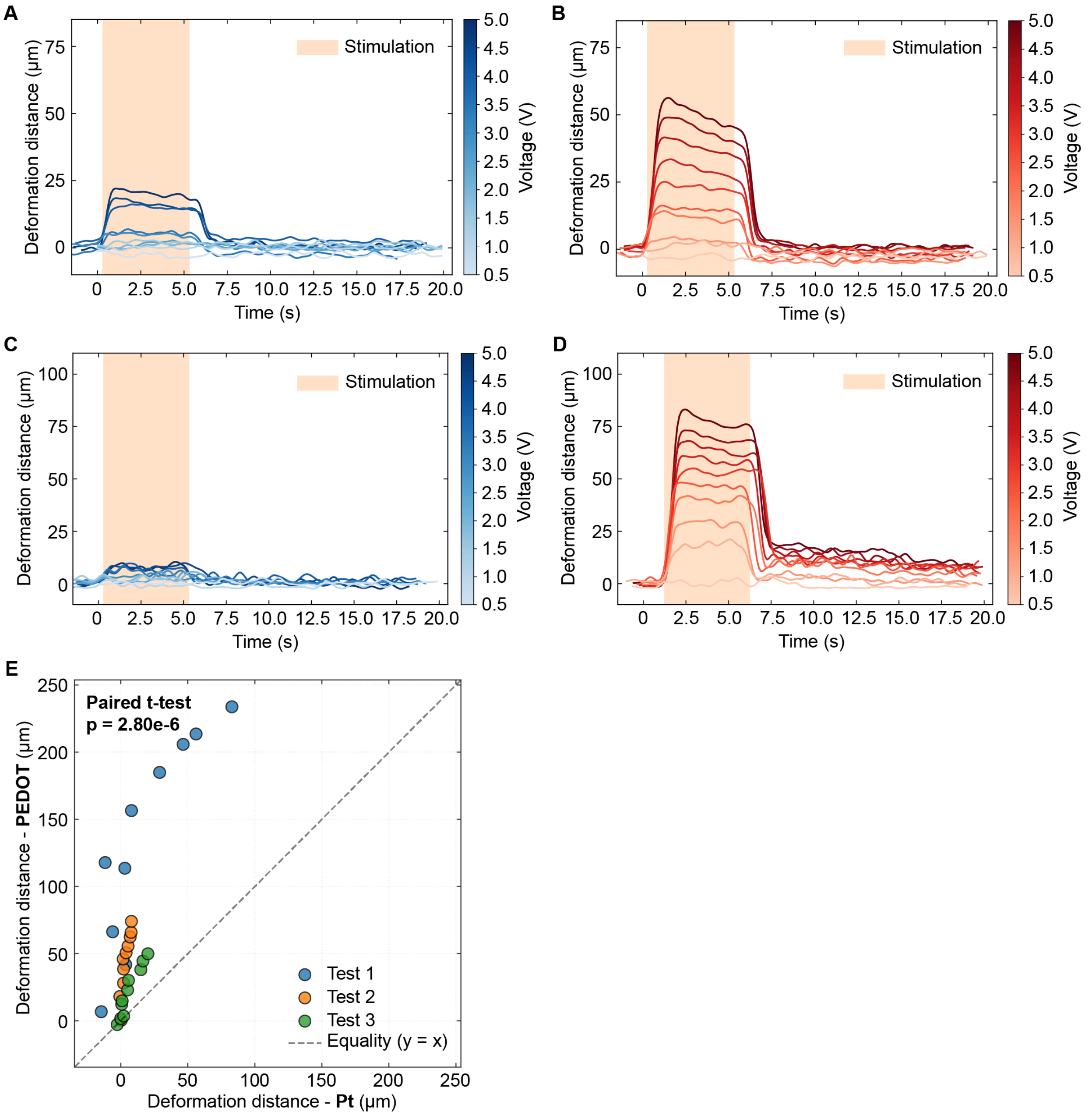


**Fig. S9.**

**Longitudinal comparison of stimulation efficacy between external Pt wire and embedded PEDOT fiber electrodes.** (**A**–**D**) Representative time-resolved contraction profiles comparing stimulation methods at different maturation stages: (**A**, **B**) Day 10 and (**C**, **D**) Day 17. Contractions were elicited using either (**A**, **C**) an external Pt wire electrode or (**B**, **D**) the embedded PEDOT fiber within the same muscle sample. All stimulations were conducted at 20 Hz with a 5 ms pulse width for a 5 s duration. (**E**) Statistical comparison of peak contraction amplitudes using a paired two-tailed Student's t-test. The embedded PEDOT fiber demonstrates a significant improvement in actuation outcome compared to external Pt stimulation (N = 30 measurements derived from three independent biological replicates).

A

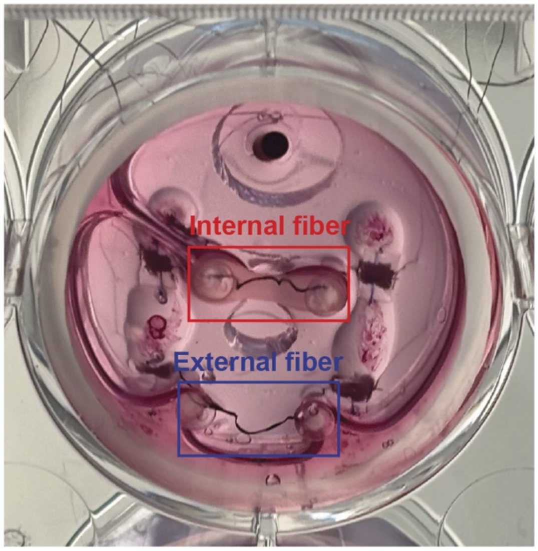


**Fig. S10.**
**Experimental configuration for isolating the effects of electrode placement and material composition.** (**A**) Optical photograph of the modified stimulation platform designed to decouple the influences of electrode material and spatial placement. The assembly features an engineered muscle tissue integrated with an internal, tissue-embedded PEDOT fiber, while a second PEDOT fiber is positioned externally within the culture medium. This configuration serves to confirm that the enhanced actuation outcomes observed are primarily driven by the proximity of the bio-interface rather than the intrinsic properties of the organic conductor alone.

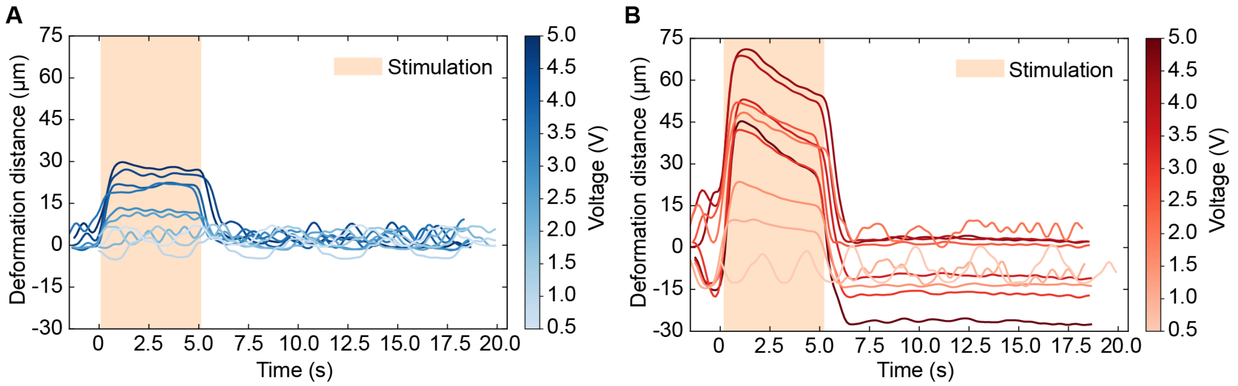


**Fig. S11.**

**Comparative analysis of actuation efficacy based on electrode spatial positioning.** Representative profiles of stimulation-induced deformation within a single muscle sample, comparing the influence of PEDOT fiber placement. (**A**) Contraction response elicited by the external PEDOT fiber positioned in the culture medium. (**B**) Contraction response elicited by the internal, tissue-embedded PEDOT fiber. All stimulations were performed at 20 Hz with a 5 ms pulse width for a 5 s duration. This direct comparison demonstrates that the intimate integration of the fiber within the tissue architecture significantly enhances electro-mechanical coupling compared to extracellular field stimulation using the same organic material.

A

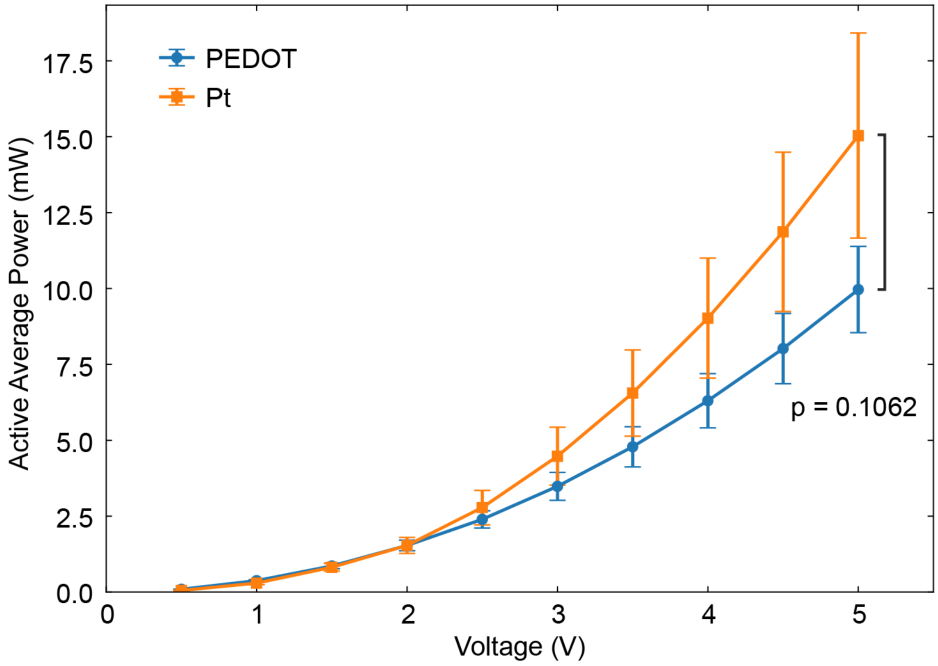


**Fig. S12.**

**Comparative power consumption analysis of stimulation interfaces.** (**A**) Average power consumption measured during the 5-s stimulation window as a function of applied voltage. The analysis compares the energy requirements of the external Pt wire electrode versus the integrated PEDOT fiber electrode. All stimulations were performed at 20 Hz with a 5 ms pulse width for a 5 s duration. The data illustrate the power efficiency gains achieved by the tissue-embedded PEDOT fiber interface, particularly at higher actuation voltages.

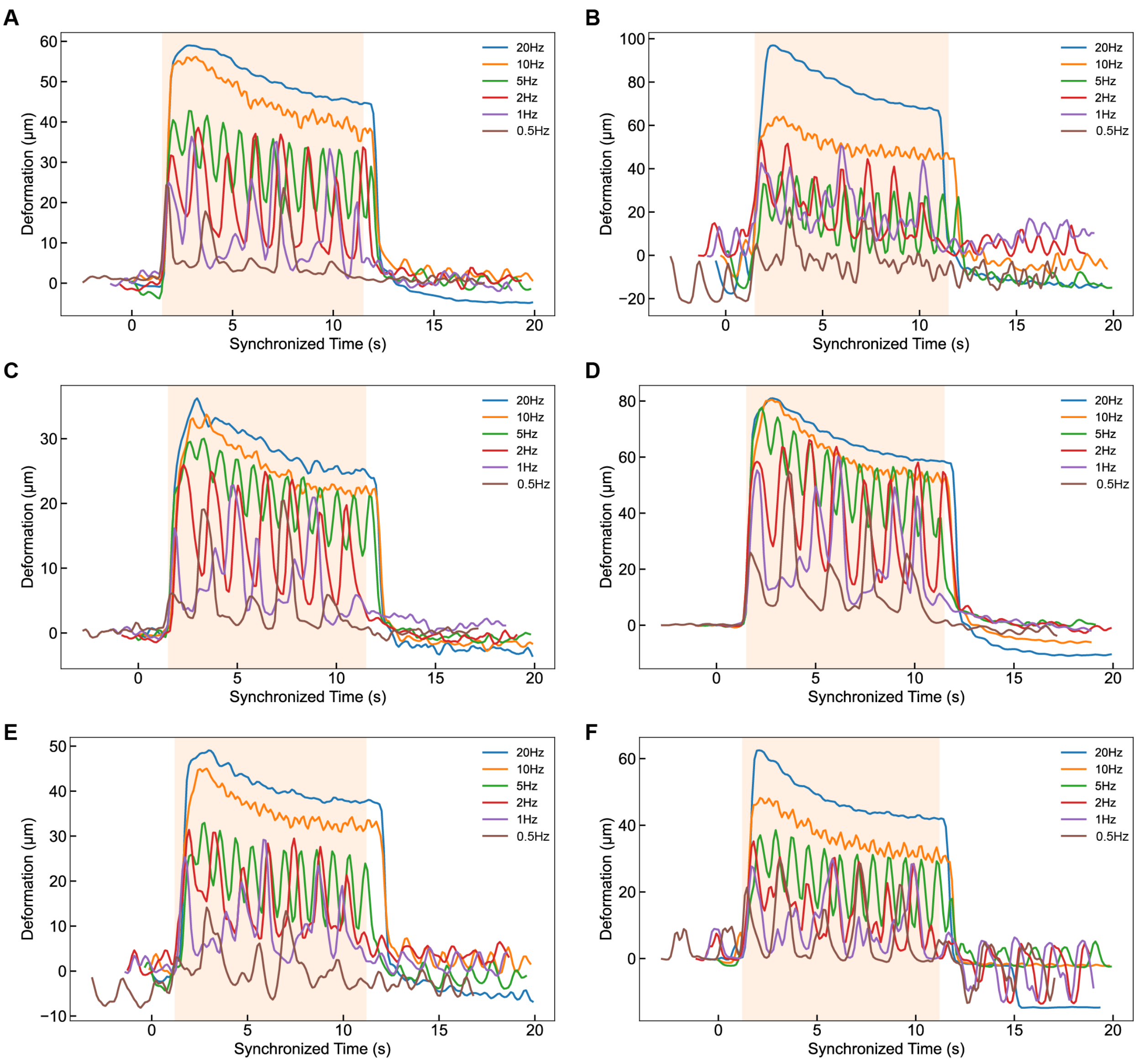


**Fig. S13.**

**Individual frequency-dependent contraction profiles of fiber-embedded muscle units.** (A-F) Contraction response of six independent engineered muscle replicates (N = 6) measured on Day 21 of maturation. Actuation was elicited via integrated, tissue-embedded PEDOT fibers across a stimulation frequency range of 0.5 to 20 Hz. All trials were conducted at 5 V with a 5 ms pulse width for a 10 s duration. These individual profiles characterize the consistent electromechanical coupling and the characteristic increase in contractile performance at higher stimulation frequencies, demonstrating the functional robustness of the biohybrid actuators at a late stage of maturation.

**A**

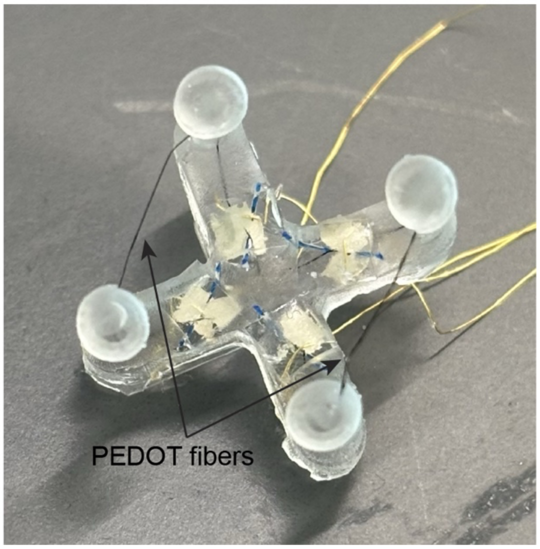


**Fig. S14.**
**Structural architecture of the biohybrid walking robot (biobot) skeleton.** (**A**) Optical photograph of the biobot skeleton, illustrating the strategic integration of two parallel PEDOT fibers within the structural frame. This configuration provides the dual-electrode framework necessary for selective actuation of individual muscle units and coordinated locomotion of the robotic system.

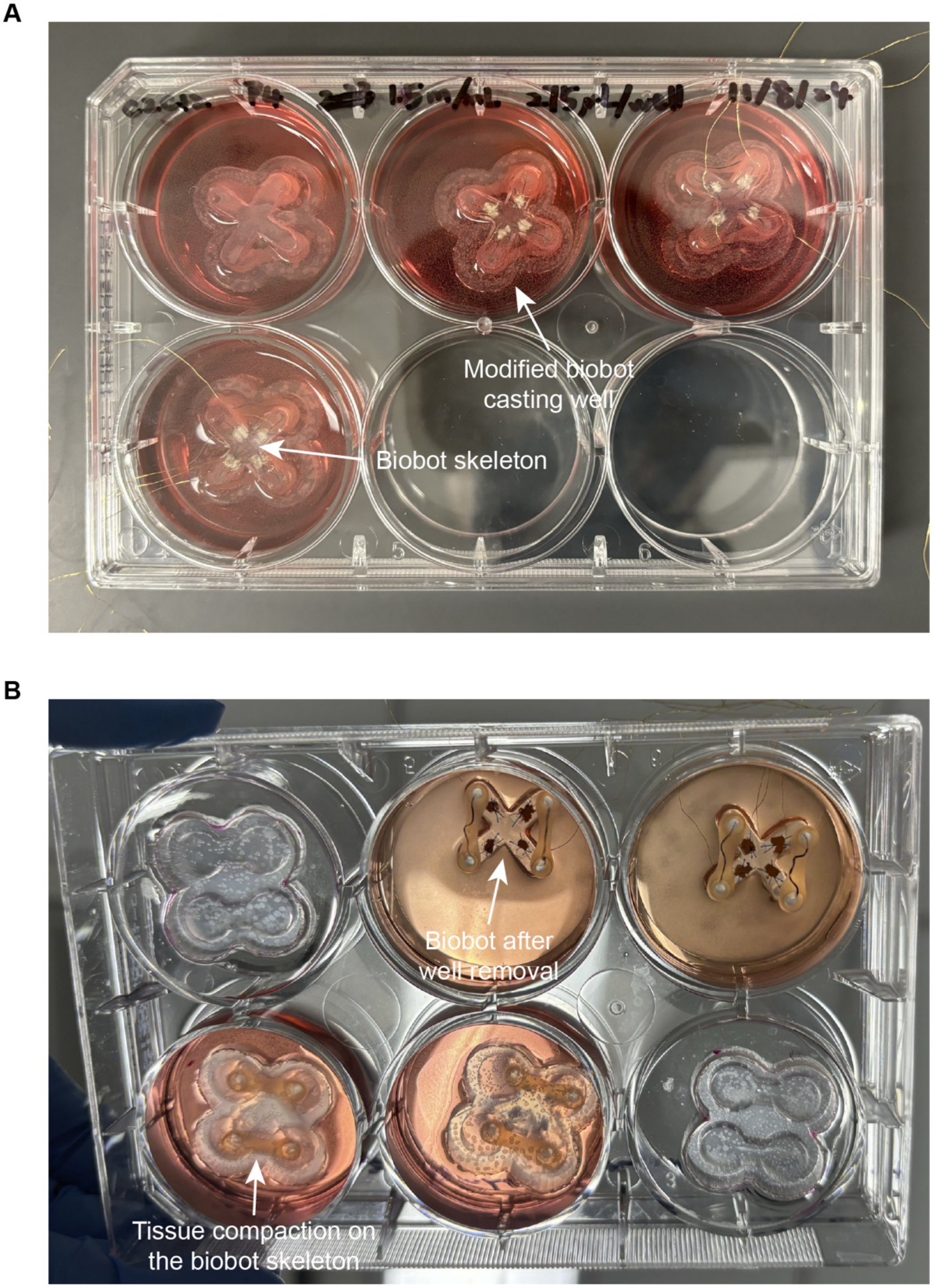


**Fig. S15.**
**Fabrication and release of the biohybrid robotic system.** (**A**) Optical photograph of the biobot muscle casting procedure, showing the precise alignment of the four-post skeleton within the cell-laden hydrogel mixture. (**B**) Optical photograph characterizing tissue compaction. The significant compaction of the engineered muscle facilitates the spontaneous release of the biobot from the dual figure-eight casting well, resulting in a free-standing, autonomous robotic unit ready for functional locomotion trials.

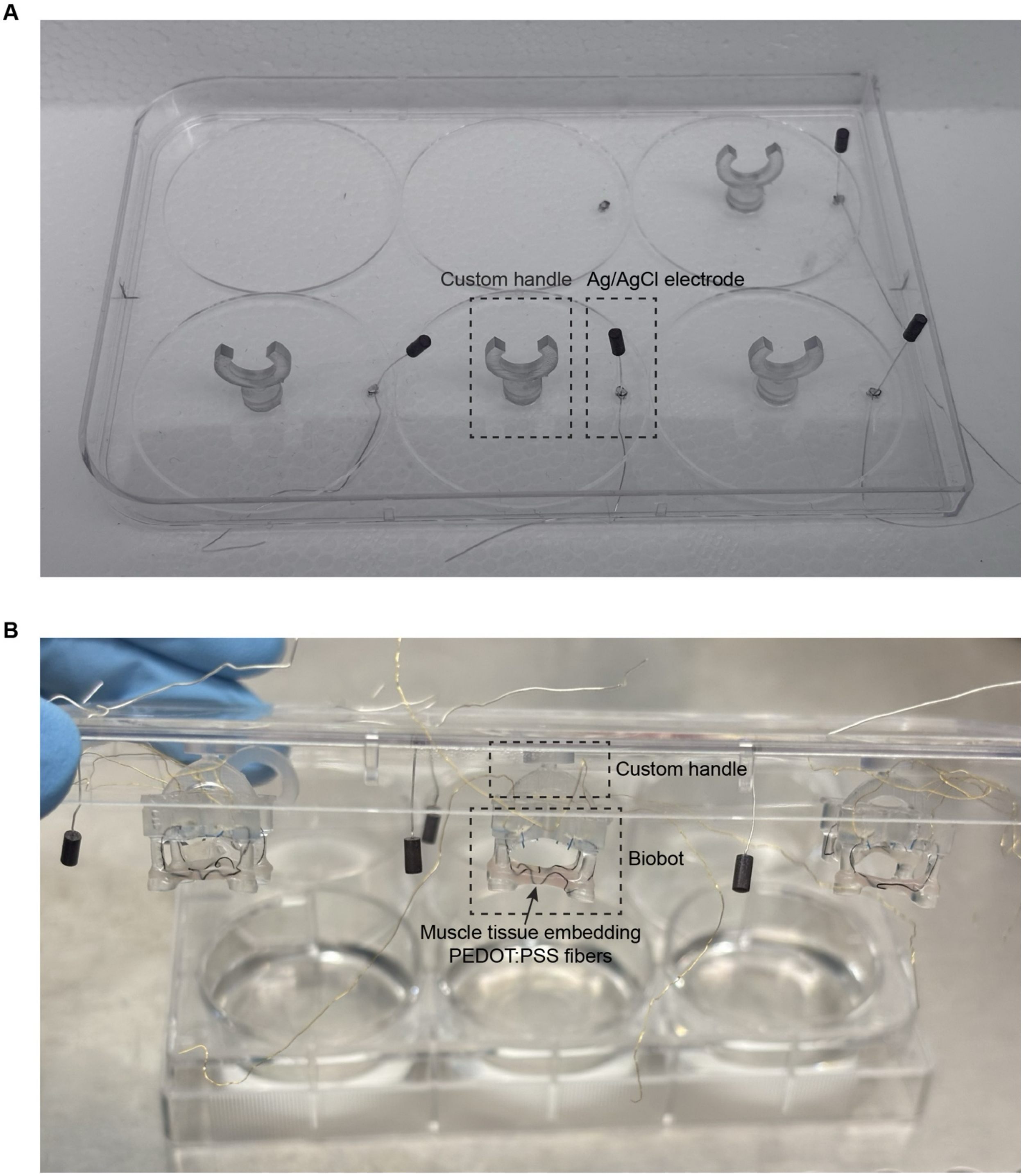


**Fig. S16.**

**Custom handling interface for biobot maintenance and functional testing.** (**A**) Optical photograph of custom-engineered handles mounted to a standard 6-well plate lid. This specialized handling system minimizes mechanical stress on the matured muscle tissues to ensure structural integrity during long-term maintenance, while simultaneously stabilizing the biobot skeleton to prevent unwanted displacement during selective actuation trials. (**B**) Optical photograph demonstrating the biobots in a suspended configuration, secured by the custom handles.

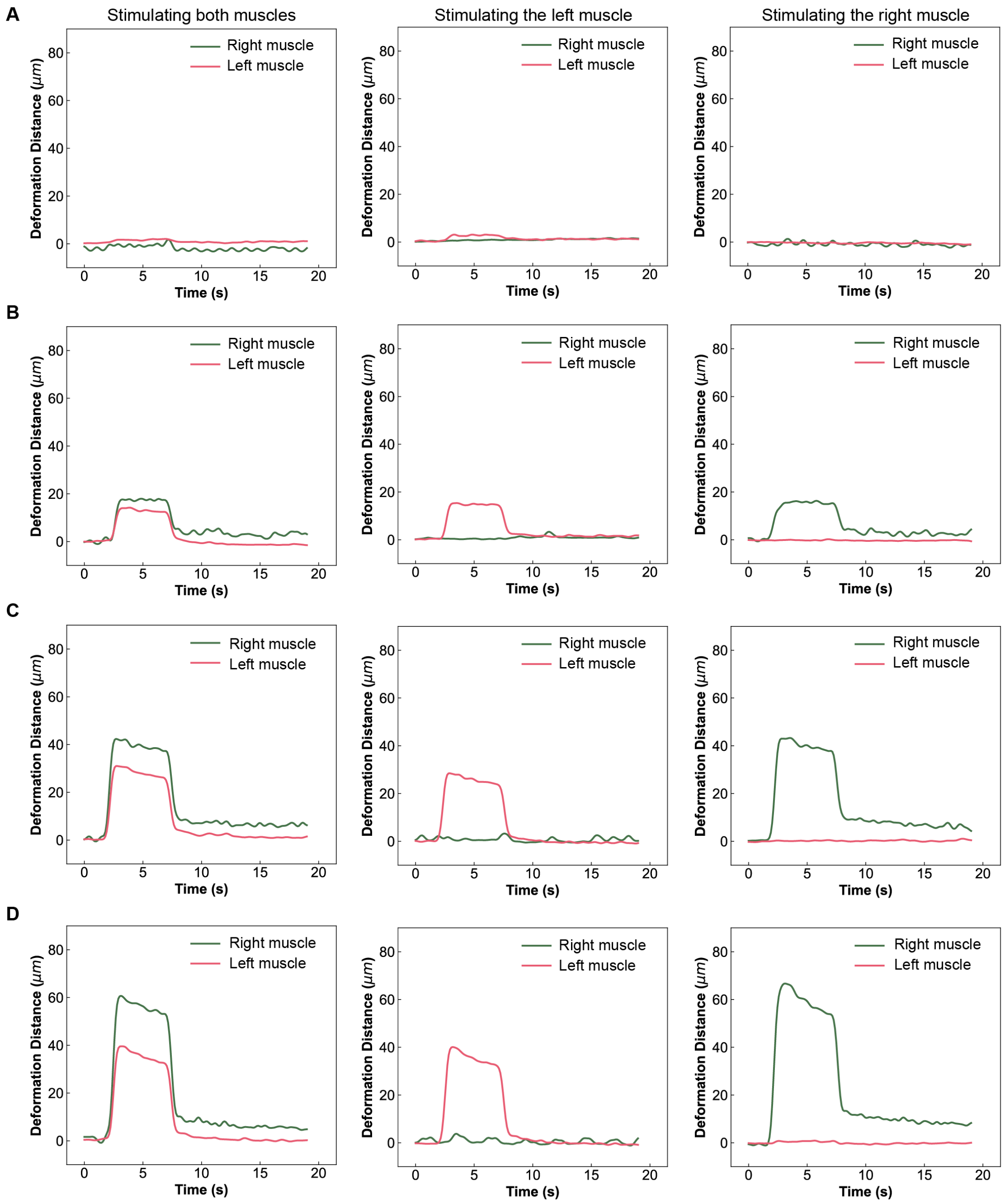


**Fig. S17.**

**Selective actuation and independent control of individual muscle units in dual-muscle biobots.** (**A-D**) Representative characterization of the selective actuation capabilities using individual PEDOT fiber interfaces as a function of applied voltage from 1 V to 4 V. Contraction profiles are displayed for three distinct stimulation modes: simultaneous activation of both muscles, independent activation of the left muscle unit only, and independent activation of the right muscle unit only. All stimulations were conducted at 20 Hz with a 5 ms pulse width for a 5 s duration. This selective control allows for precise individual modulation of each muscle unit, highlighting the utility of the integrated, independent PEDOT fiber architecture for potential complex robotic maneuvers.

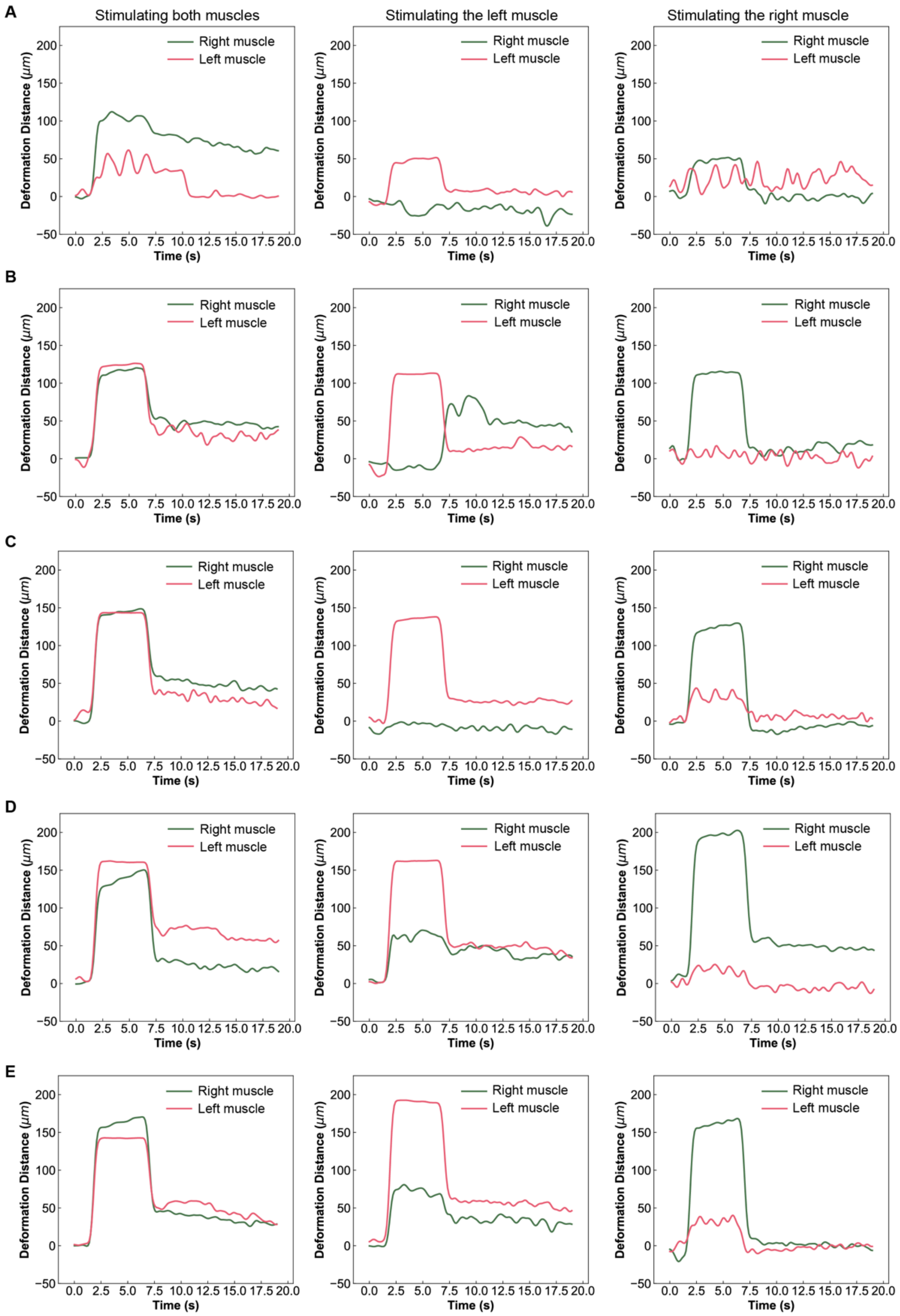


**Fig. S18.**

**Characterization of cross-talk and co-activation thresholds during selective actuation.** (**A-E**) Representative contraction profiles demonstrating varying levels of muscle co-activation as a function of applied voltage (1–5 V). Profiles compare simultaneous activation of both muscle units against independent left-only and right-only stimulation modes (20 Hz, 5 ms pulse width, 5 s duration). These data define the voltage-dependent operational window for achieving high-fidelity independent control versus synchronized dual-actuator recruitment.

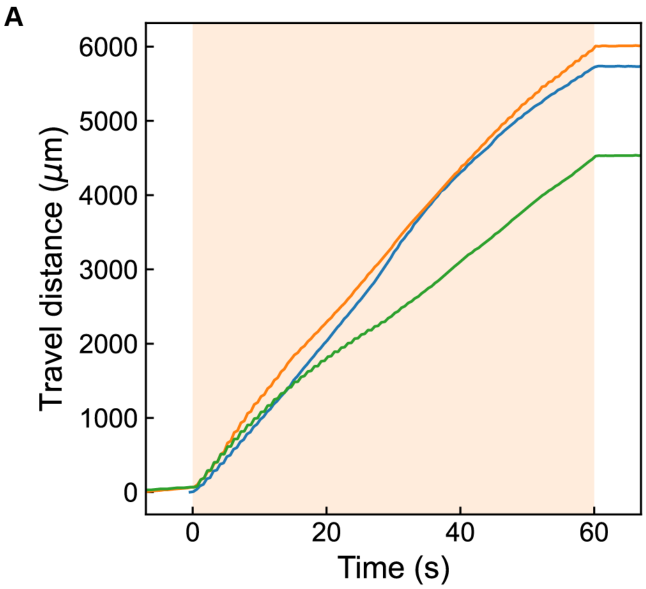


**Fig. S19.**

**Kinetic analysis of biobot locomotion under alternative 1 Hz waveform stimulation.** (**A**) Quantitative tracking of biobot travel distance over time during a 60 s interval of alternative 1 Hz waveform stimulation (fig. S20). The robotic system achieved a steady-state speed of 5.43 ± 0.79 mm/min (mean ± std, N = 3 independent trials). These data demonstrate the reliability of the bio-actuated locomotion and the ability to maintain consistent gait velocity under specific low-frequency electrical pacing.

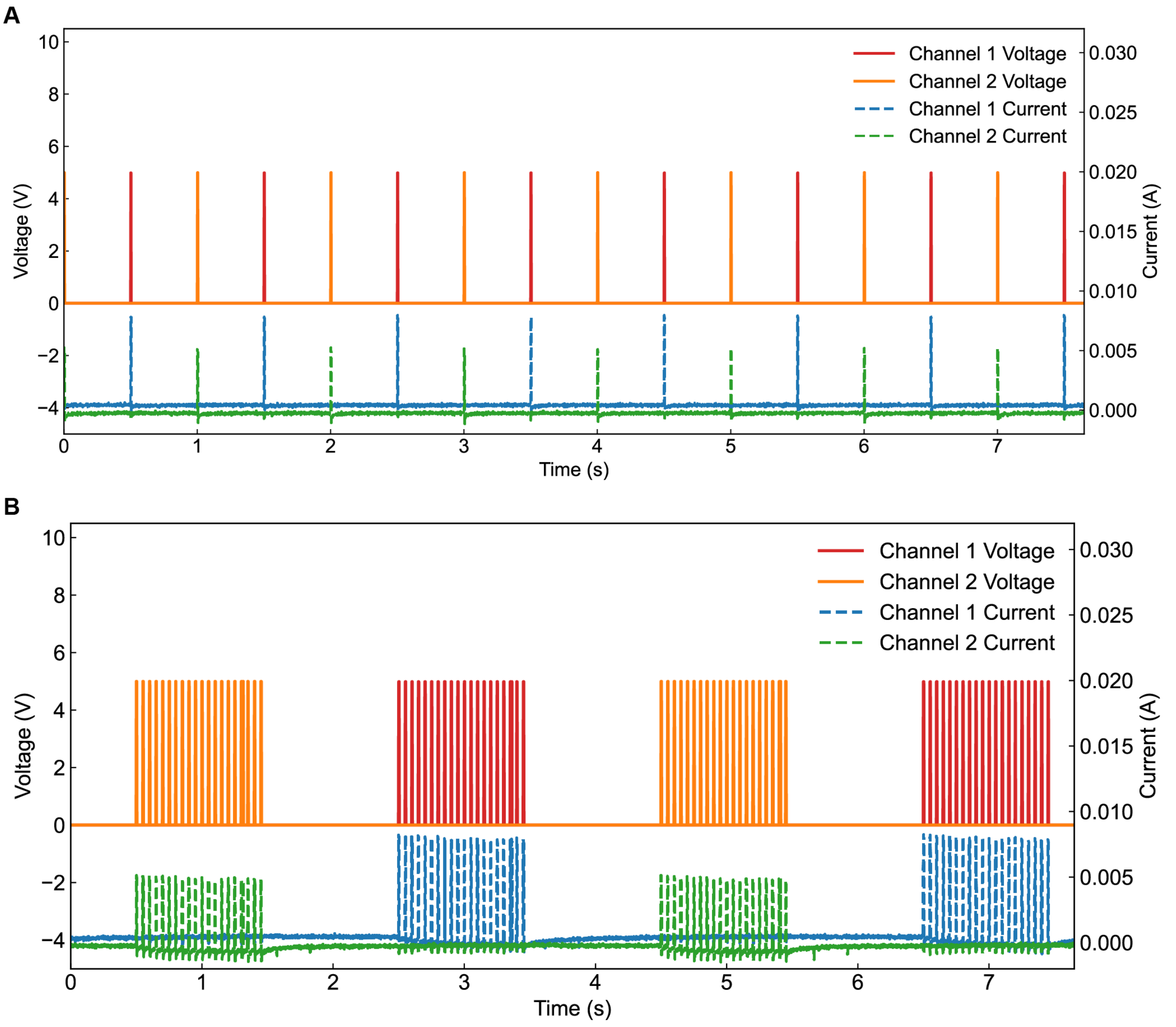


**Fig. S20.**

**Two different alternative stimulation protocols and current output.** (**A**–**B**) Characterization of two distinct alternative stimulation protocols and their corresponding recorded currents. Both protocols were executed at a 5 V amplitude with a 5 ms pulse width. (**A**) A 1 Hz waveform stimulation per channel, resulting in a 2 Hz aggregate pacing frequency for the dual-muscle biobot. (**B**) A 20 Hz waveform stimulation per channel, utilizing a 0.5 Hz aggregate pacing frequency for the robotic system. These protocols demonstrate the versatility of the control architecture in modulating both the individual muscle recruitment and the overall gait frequency of the biobot.

A

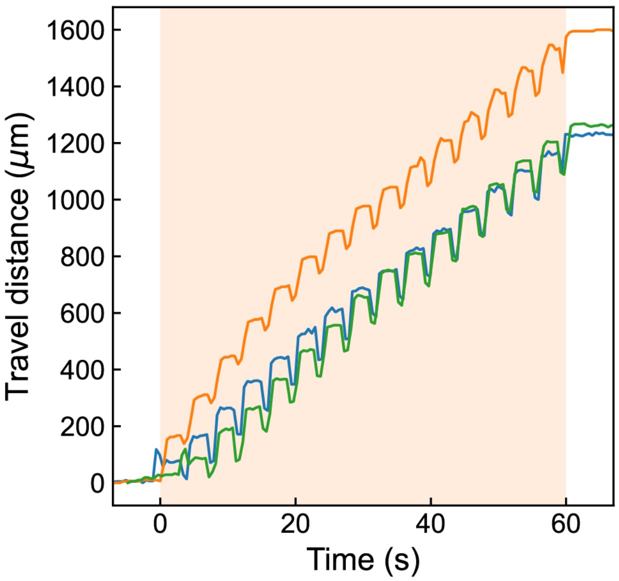


**Fig. S21.**

**Kinetic analysis of biobot locomotion under alternative 20 Hz waveform stimulation.** (**A**) Quantitative tracking of biobot travel distance over time during a 60 s interval of alternative 20 Hz waveform stimulation (fig. S20). The robotic system achieved a steady-state translational speed of 1.37 ± 0.20 mm/min (mean ± std, N = 3 independent trials). When contrasted with the pacing results in Fig. S19, these data highlight the ability to precisely tune the locomotion velocity of the biobot by adjusting the underlying gait frequency and waveform characteristics.

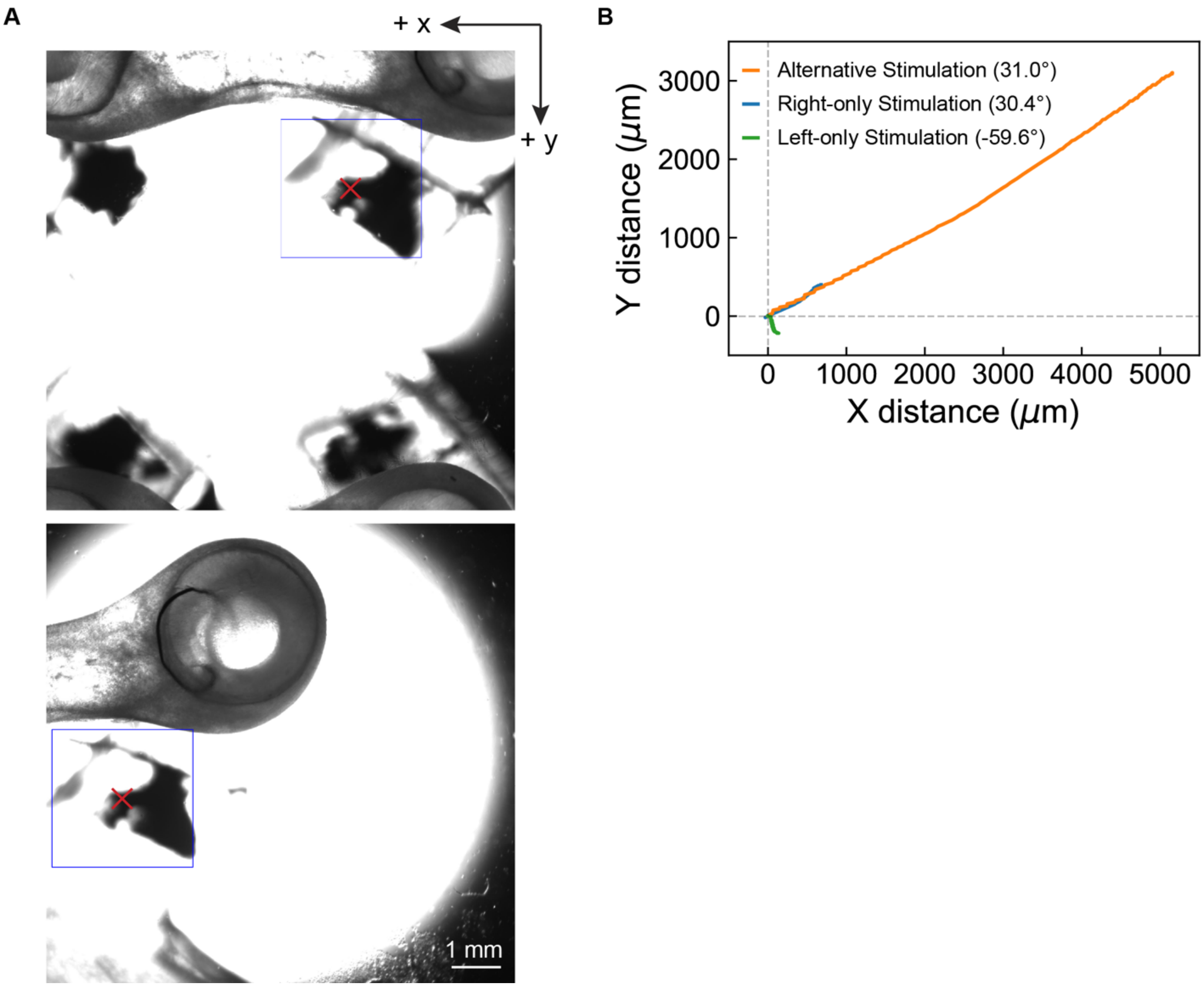


**Fig. S22.**

**Kinematic tracking and directional trajectory modulation of biohybrid locomotion.** (**A**) Representative Region of Interest (ROI) capturing a specific feature at the initial and final positions of the biobot during a locomotion trial. Automated feature tracking characterizes the movement and orientation changes over the duration of the test. Scale bar, 1 mm. (**B**) Characterization of trajectory angle modification via asymmetrical muscle recruitment. Traces illustrate the walking paths during 1 Hz waveform stimulation (at 5 V with 5 ms pulse width for a 60 s duration) under bilateral (both muscles) or unilateral (single-muscle) actuation. These trajectories demonstrate the steering capabilities of the integrated PEDOT fiber architecture, enabling controlled changes in the biobot's heading through selective activation of individual muscle units.

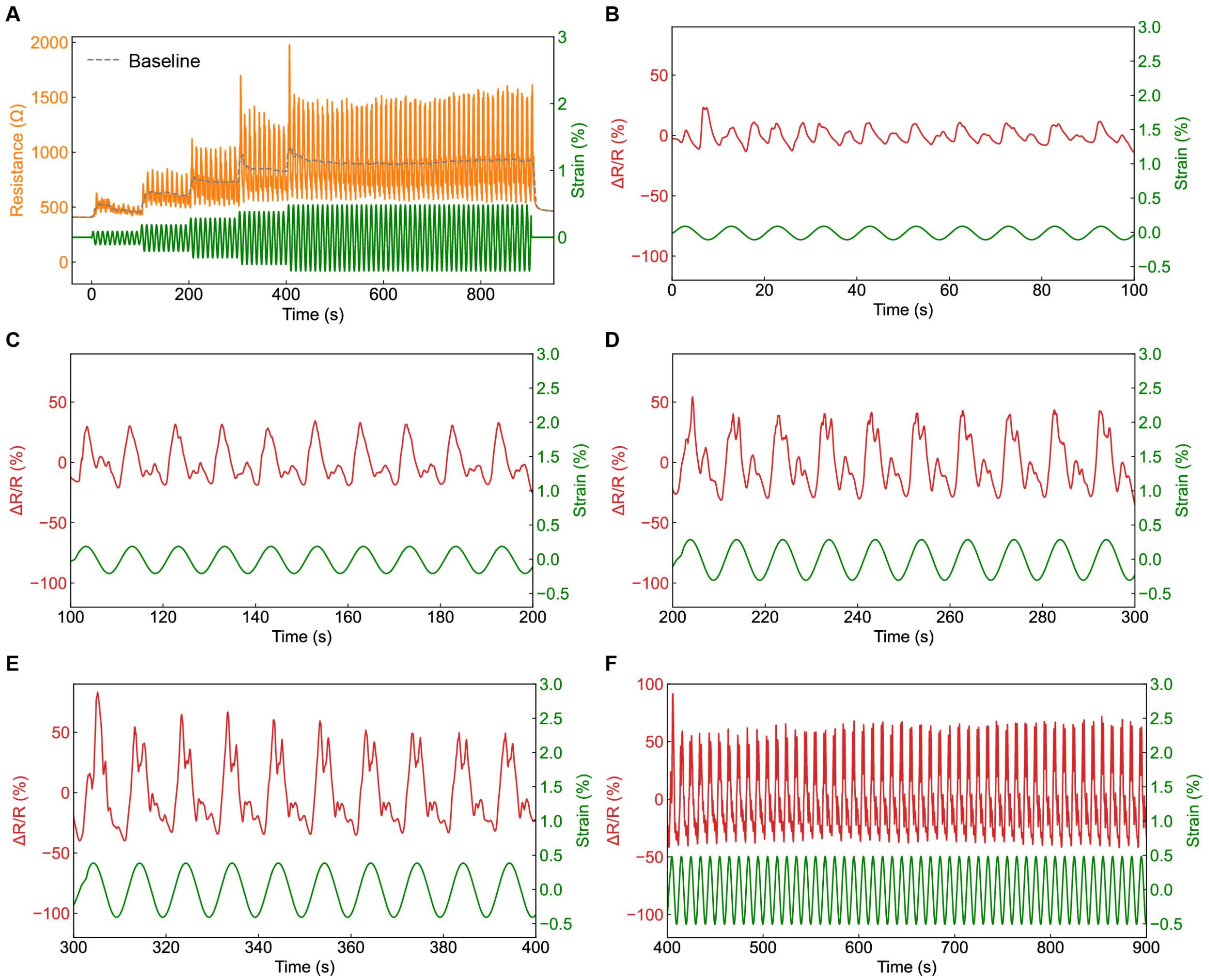


**Fig. S23.**
**Electromechanical characterization and stability of PEDOT fibers.** (**A**) Raw resistance profiles (yellow) of a standalone PEDOT fiber under cyclic sinusoidal loading (green), displayed without baseline correction. The observed baseline drift is attributed to irreversible micro-crack propagation and polymer chain realignment during the initial loading cycles at increasing strain amplitudes. (**B**-**F**) Relative resistance change (ΔR/R, red) under cyclic sinusoidal strains (green) ranging from 0.1% to 0.5%. These data characterize the fiber's sensitivity and signal stability within the physiological strain regime of engineered muscle tissues, highlighting its utility as a high-resolution, integrated strain sensor.

**A**

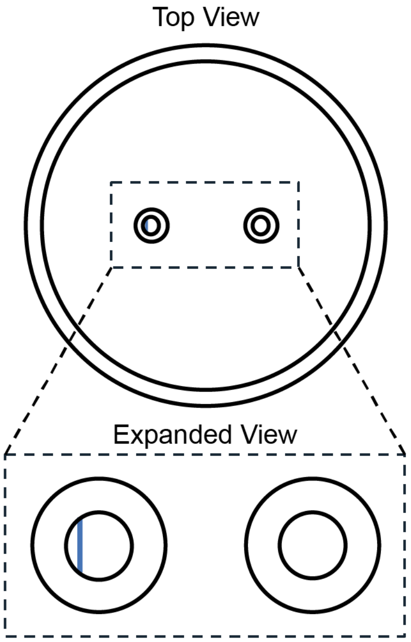


**Fig. S24.**
**Spatial optimization of the integrated sensing architecture within the post structure.** (**A**) Schematic top-view illustrating the precise placement of the U-shaped PEDOT fiber within the elastomeric post. The sensing plane (indicated by the blue line) is strategically offset from the geometric center (neutral axis) of the device. This off-centered positioning serves to maximize strain sensitivity during muscle-induced post-deflection, ensuring high-fidelity transduction of contractile force into resistance change.

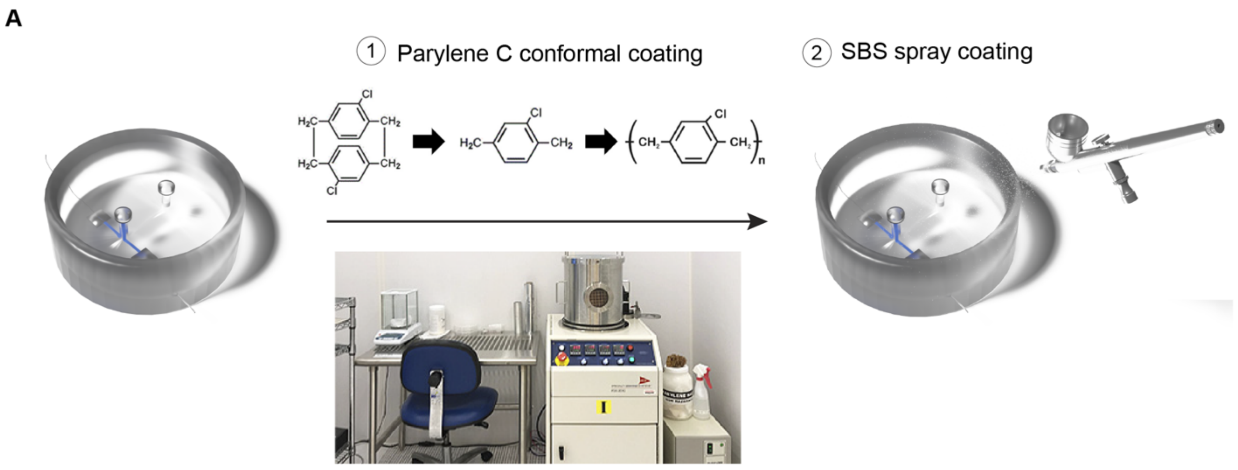


**Fig. S25.**

**Encapsulation workflow for integrated strain-sensing interfaces.** (**A**) Schematic illustration of the multi-step encapsulation process designed to provide electrical insulation and moisture protection for the PEDOT-based sensing units. The photographic inset depicts the chemical vapor deposition (CVD) system utilized for Parylene-C coating (image adapted from the Northwestern University Atomic and Nanoscale Characterization Experiment Center, NUFAB). This encapsulation layer ensures stable sensor performance in liquid environments by protecting the electrical interfacing and preventing ion-induced electrochemical noise.

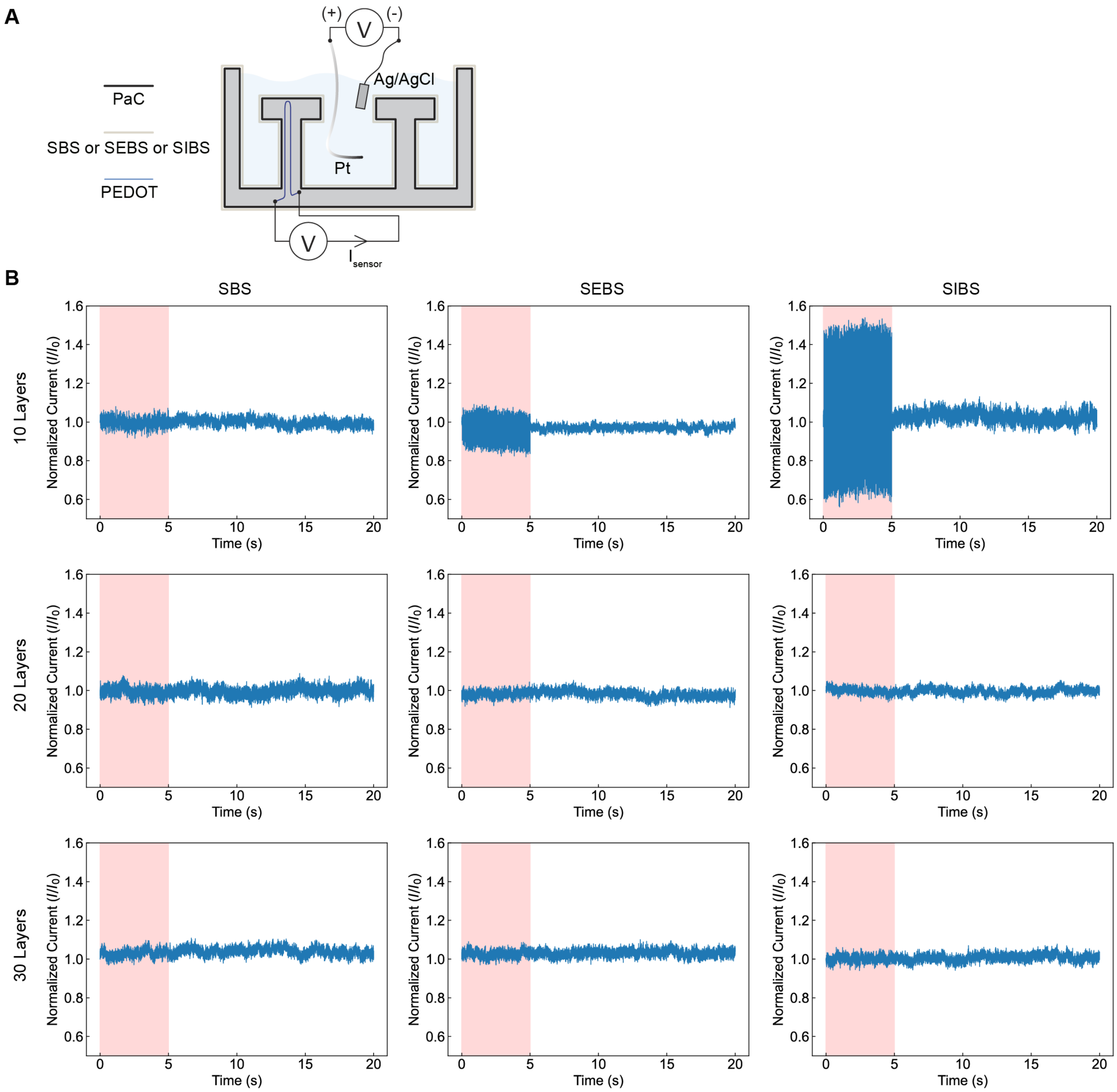


**Fig. S26.**

**Optimization of elastomeric encapsulation for electrical signal isolation.** (**A**) Schematic of the experimental setup utilized to test stimulation-induced interference. Strain sensors were evaluated in the absence of muscle tissue to characterize the baseline electrical noise during stimulation (5 V, 20 Hz, 5 ms pulse width, 5 s duration). (**B**) Quantitative comparison of stimulation interference across three candidate elastomers: SBS, SEBS, and SIBS, applied at varied thicknesses (10, 20, and 30 spray-coated layers). Among the 10-layer configurations, SBS exhibited the superior signal protection, demonstrating the highest efficacy in suppressing stimulation-induced artifacts.

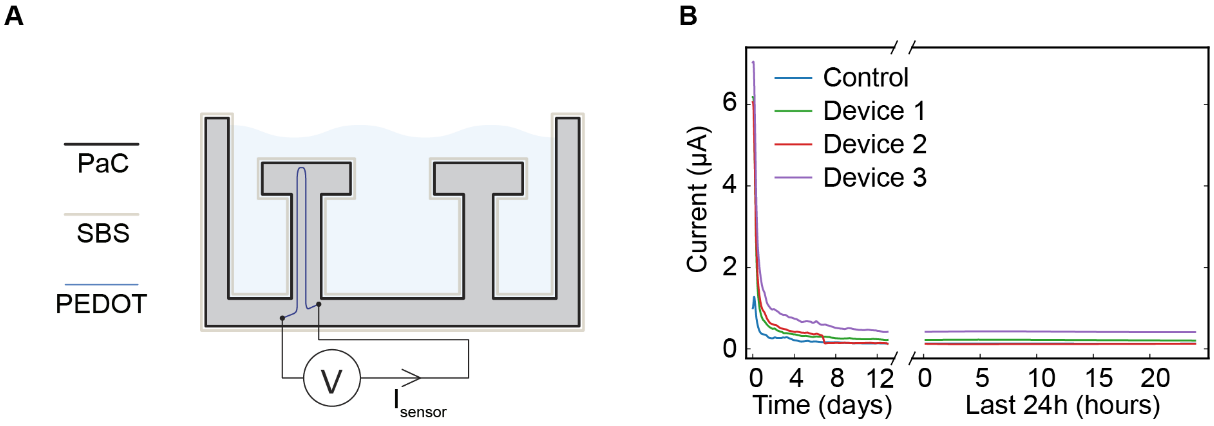


**Fig. S27.**
**Long-term operational stability and encapsulation efficacy.** (**A**) Schematic of the experimental setup for the 14-day longitudinal stability trial. (**B**) Long-term electrochemical stability comparison between uncoated (control) and encapsulated (coated, N = 3) sensing devices during continuous immersion in PBS. In the late phase of immersion, both the control and encapsulated devices maintain stable baseline performance, whereas the uncoated control devices exhibit slightly more rapid degradation during early immersion phase. Notably, the abrupt decrease in the current profile for Device 2 around Day 7 implies the onset of its encapsulation failure, characterizing the temporal limits of the current protective barrier.

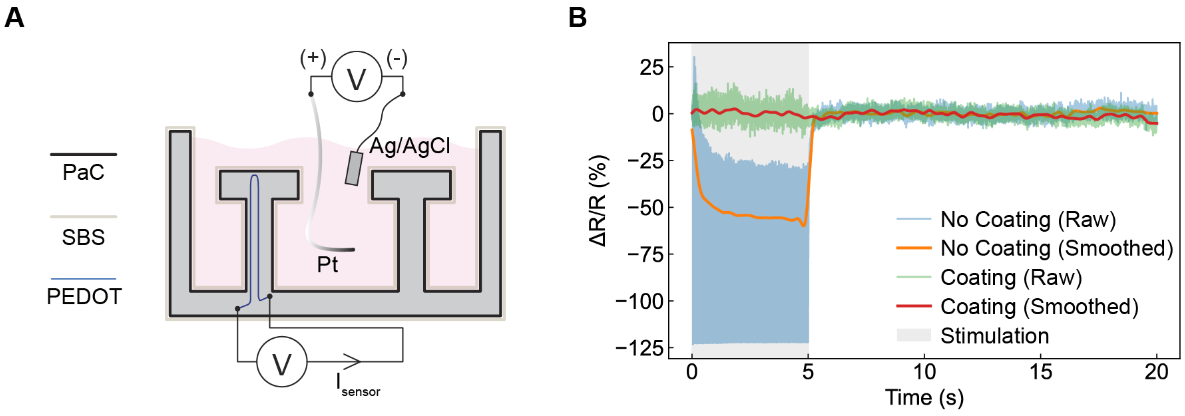


**Fig. S28.**

**Evaluation of sensing signal integrity following long-term environmental conditioning.** (**A**) Experimental setup for assessing stimulation-induced interference on encapsulated versus uncoated sensing devices. Both configurations were evaluated after a 10-day pre-soak in tissue culture medium at 37 °C to simulate physiological conditions in the absence of muscle tissue. (**B**) A characteristic example demonstrating successful signal protection provided by the multilayer coatings during active stimulation. The sensing signal remains stable and free of significant artifacts during the stimulation interval (5 V, 20 Hz, 5 ms pulse width, for 5 s duration). These results confirm that the multilayer encapsulation architecture maintains high electrical isolation and moisture barrier properties under chronic physiological immersion.

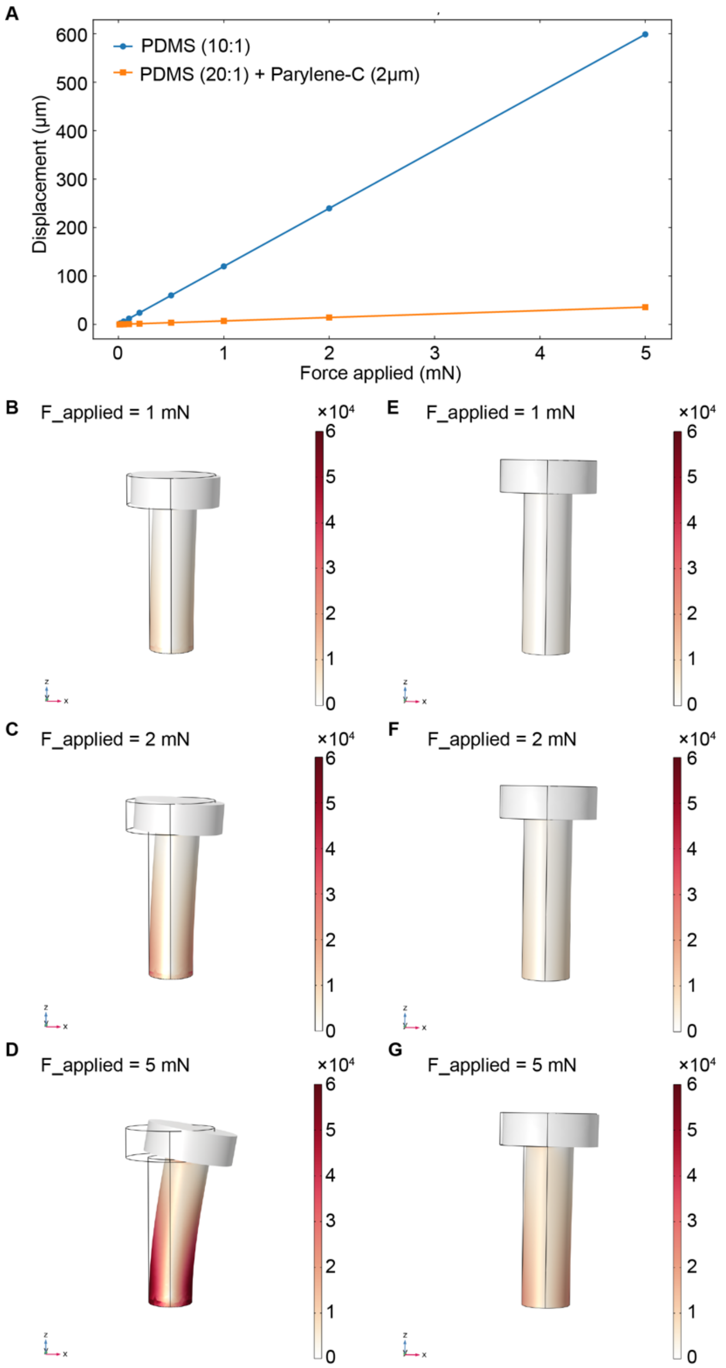


**Fig. S29.**

**Finite element analysis (FEA) of mechanical stiffness and stress distribution in uncoated and encapsulated pillars.** (**A**) Computational simulation characterizing the displacement-force relationship for uncoated (stimulation device) and encapsulated (sensing device) PDMS pillars. The linear response profiles quantify the influence of the protective coatings on the structural stiffness, characterizing the reduced displacements expected during muscle contraction. (**B-D**) Von Mises stress distribution of the uncoated pillar under applied point loads of 1 mN, 2 mN, and 5 mN, respectively. (**E-G**) Stress distribution profiles for the encapsulated pillar under identical loading conditions (1 mN, 2 mN, and 5 mN).

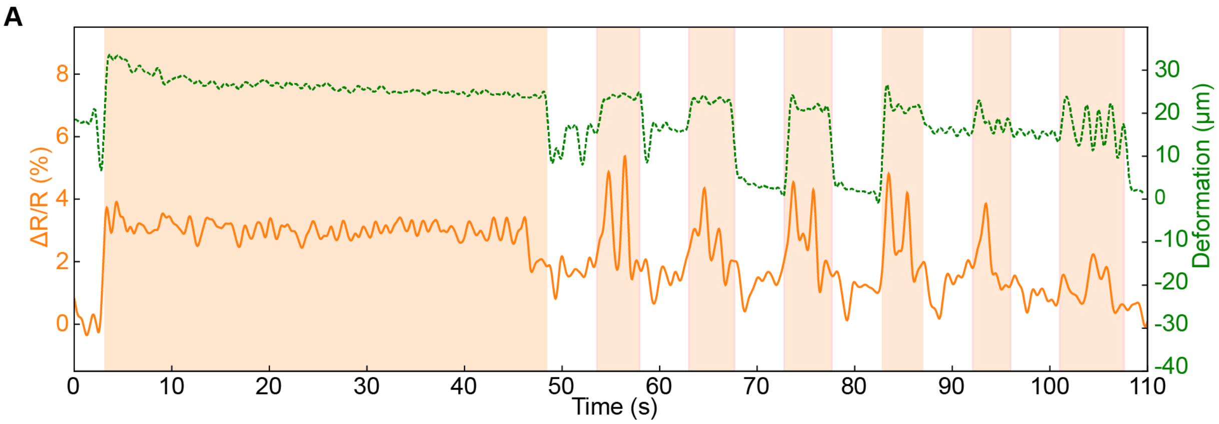


**Fig. S30.**

**Real-time sensor feedback and synchronized validation of the closed-loop control system.** **(A)** Representative raw data traces illustrating the performance of the integrated closed-loop control system driven by real-time sensor feedback. Both the sensor resistance signal and the corresponding microscopic deformation data were processed using a low-pass filter (1 Hz cut-off frequency). Light-orange shaded regions denote active stimulation intervals. These synchronized datasets demonstrate the high-fidelity correlation between the PEDOT sensor output and physical displacement, validating the system's ability to autonomously modulate the bio-actuator activity based on the current functional state and fatigue levels.

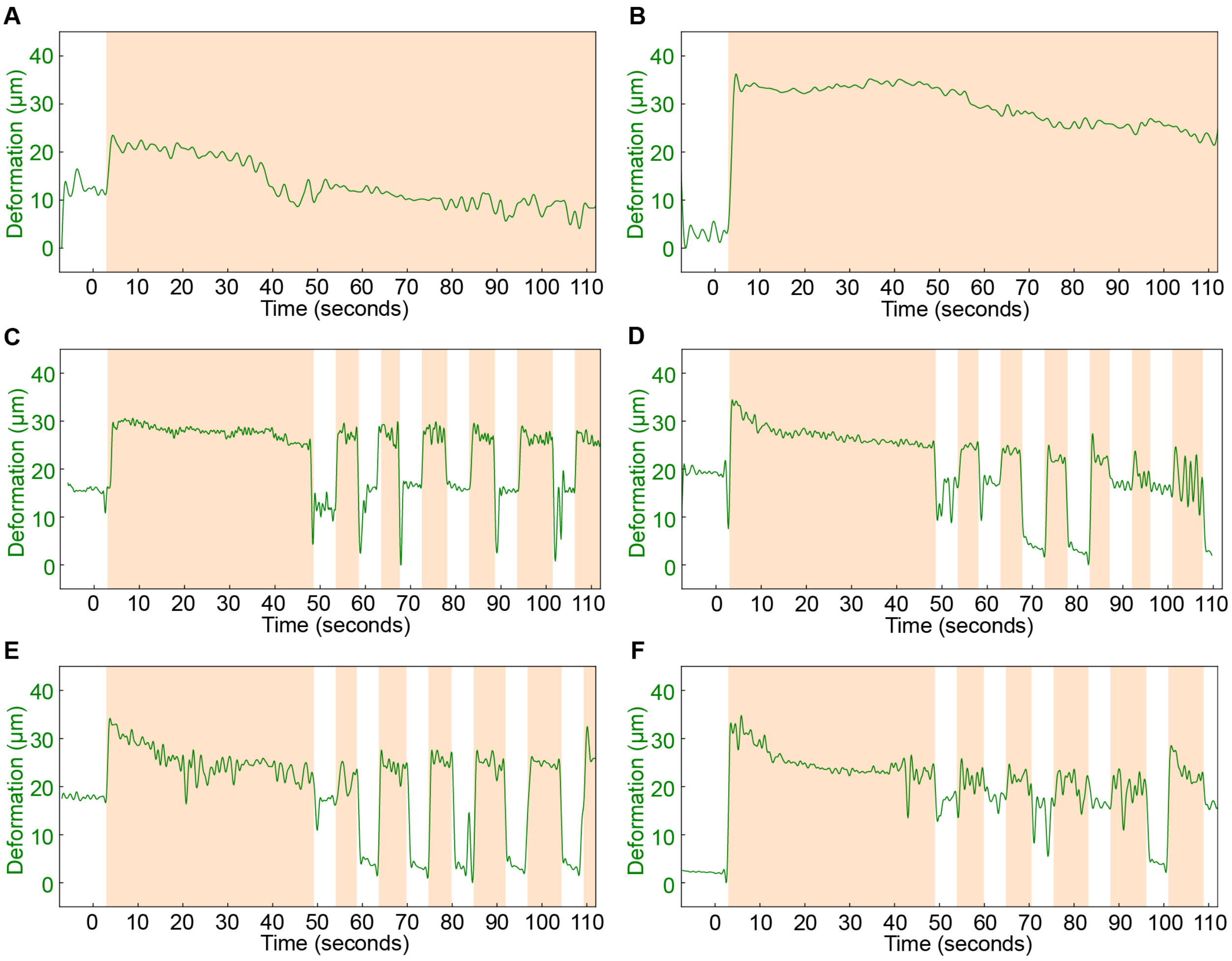


**Fig. S31.**

**Comparative performance and fatigue analysis under open-loop and adaptive closed-loop control.** (**A-B**) Representative muscle contraction profiles during continuous, non-stop stimulation under an open-loop control regime (N = 2 independent studies). (**C-F**) Representative contraction profiles under the adaptive closed-loop control strategy (N = 4 independent studies), where stimulation is modulated based on real-time sensor feedback. Light-orange shaded regions indicate active stimulation intervals. These results demonstrate the efficacy of the integrated PEDOT strain sensor in enabling a feedback-driven architecture that effectively mitigates biological fatigue and preserves long-term contractile performance.

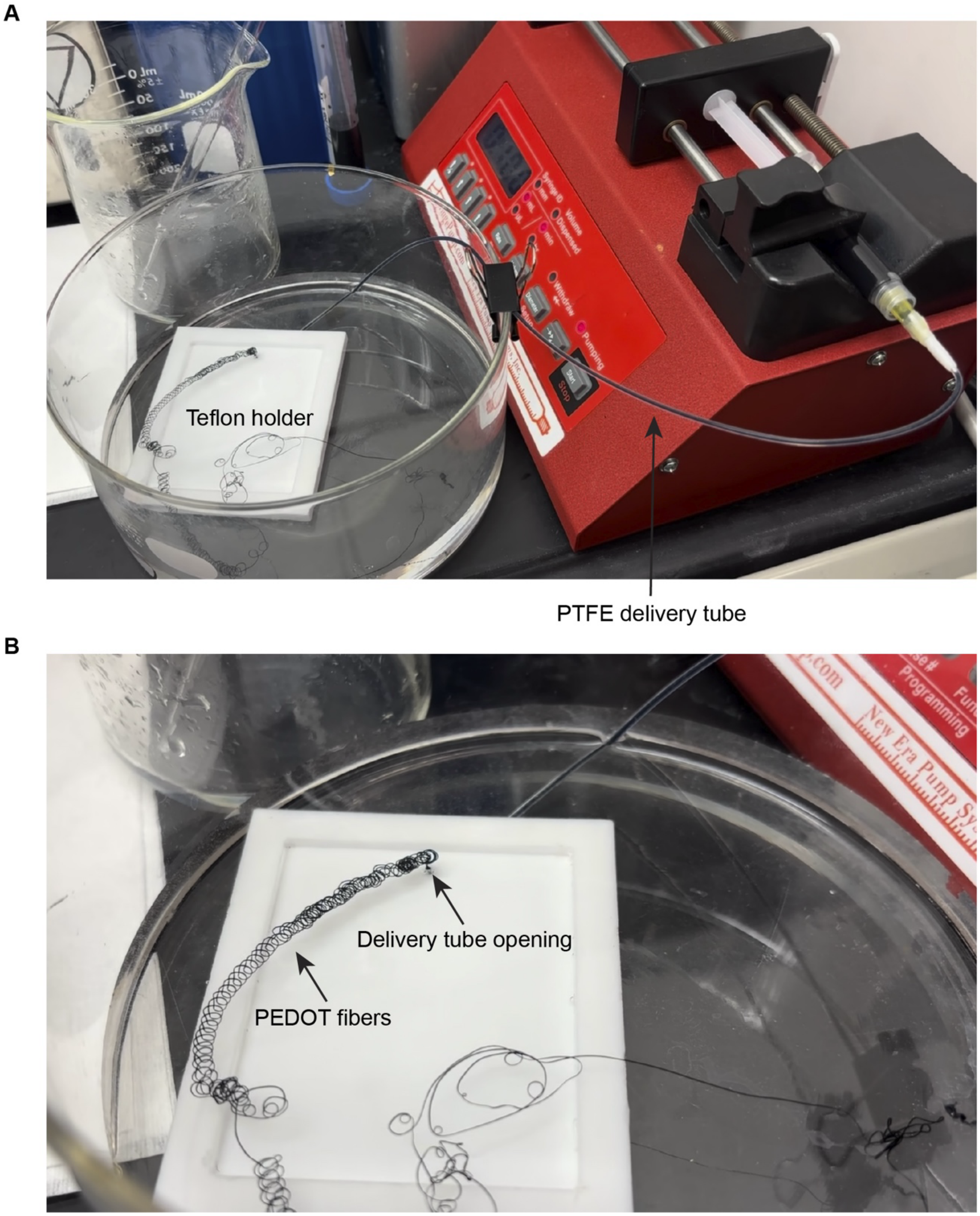


**Fig. S32.**
**Wet-spinning fabrication setup for high-performance PEDOT fibers.** (**A**) Photographic overview of the experimental wet-spinning setup, showing the integrated syringe pump system and the coagulation bath assembly used for fiber synthesis. (**B**) Detailed photographic view of the PEDOT:PSS solution extruded into the coagulation bath vertically. This orientation is optimized to minimize floating distortion and ensure the production of fibers with uniform diameter and consistent electromechanical properties.

**A**

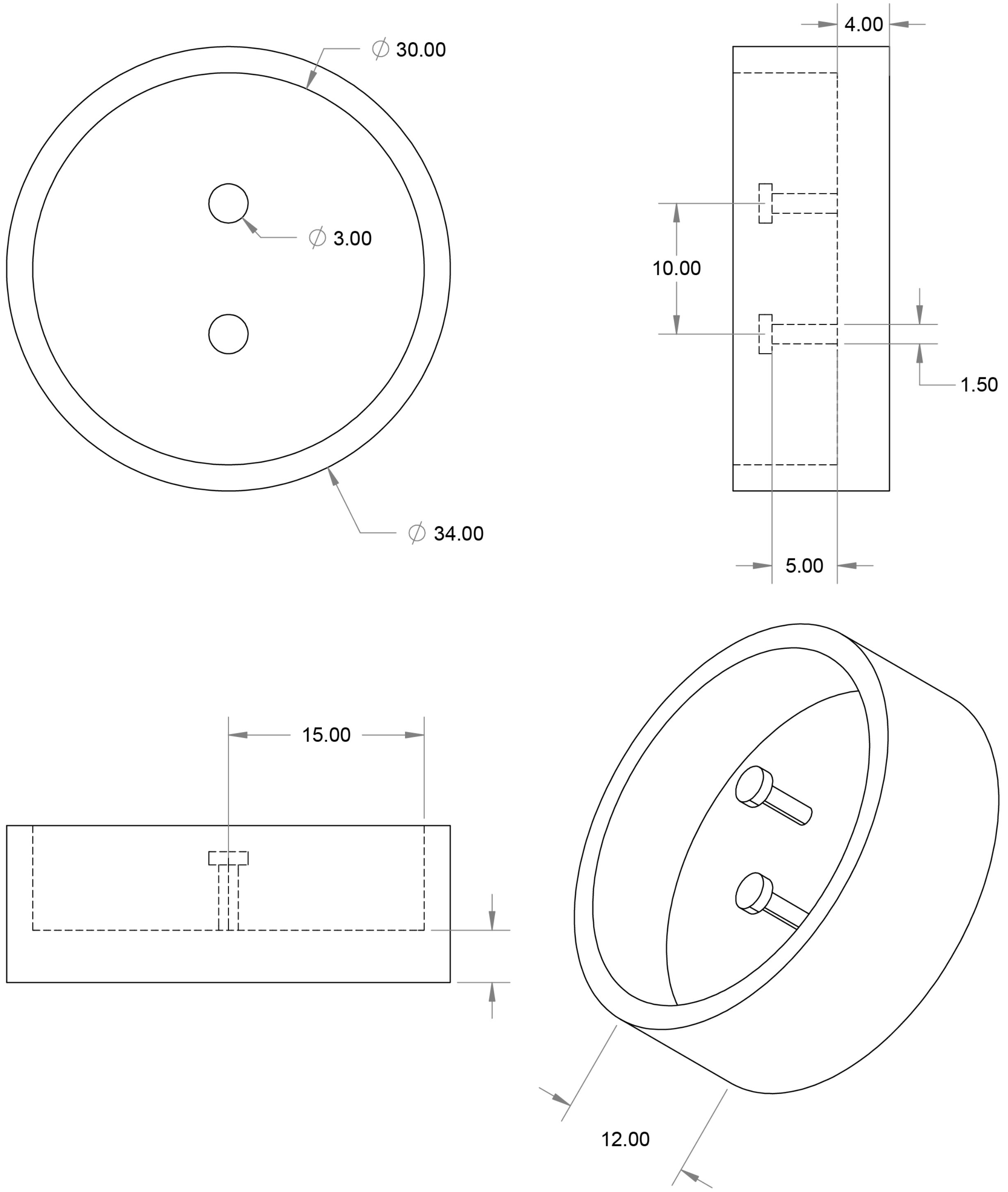


**Fig. S33.**

**Computer-aided design (CAD) and geometric specifications of the biohybrid platforms.** (**A**) Detailed CAD schematics illustrating the structural dimensions (in mm) of the 2-post stimulating and sensing devices. The design specifies critical parameters, including the pillar height, diameter, and inter-post spacing, etc.

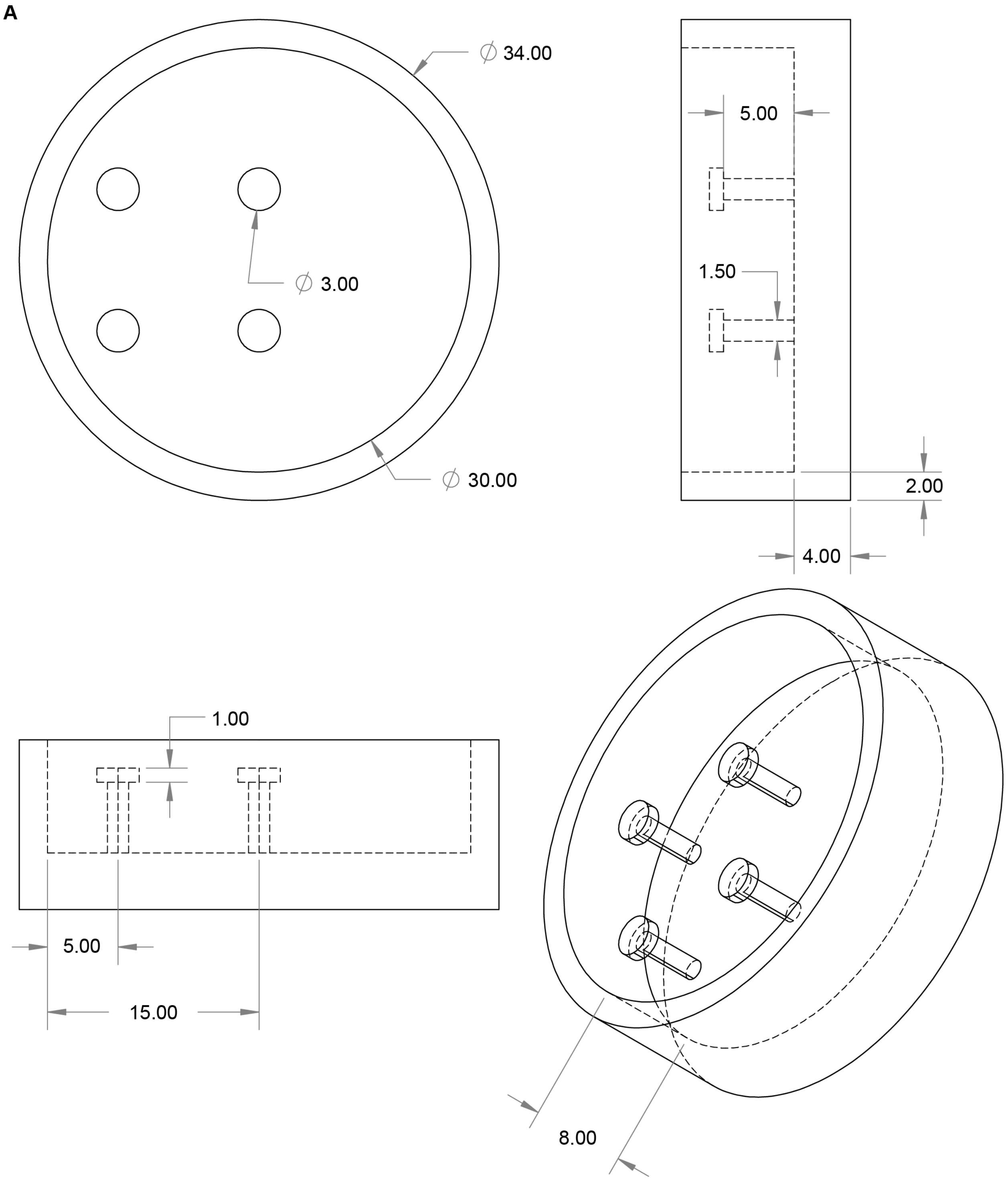


**Fig. S34.**

**CAD and geometric specifications of the 4-post stimulating platform.** (**A**) Dimensions of devices (mm). Detailed CAD schematics illustrating the structural dimensions (in mm) of the 4-post stimulating devices. The design defines the precise spatial arrangement, including pillar diameters, heights, and transverse/longitudinal inter-post spacing, etc.

**A**

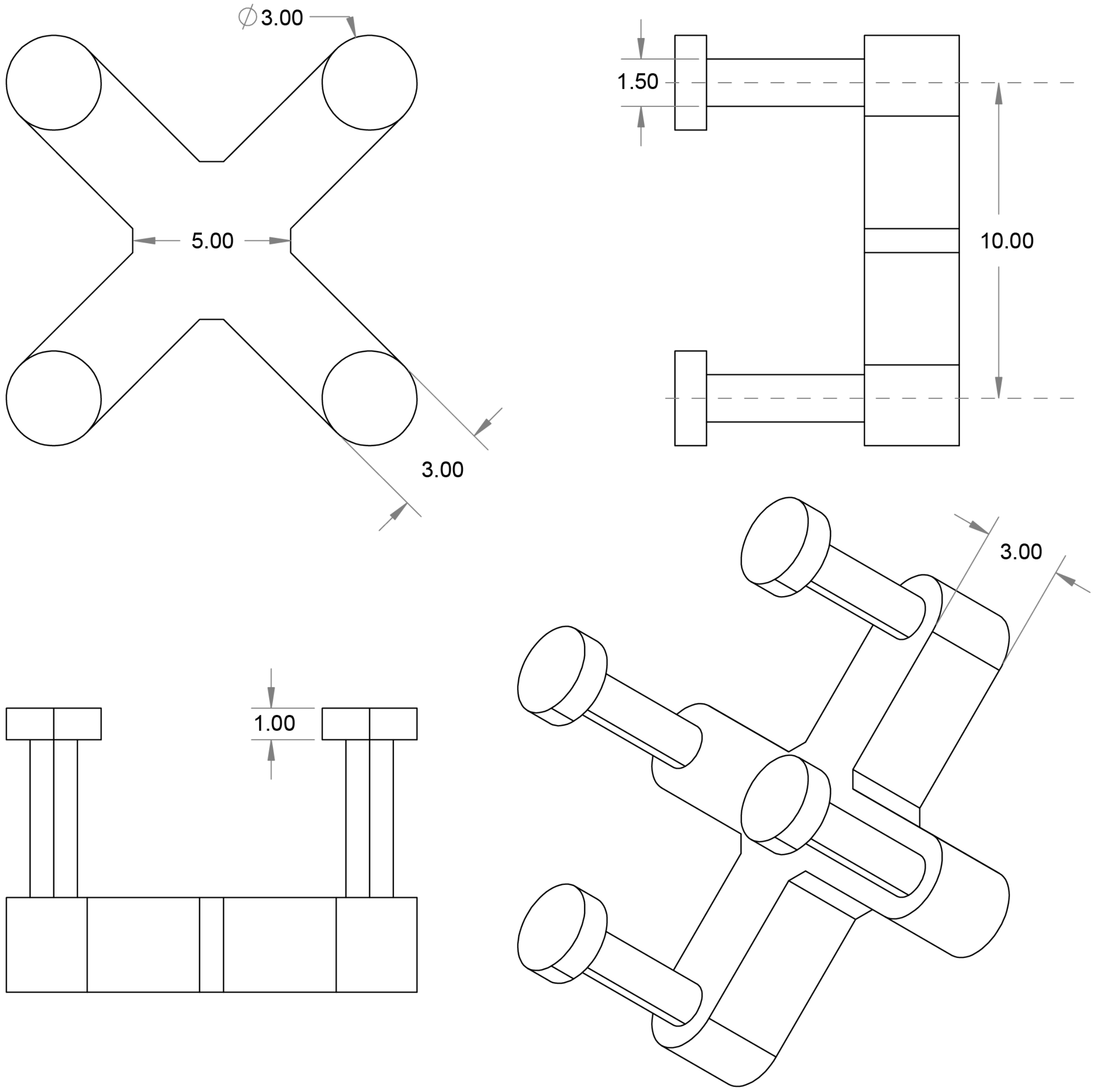


**Fig. S35.**

**CAD and geometric specifications of the 4-post biobot skeletons.** (**A**) Detailed CAD schematics illustrating the structural dimensions (in mm) of the 4-post biobot skeletons. The design defines the global geometry, including the thickness, pillar diameters, heights, and transverse/longitudinal inter-post spacing, etc.

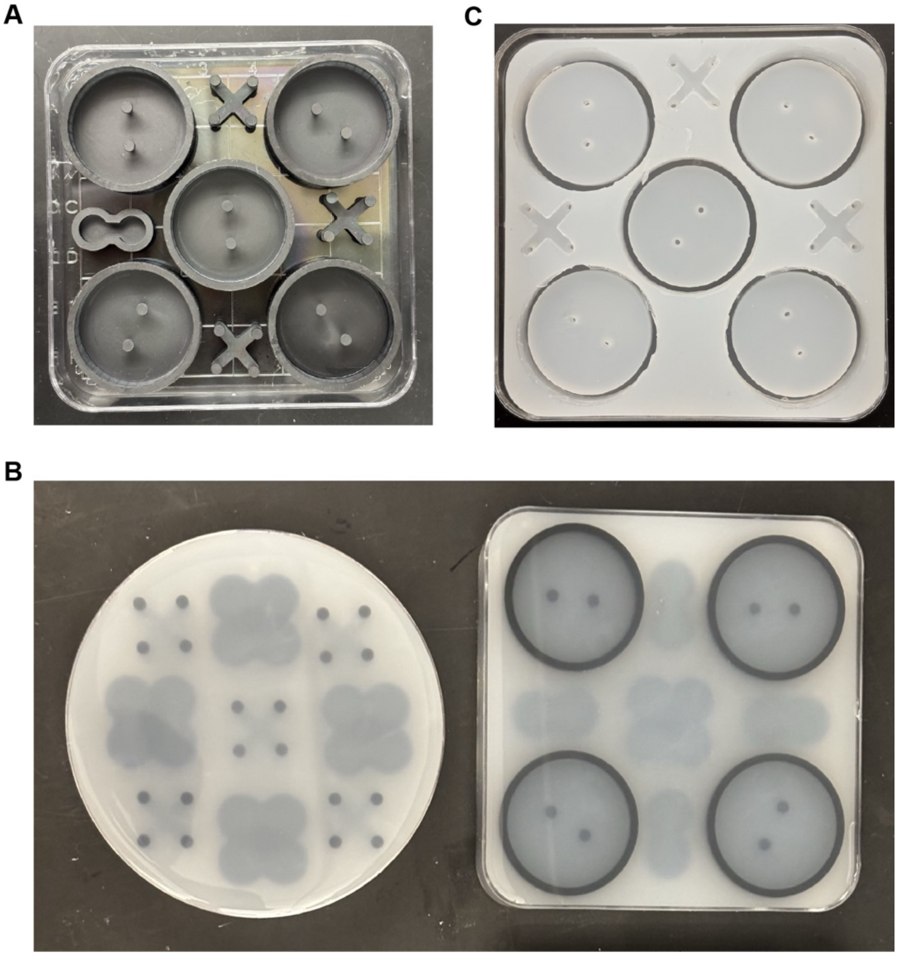

**Fig. S36.**
**Fabrication workflow for soft lithographic Ecoflex negative molds.** (**A**) Photographic overview of the high-resolution 3D-printed resin masters, which define the primary skeletal geometry of the bio-actuators. (**B**) Photographic image depicting the casting and curing of Ecoflex silicone within Petri dishes containing the resin masters. (**C**) Photographic image of the finalized Ecoflex molds following the delamination and release of the resin masters. This mold-based fabrication process enables the rapid and repeatable production of PDMS stimulation/sensing devices and biobot skeletons with high fidelity to the original CAD specifications.

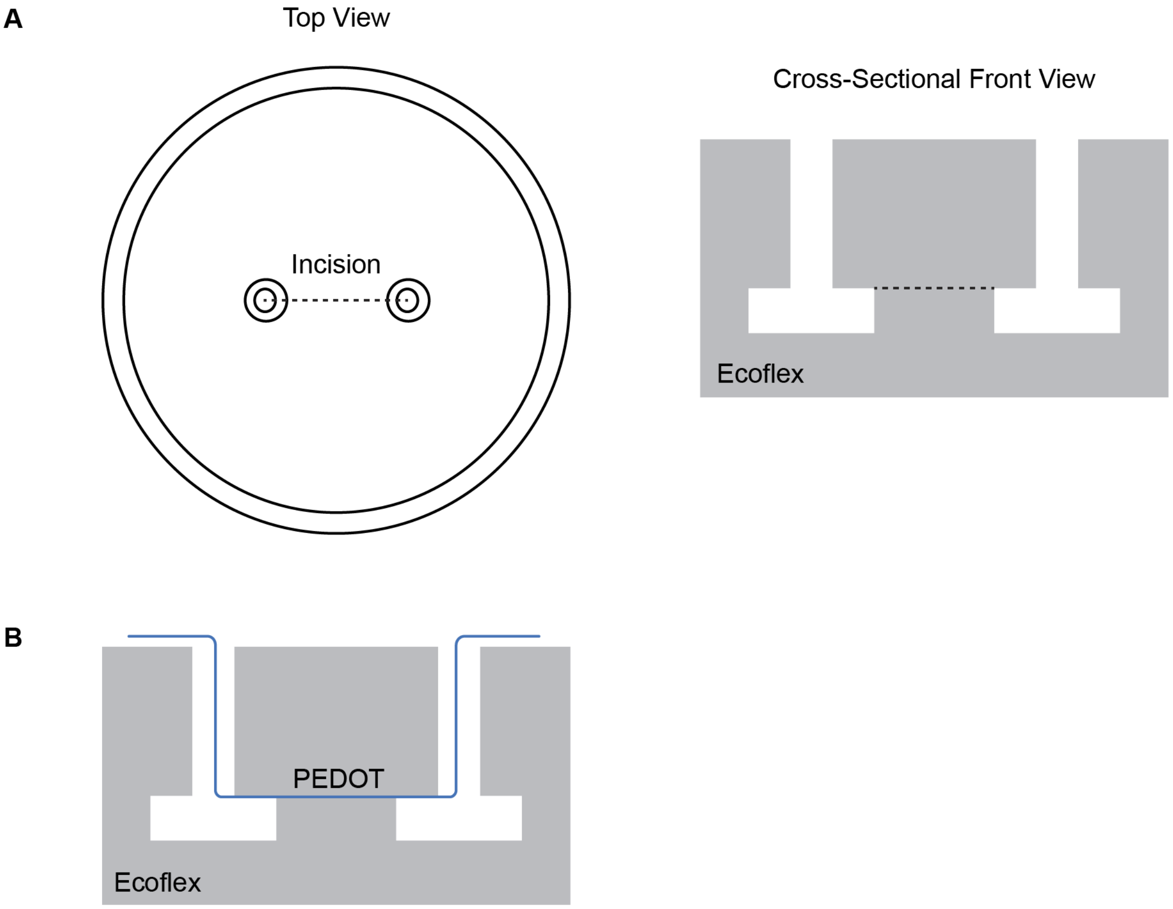


**Fig. S37.**

**Methodological schematics for fiber integration for stimulation devices during the mold-casting process.** (A) Schematic illustration of the precise incision locations within the Ecoflex negative mold. These incisions are strategically placed to facilitate the insertion of fiber electrodes while maintaining the mold's seal during PDMS curing. (B) Schematic depicting the placement and alignment of the PEDOT fibers within the mold cavity. This localized positioning ensures that the fibers are securely embedded at the target coordinates within the pillar structure, ensuring reliable electrical contact and mechanical coupling with the final biohybrid actuator.

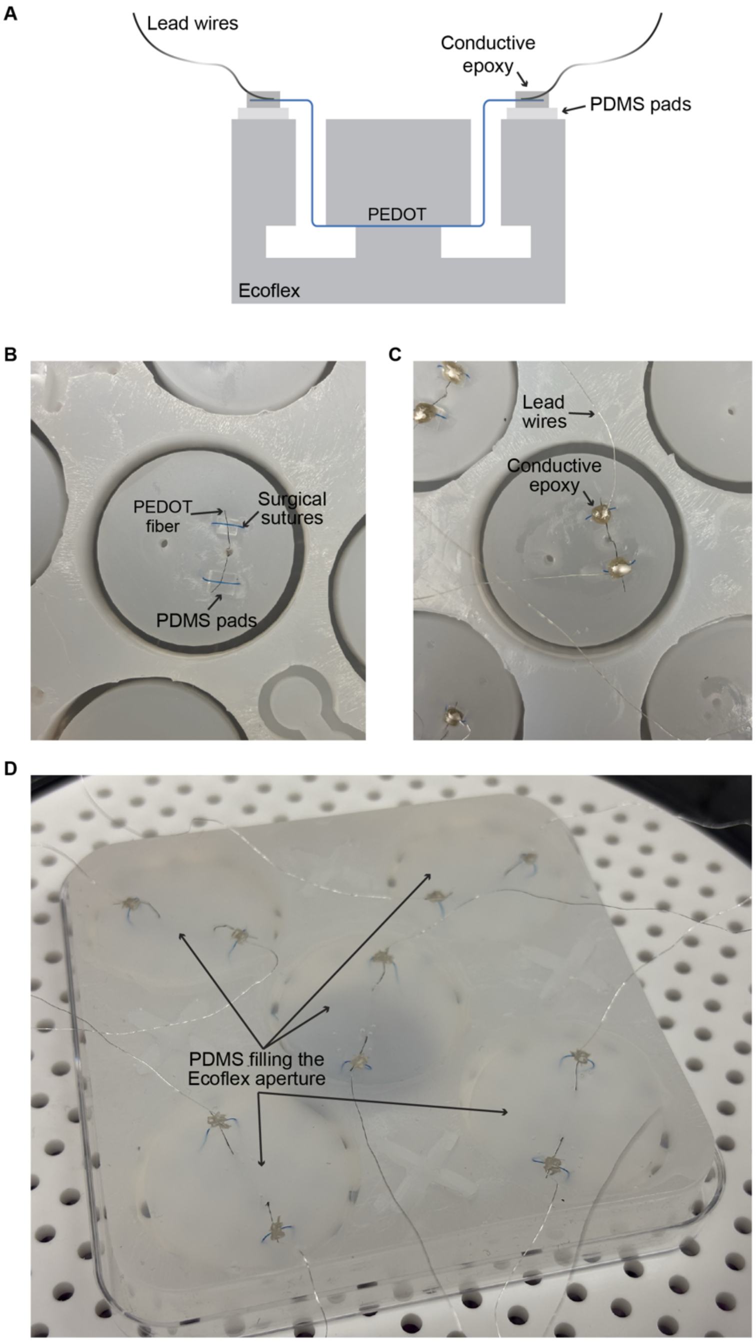


**Fig. S38.**

**Electrical interconnect architecture and encapsulation of fiber-to-wire interfaces.** (A) Cross-sectional schematics illustrating the electrical interfacing strategy used to transition from the integrated PEDOT fibers to external lead wires. (B-C) Photographic images depicting the physical implementation of the electrical interfacing, showing the alignment and bonding of the fibers to the lead wires. (D) Photographic image of the PDMS filling the Ecoflex mold apertures, resulting in molded PDMS devices. This step also serves as a robust 'potting' compound, protecting the electrical junctions from moisture ingress and mechanical delamination during handling and subsequent immersion in physiological media.

A

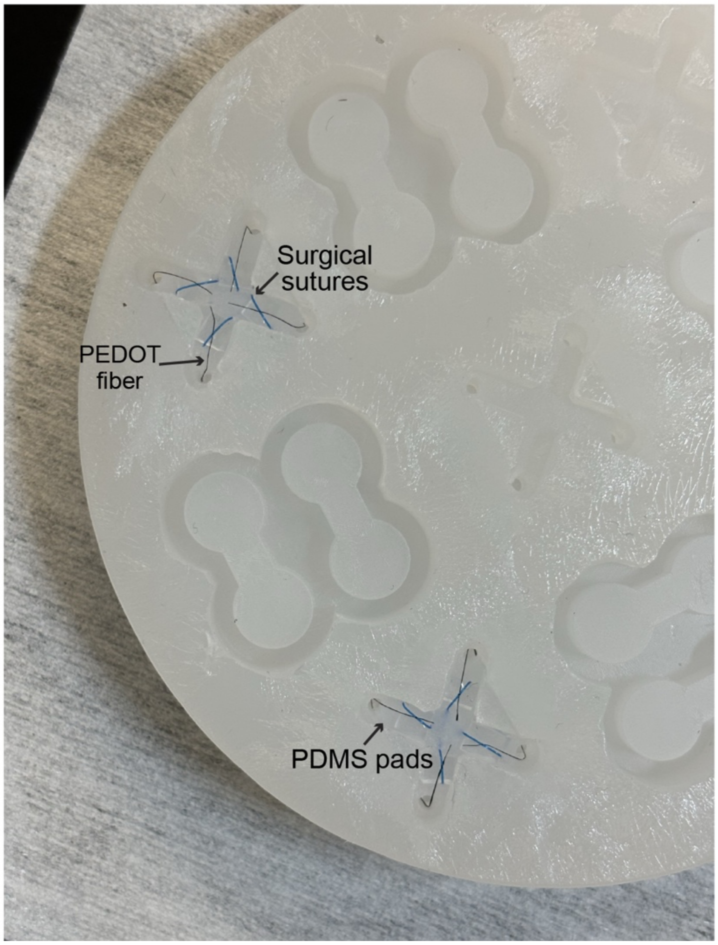


**Fig. S39.**

**Strategic positioning and securing of electrical interfaces within the biobot skeleton.** (A) Photographic image illustrating the protocol for securing the PEDOT fiber ends to facilitate stable electrical interfacing.

A

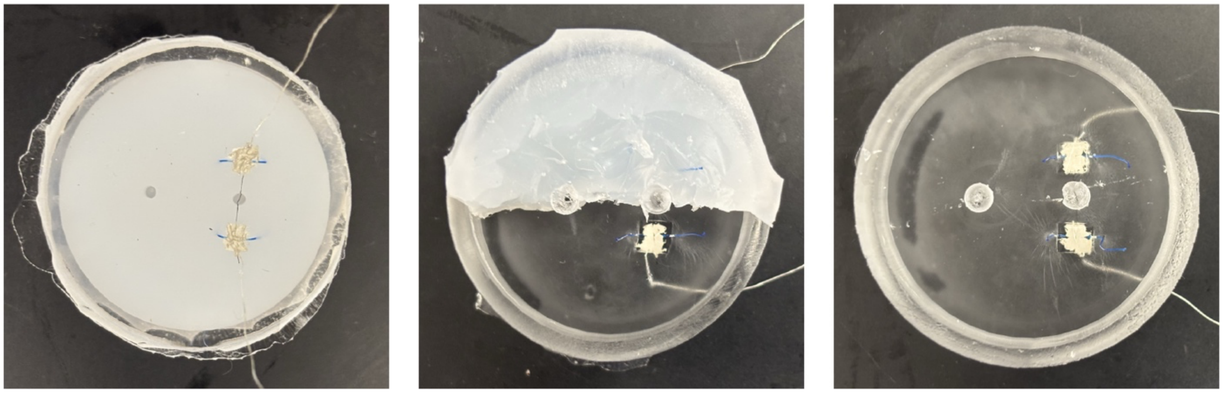

**Fig. S40.**

**Release of the molded PDMS architectures.** (A) Photographic image detailing the manual dissection protocol used to release the polymerized PDMS devices from the Ecoflex negative molds. The high flexibility and low surface energy of the parylene-coated Ecoflex mold, combined with strategic relief incisions, allow for the retrieval of the sensing (shown) and stimulation devices without inducing mechanical damage to the delicate PEDOT fiber interfaces or the elastomeric pillars. This final fabrication step ensures the structural fidelity and operational readiness of the biohybrid components prior to muscle tissue integration.

A

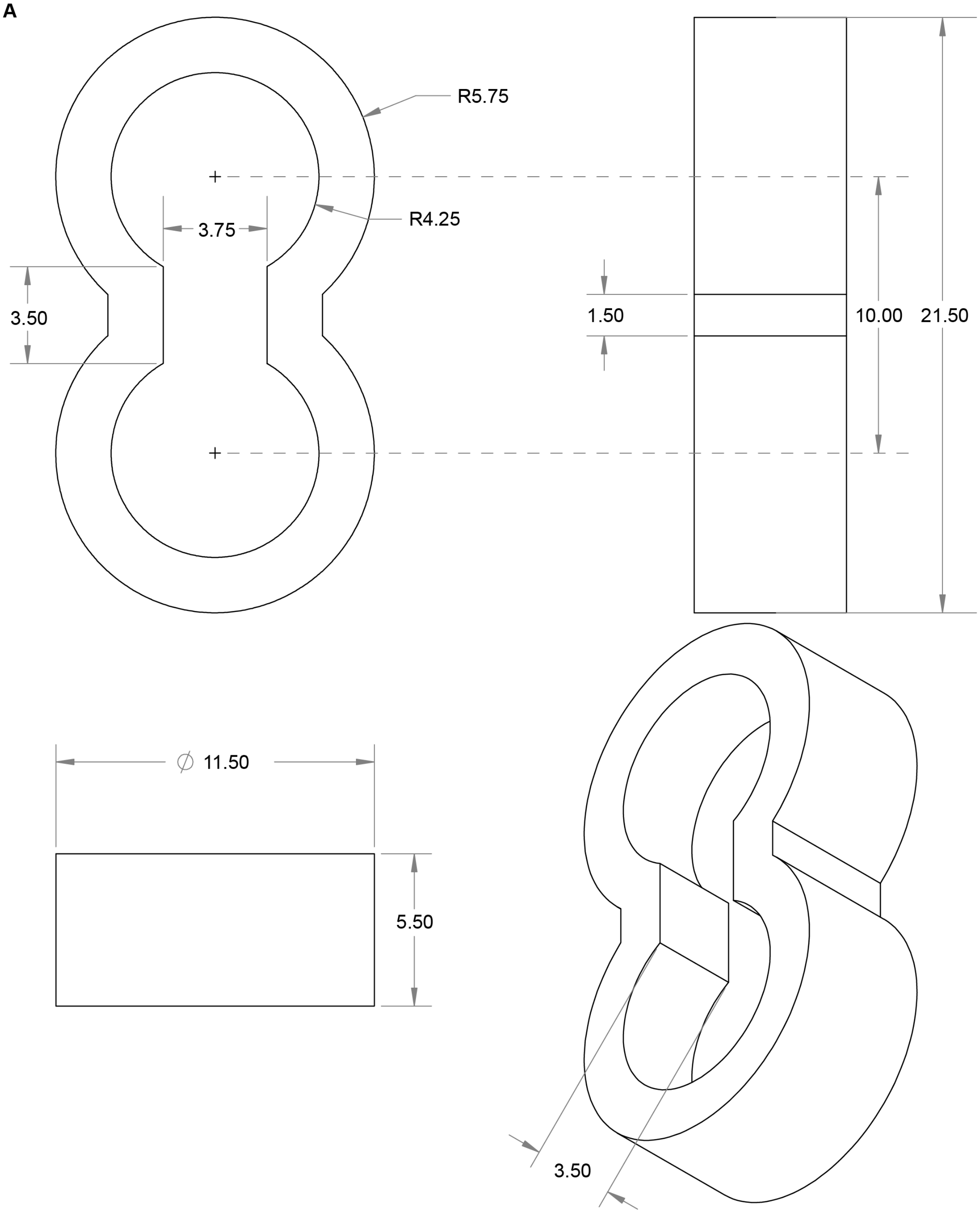


**Fig. S41.**

**CAD and geometric specifications of the figure-eight casting well.** (**A**) Detailed CAD schematics illustrating the structural dimensions (in mm) of the figure-eight casting well.

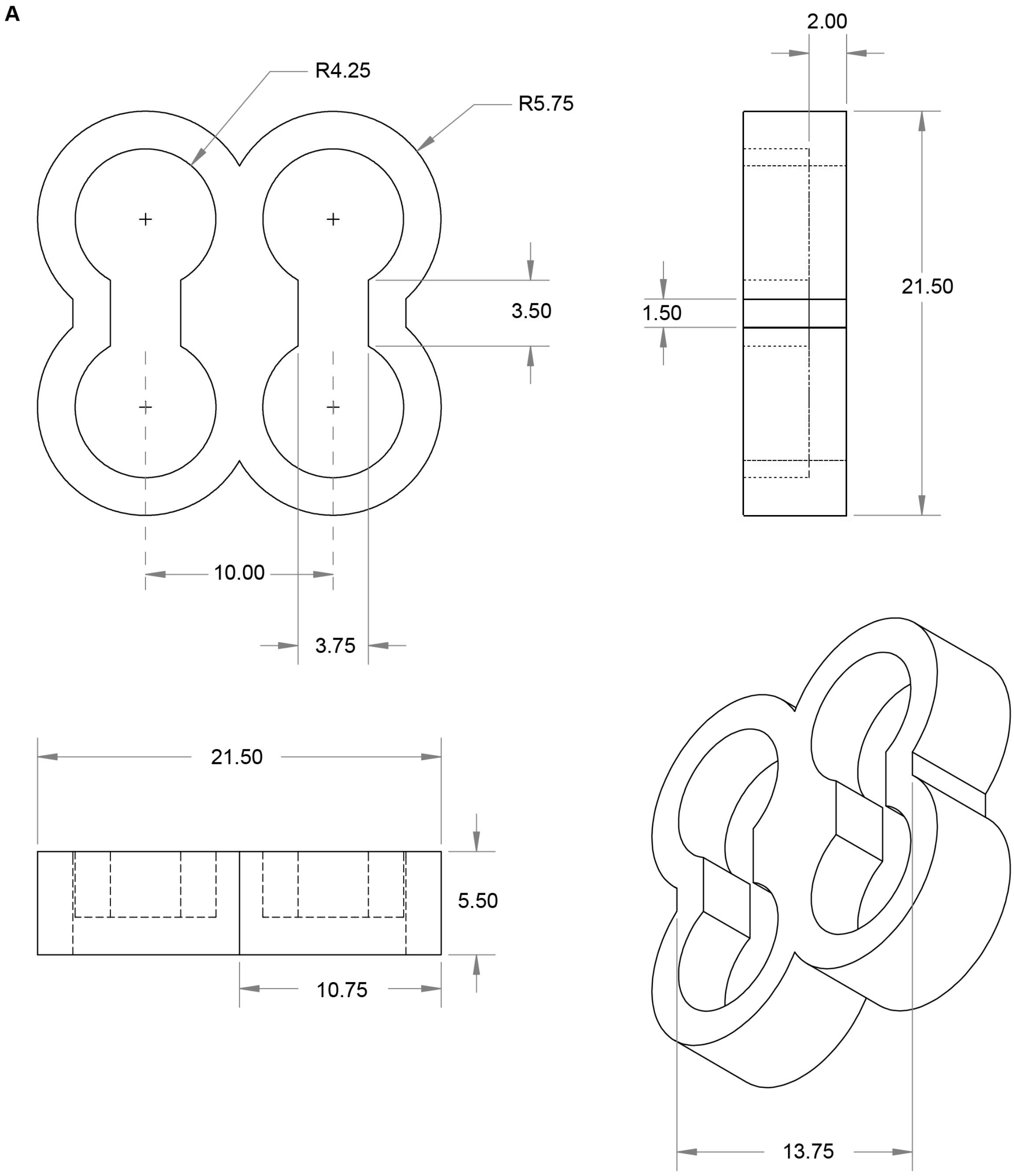


**Fig. S42.**

**CAD and geometric specifications of the dual figure-eight casting well.** (**A**) Detailed CAD schematics illustrating the structural dimensions (in mm) of the dual figure-eight casting well.

**A**

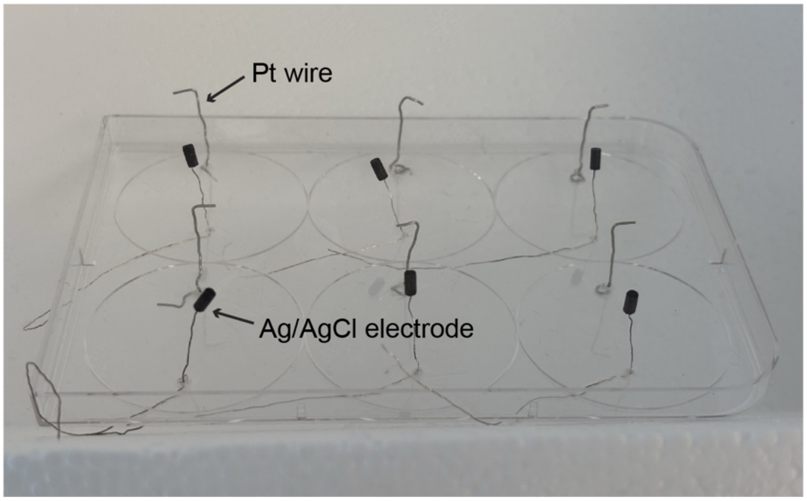


**Fig. S43.**

**Custom-engineered plate lid for sterilized electrical stimulation.** (**A**) Photographic image of the modified 6-well plate lid designed for stimulation across multiple experimental groups. Each well-position on the lid is integrated with a dedicated Pt wire and an Ag/AgCl reference electrode. This configuration ensures a stable and uniform electric field within each well, enabling the high-throughput maturation and functional testing of biohybrid actuators under identical electrochemical conditions. The integrated design minimizes contamination risk and evaporation while providing reliable electrical connectivity for long-term actuator studies.

A

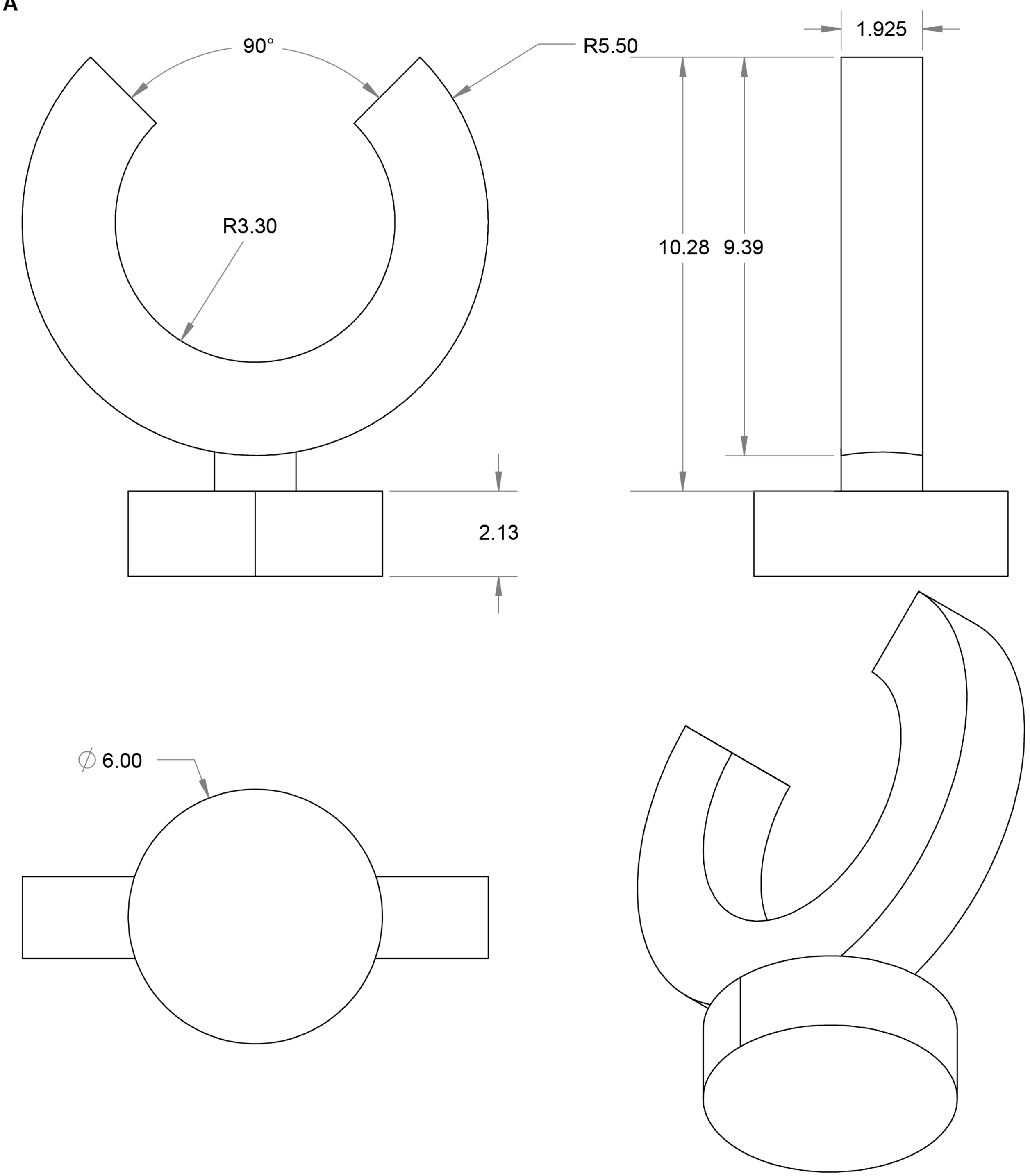


**Fig. S44.**

**CAD and geometric specifications of the biobot handle.** (**A**) Detailed CAD schematics illustrating the structural dimensions (in mm) of the dedicated biobot handle.

A

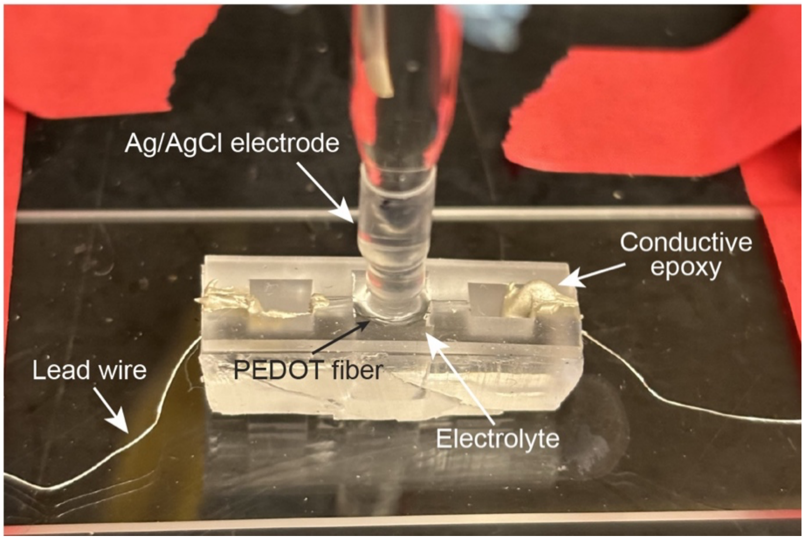


**Fig. S45.**

**Experimental configuration for electrochemical impedance spectroscopy (EIS).** (**A**) Photographic image of the experimental setup for EIS measurements, featuring the PEDOT fiber as the working electrode and an Ag/AgCl counter/reference electrode. Both electrodes are submerged in Phosphate-Buffered Saline (PBS) to simulate physiological electrochemical conditions.

A

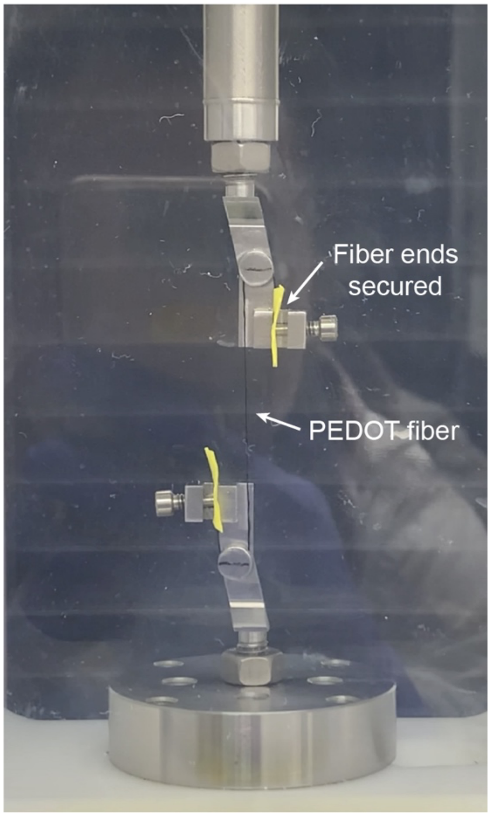


**Fig. S46.**

**Experimental configuration for in situ mechanical characterization.** (**A**) Photographic image of the experimental setup for the mechanical testing of PEDOT fibers. The fiber ends are securely anchored to a high-precision micro-tester and submerged in a Phosphate-Buffered Saline (PBS) bath maintained at a physiological temperature of 37 °C. This configuration allows for the real-time acquisition of stress-strain data under simulated operational conditions.

**Table S1. Randles equivalent circuit fitting parameters for PEDOT fibers across varying spinning speeds (N = 3).**

| | $R_s$ (Ω) | $SD_s$ (Ω) | $C_{dl}$ (μF) | $SD_{dl}$ (μF) | $R_{ct}$ (Ω) | $SD_{ct}$ (Ω) | $C_{dl}$% error | $R_s$% error | $R_{ct}$% error |
|---|---|---|---|---|---|---|---|---|---|
| 0.1 mL/min a | 548.07 | 3.90 | 612.96 | 8.19 | 132549.92 | 16306.86 | 1.34 | 0.71 | 12.30 |
| 0.1 mL/min b | 752.69 | 4.53 | 688.71 | 8.30 | 246633.40 | 54447.78 | 1.21 | 0.60 | 22.08 |
| 0.1 mL/min c | 680.60 | 4.09 | 616.64 | 7.16 | 267581.00 | 56412.08 | 1.16 | 0.60 | 21.08 |
| 0.2 mL/min a | 659.08 | 4.87 | 1661.06 | 28.34 | 82627.38 | 19116.76 | 1.71 | 0.74 | 23.14 |
| 0.2 mL/min b | 641.92 | 4.03 | 1037.63 | 13.61 | 153584.86 | 33467.80 | 1.31 | 0.63 | 21.79 |
| 0.2 mL/min c | 645.53 | 4.24 | 1546.09 | 22.87 | 107479.27 | 26160.60 | 1.48 | 0.66 | 24.34 |
| 0.3 mL/min a | 592.14 | 4.95 | 940.04 | 16.08 | 114729.69 | 22602.70 | 1.71 | 0.84 | 19.70 |
| 0.3 mL/min b | 619.89 | 5.06 | 769.19 | 12.48 | 156976.12 | 33307.64 | 1.62 | 0.82 | 21.22 |
| 0.3 mL/min c | 625.85 | 5.43 | 773.61 | 13.48 | 88885.05 | 11304.85 | 1.74 | 0.87 | 12.72 |

**Table S2. Comparative average power consumption (mean ± std mW) of PEDOT vs. Pt electrodes during active stimulation.**

| Voltage (V) | PEDOT (N = 3) | Pt (N=3) |
|---|---|---|
| 0.5 | 0.091 ± 0.006 | 0.052 ± 0.010 |
| 1 | 0.376 ± 0.034 | 0.295 ± 0.054 |
| 1.5 | 0.859 ± 0.090 | 0.816 ± 0.126 |
| 2 | 1.535 ± 0.174 | 1.535 ± 0.262 |
| 2.5 | 2.395 ± 0.284 | 2.783 ± 0.567 |
| 3 | 3.482 ± 0.459 | 4.475 ± 0.953 |
| 3.5 | 4.786 ± 0.662 | 6.555 ± 1.421 |

| | | |
|---|---|---|
| 4 | 6.303 ± 0.894 | 9.023 ± 1.977 |
| 4.5 | 8.020 ± 1.155 | 11.867 ± 2.624 |
| 5 | 9.964 ± 1.421 | 15.037 ± 3.379 |

**Table S3. Composition of C2C12 myoblast growth medium (GM).**

| Component | Volume (mL) | Concentration | Final Concentration |
|---|---|---|---|
| **DMEM** | 45 | - | ~86% (v/v) |
| **FBS** | 5 | - | ~10% (v/v) |
| **Pen/Strep** | 0.5 | 10000 U/mL | 100 U/mL |
| **L-glutamine** | 0.5 | 200 mM | 2 mM |
| **ACA** | 1 | 50 mg/mL | 1 mg/mL |

**Table S4. Formulation of C2C12 myoblast and hydrogel mixture for tissue casting.**

| Component | Volume (µL) | Concentration | Final Concentration |
|---|---|---|---|
| **C2C12** | 150 | $8.33*10^6$ cells/mL | $5*10^6$ cells/mL |
| **Matrigel** | 75 | - | ~30% (v/v) |
| **Fibrinogen** | 25 | 40 mg/mL | 4 mg/mL |
| **Thrombin** | 5 | 100 U/mL | 2 U/mL |

**Table S5. Composition of C2C12 myoblast differentiation medium (DM).**

| Component | Volume (mL) | Concentration | Final Concentration |
|---|---|---|---|
| **DMEM** | 450 | - | ~86% (v/v) |

| | | | |
|---|---|---|---|
| **Horse serum** | 50 | - | ~10% (v/v) |
| **Pen/Strep** | 5 | 10000 U/mL | 100 U/mL |
| **L-glutamine** | 5 | 200 mM | 2 mM |
| **ACA** | 10 | 50 mg/mL | 1 mg/mL |
| **IGF-1** | 25 μL | 1 mg/mL | 50 ng/mL |

### Table S6. List of parameters used in the Finite Element Analysis (FEA) simulations.

| | PDMS 10:1 | PDMS 20:1 | Parylene C |
|---|---|---|---|
| Young’s modulus | 1.4 [MPa] | 0.75 [MPa] | 2.8 [GPa] |
| Poisson’s ratio | 0.49 | 0.49 | 0.4 |
| Density | 970 [kg/m$^3$] | 970 [kg/m$^3$] | 1289 [kg/m$^3$] |

**Movie S1.**
**Simultaneous bilateral muscle activation.**

**Movie S1.**
**Selective activation of the left muscle bundle.**

**Movie S3.**
**Selective activation of the right muscle bundle.**

**Movie S4.**
**Biobot locomotion under alternating 1 Hz stimulation.**

**Movie S5.**
**Biobot locomotion under alternating 20 Hz stimulation.**

**Movie S6.**
**Unilateral biobot locomotion via left-muscle stimulation.**

**Movie S7.**
**Unilateral biobot locomotion via right-muscle stimulation.**
